# Methionine synthase reductase regulates heterochromatin independently of methionine synthesis through mitochondrial homeostasis

**DOI:** 10.64898/2025.12.19.695597

**Authors:** Fernanda Rezende Pabst, Andrey Tvardovskiy, Iratxe Estibariz, Veronica Finazzi, Pauline Couacault, Anna Artati, Michael Witting, Antonio Scialdone, Till Bartke, Robert Schneider, Daphne Selvaggia Cabianca

## Abstract

Metabolic enzymes can influence chromatin organization by modulating the availability of key metabolites, yet how specific metabolic reactions affect chromatin function remains poorly understood. Here, we show that in *Caenorhabditis elegans,* the methionine-cycle enzyme methionine synthase reductase (MTRR-l/MSR) regulates heterochromatin independently of methionine synthesis. Loss of MTRR-1, but not of the methionine synthase METR-l/MS, specifically reduces heterochromatic histone methylation, derepresses repetitive elements, and causes developmental delay. Multi-omics profiling revealed that *mtrr-1* mutants activate transcriptional programs associated with mitochondrial stress and accumulate long-chain acylcamitines, indicating disrupted mitochondrial homeostasis. Functional assays confirmed altered mitochondrial respiration in *mtrr-1* mutants, while genetic suppression of the PMK-3/MAPK mitochondrial retrograde signaling pathway partially restored repeat silencing. Consistently, direct perturbation of mitochondrial function was sufficient to induce heterochromatin derepression in wild type animals. Together, our results reveal a previously unrecognized mitochondria-to-chromatin axis controlled by the methionine-cycle enzyme MTRR-l/MSR, demonstrating that mitochondrial homeostasis is required for heterochromatin maintenance independently of canonical methionine metabolism.

## Introduction

Chromatin-modifying enzymes use metabolites to add or remove histone post-translational modifications (PTMs), revealing a tight connection between intracellular metabolism and chromatin regulation^1^. This raises the fundamental question of how metabolic state shapes chromatin regulation.

One of the most critical chromatin modifications is the methylation of histones and DNA, which depends on S-adenosylmethionine (SAM). Proper regulation of histone methylation marks is essential for all stages of development^2^, and their dysregulation has been linked to aging and cancer^3^, highlighting the need for precise control of this process. Several studies have shown that reducing intracellular SAM levels by depleting SAM synthetases alters histone^4^’^7^ and DNA^5^’^8^ methylation across species and cell types, underscoring the importance of controlling SAM production.

SAM is synthesized from the amino acid methionine^9^. While reductions in dietary methionine have been shown to alter histone methylation^6, 10–12^, the contribution of intracellular methionine synthesis to chromatin modification and regulation is unclear.

In mammals, methionine can be produced endogenously by methylating homocysteine using either 5-methyltetrahydrofolate or betaine as methyl donors. The first reaction requires the enzymes Methionine Synthase (MS) and Methionine Synthase Reductase (MSR), linking the methionine cycle to the folate cycle^13^’^14^. The second pathway is centered on Betaine-Homocysteine Methyltransferase (BHMT).

Despite the central role of methionine metabolism in methylation reactions, studies examining how intracellular methionine synthesis influences DNA or histone methylation in non-cancer cells remain sparse. While deletion of Bhmt alters DNA CpG methylation in mice^15^, loss-of-function studies of MSR have reported divergent effects on CpG DNA methylation^16^’^17^. Strikingly, however, the roles of MS and MSR in regulating histone PTMs have not been comprehensively or unbiasedly investigated, leaving a critical and fundamental gap in our understanding of how intracellular methionine synthesis contributes to epigenetic processes. Here, we use the nematode *Caenorhabditis elegans,* which (i) relies exclusively on the MS/MSR pathway to produce methionine^18^ and (ii) lacks CpG DNA methylation^19^, to investigate the role of endogenous methionine synthesis in regulating histone methylation, thereby avoiding indirect effects from secondary methionine synthesis pathways and confounding feedback from or to DNA methylation.

## Results

### Loss of MTRR-l/MSR, but not METR-l/MS, causes a pronounced growth defect and selectively reduces heterochromatic histone methylation

In C. *elegans,* methionine can be directly absorbed from the diet or synthesized intracellularly via the enzymes METR-1 (MS) and MTRR-1 (MSR). METR-l/MS catalyzes the methylation of homocysteine to generate methionine, using the methyl group donated by 5-methyl-tetrahydrofolate and vitamin B12 as cofactor (Fig. la). After several rounds of catalytic cycles, the cobalt atom of the METR-1/MS-bound vitamin B12 becomes oxidized, inactivating the enzyme^20^. MTRR-l/MSR is a di-flavin oxidoreductase^21^ required to reduce the oxidized vitamin B12 and restore METR-l/MS functionality^22^ (Fig. la).

To explore the role of endogenous methionine synthesis in chromatin regulation, we generated *metr-1* and *mtrr-1* knockout animals using CRISPR/Cas9-mediated deletion of their entire open reading frames. Although all mutants reached adulthood and remained fertile, we observed differences in their growth. Measurements of body size 72 hours after the onset of the first larval stage (LI) on a standard *E. coli* OP50 diet revealed that *metr-1* mutants were smaller than wild-type animals. Interestingly, loss of *mtrr-1* caused an even more pronounced growth defect (Fig. lb) and delayed adulthood by approximately one day (data not shown), suggesting that MTRR-1 has a function independent of methionine synthesis.

To investigate the roles of METR-1 and MTRR-1 in regulating histone H3 methylation, we performed unbiased mass spectrometry on *metr-1* and *mtrr-1* mutants and wild-type controls at the LI stage, thereby avoiding variation in developmental stage between genotypes. Loss of either MTRR-1 or METR-1 altered the global abundance of specific histone H3 lysine (K) methylation marks. Specifically, trimethylation on H3K27 (H3K27me3), H3K9me2, and H3K23me3 were reduced in both mutants, whereas H3K36 methylation remained unchanged (Fig. lc). Notably, the specific effects on histone methylation did not correlate with the relative abundance of the individual histone methylation marks^23^. Strikingly, however, the histone methylation profiles of *metr-1* and *mtrr-1* mutants only partially overlapped, with the effects of the *mtrr-1* mutation generally being stronger (Fig. lc and SI a).

Chromatin exists in two major, functionally distinct forms: transcriptionally active, open euchromatin and transcriptionally repressed, compact heterochromatin, each characterized by specific histone methylation marks that both reflect and contribute to its functional state^24^. Across species, typical euchromatic marks include H3K4me3 and H3K36me3, whereas canonical heterochromatic marks are H3K9me2/me3 and H3K27me3. We found that *metr-1* deletion significantly reduced the levels of both euchromatic (H3K4me2 and H3K4me3) and heterochromatic (H3K9me2 and H3K27me3) methylation marks. By contrast, loss of MTRR-1 caused a specific reduction in heterochromatic histone methylation, affecting both canonical (H3K9me2 and H3K27me3) and less well-characterized, heterochromatin-associated^25–28^ (H3K23me2 and H3K23me3) marks.

The mild effect of METR-1 loss on histone methylation is consistent with previous reports indicating that methionine is an essential dietary amino acid in C. *elegans*^19^ and humans^30^. Feeding wild-type animals a low-methionine diet (using *E. coli metE* mutants) did not affect growth compared with animals fed *motB E. coli,* a reduced-motility bacterial strain used as a control (Fig. Sib). In contrast, *metr-1* mutants under the same low-methionine conditions exhibited a further reduction in body size (Fig. Sib), indicating that animals lacking METR-1 cannot compensate for low dietary methionine.

In sum, loss of the methionine synthase MTRR-1 causes a growth delay and a pronounced reduction in histone methylation that is specific to heterochromatin marks, pointing to a distinct, METR-1-independent function of MTRR-1.

### Loss of MTRR-1, but not of METR-1, impairs heterochromatin silencing

Across species, H3K9 methylation is a hallmark of constitutive heterochromatin and plays a key role in silencing repetitive elements^31^. H3K23me2 and H3K23me3 are less well understood, evolutionarily conserved marks associated with heterochromatin^23^’^25–28^. In C. *elegans,* H3K23me2 co-occupies the histone tail with H3K9me3^25^, and H3K9me2 levels positively correlate with H3K23me3^23^. To assess whether the global reduction in heterochromatic marks observed in *mtrr-1* mutants translates into a functional heterochromatin defect, we performed three different experiments. We performed RNA-seq to measure the expression of repetitive elements in *mtrr-1-* and *metr-1-* null animals. Using TElocal^32^, we found that *mtrr-1* mutants exhibit altered expression of 74 repetitive elements (Fig. 2a), of which 74% were upregulated, consistent with a loss of silencing. In contrast, worms lacking METR-1 displayed only 2 dilferentially expressed repeats (Fig. 2b), indicating that heterochromatin silencing is largely unperturbed in absence of the methionine synthase. The same result was obtained using an independent loss-of-function *metr-1* allele *(metr-1 (ok521))* (Fig. S2a), previously used to study methionine synthesis^33^’^34^. Because this deletion does not encompass the entire ORF and may produce a truncated protein, we used the CRISPR/Cas9 *metr-1* null allele generated in this study for all further analyses, unless otherwise indicated.

**Figure 1.**
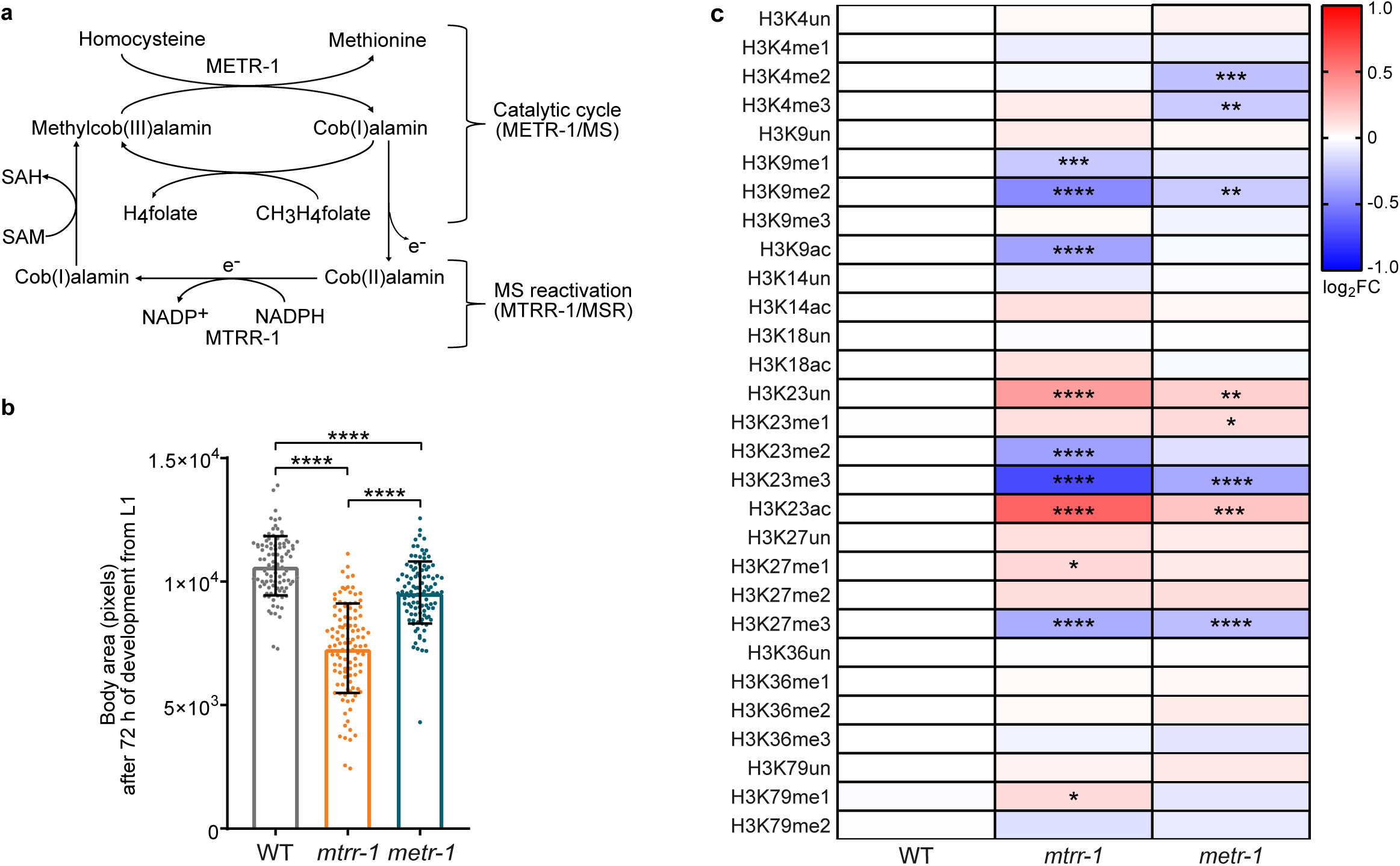
***mtrr-1* mutants show stronger growth defects than *metr-1* mutants and a specific reduction in heterochromatic histone methylation. a.** Schematic representation of the methionine cycle adapted from20, illustrating the catalytic reaction carried out by METR-1/MS and the reactivation step mediated by MTRR-1/MSR. **b**. Quantification of the body area (pixels) of *mtrr-1* and *metr-1* mutants grown on NGM plates with OP50 bacteria and measured after 72 h from the onset of the L1 stage. Statistical significance was assessed using a one-way ANOVA with Games-Howell’s multiple comparisons test. Data represent the mean of three independent biological replica, each including at least 20 animals. ****p < 0.0001. **c**. Heatmap of the relative log₂ fold change (log₂ FC) in the indicated histone H3 modifications in *mtrr-1* and *metr-1* mutant L1s, relative to WT worms. Statistical significance was determined using a two-way ANOVA with Dunnett’s multiple comparison test. Data represent the mean of four independent biological replicates. *p<0.05, **p<0.01, ***p<0.001, ****p<0.0001. For (b, c) exact p and n values are in Supplementary Table 2.

**Figure 2.**
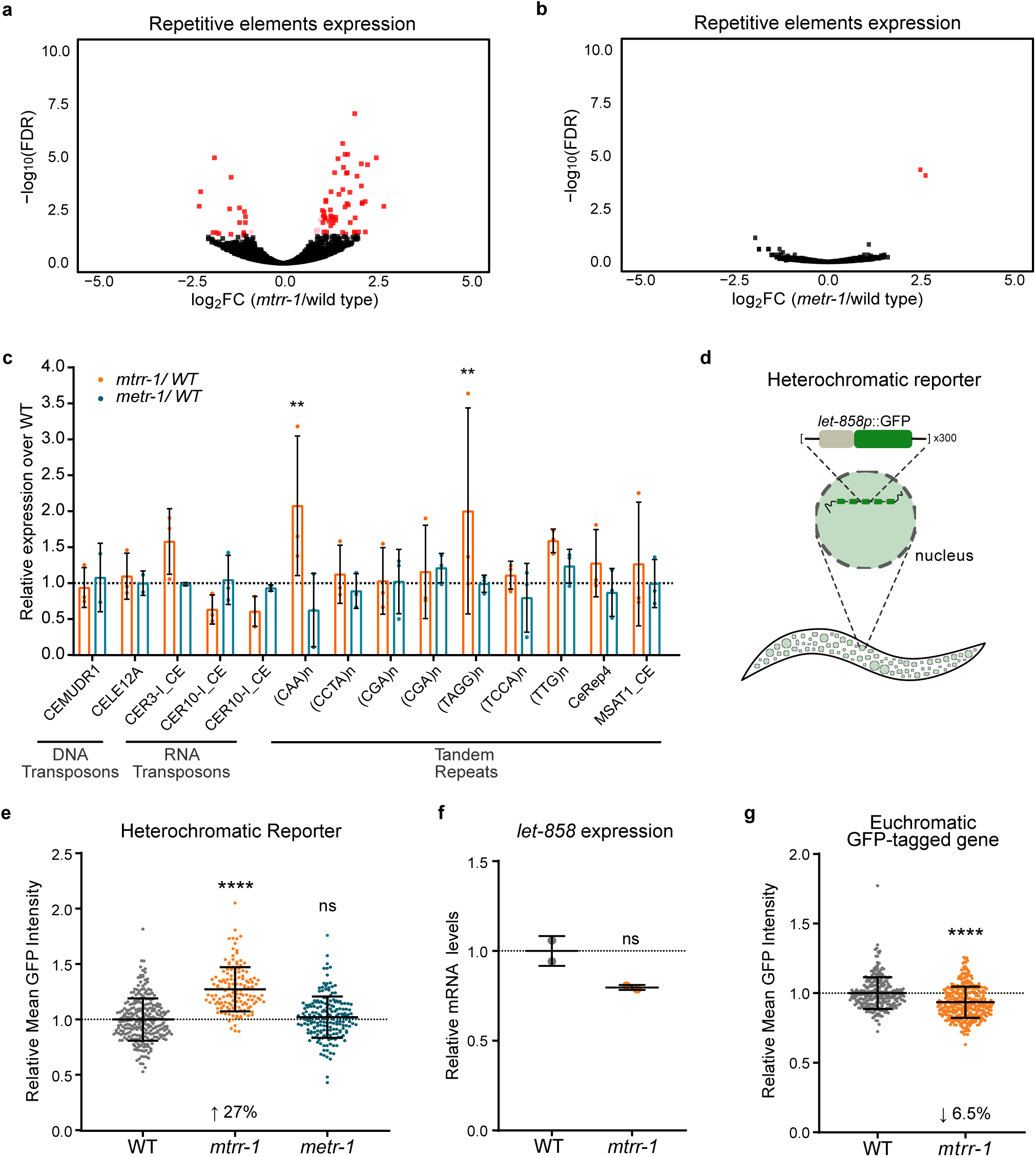
Repetitive elements are derepressed in *mtrr-1* mutants. **a.** Volcano plot of repetitive elementś expression in *mtrr-1* mutants compared to wild type (WT). Significant up- and down-regulated transcripts are highlighted (in red FDR <0.05 and log_2_FC ≥│1│, in pink FDR <0.05 and log_2_FC <│1│, data represent the mean of two independent biological replicates). **b**. As in (a), but for *metr-1* mutants compared to wild type (WT). Data represent the mean of three independent biological replicates. **c**. Barplot of the expression of the indicated repetitive elements measured by qPCR, comparing the expression of *mtrr-1* and *metr-1* mutants relative to wild type (WT). Statistical significance was determined by two-way ANOVA with Dunnett’s Multiple Comparison test. **p<0.01. Data represent the mean of at least two independent biological replicates. **d**. Schematic representation of the heterochromatic reporter *pkIs1582[let-858::GFP;rol-6(su1006)]*4 **e**. Beeswarm plot of the quantification of the mean GFP intensity expressed from the heterochromatic reporter (d) in L1s of *mtrr-1* and *metr-1* mutants relative to wild type (WT). Statistical significance was determined by Kruskal-Wallis test. Data are from three independent biological replicates, each with at least 20 animals. ****p<0.0001, ns = not significant. **f**. Quantification of relative mRNA levels of *let-858* in *mtrr-1* mutant L1 larvae relative to wild type (WT). Statistical significance was with Student’s two-tailed t-test. Data are from two independent biological replicates. ns = not significant. **g**. Beeswarm plot of the mean GFP intensity of MRG-1-GFP (a euchromatic, active single copy transgene) in *mtrr*-1 mutants relative to wild type (WT). Statistical significance was determined with Mann-Whitney test. Data are from four independent biological replicates, each with at least 45 animals. ns = not significant. For (c, e, f, g) exact p and n values are in Supplementary Table 2.

H3K9 methylation—particularly H3K9me2, whose levels are reduced in *mtrr-1* mutants—has been shown to repress specific repetitive elements in C. *elegans*^35^. We performed RT-qPCR to measure the expression of known H3K9me2/3-repressed elements in *metr-1* and *mtrr-1* mutants. Consistent with our RNA-seq results, only deletion of *mtrr-1,* and not *metr-1,* led to derepression of repetitive elements (Fig. 2c).

We next used a GFP-based heterochromatic reporter (Fig. 2d), previously used to identify factors regulating heterochromatin silencing^4^ as a third means to assess the effects of *mtrr-1* and *metr-1* mutations on heterochromatin silencing, by live imaging. Consistent with our previous results, only *mtrr-1* mutants, but not *metr-1,* showed derepression of the reporter (Fig. 2e). This effect is specific to the heterochromatic nature of the reporter: (i) it is not mediated by the reporter’s promoter, as *let-858* is not derepressed in its endogenous context (Fig. 2f), and (ii) loss of MTRR-1 does not increase GFP expression when this is driven from a euchromatic, actively expressed allele (Fig. 2g). Additionally, we found that derepression of the heterochromatic reporter in *mtrr-1* mutants could not be rescued by knock down of genes in: (i) folate metabolism *(dao-3* and *K07E3.4),* (ii) transsulfuration pathway *(cbs-1)* or (iii) the propionate response *(nhr-10)* (Fig. S2b). These results indicate that neither accumulation of folate-cycle intermediates, overactivation of the transsulfuration pathway, nor enhanced production of acetyl-CoA from propionate alone drives the reduced heterochromatin silencing in *mtrr-1* mutants. Because these pathways are expected to be perturbed in both *metr-1* and *mtrr-1* mutants^36^, their inability to rescue supports the conclusion that MTRR-1 regulates heterochromatin independently of methionine synthesis.

### Gene expression is more strongly altered upon *mtrr-1* deletion compared with *metr-1,* and is characterized by the activation of stress response pathways

To further investigate METR-1-independent functions of MTRR-1, we quantified gene expression by RNA-seq in *mtrr-1* and *metr-1* mutants compared with wild-type animals. Deletion of *mtrr-1* altered the expression of 655 genes (484 upregulated and 171 downregulated; Fig. 3a), whereas *metr-1* mutants showed minimal changes (19 upregulated and 23 downregulated; Fig. 3b). Approximately 40% of the genes altered in *metr-1* mutants were similarly affected in *mtrr-1* mutants (42% of upregulated and 39% of downregulated genes; Fig. 3c), consistent with their shared biochemical relationship. Similar results were obtained using the independent loss-of-function allele *metr-1 (ok521)* (Fig. S3a,b), with ∼70% of differentially expressed genes in the CRISPR-derived *metr-1* knockout overlapping with those in *metr-l(ok521)* (Fig. S3c). Remarkably, however, most differentially expressed genes in *mtrr-1* mutants were unique to this background, indicating that MTRR-1 influences gene expression independently of its role in methionine synthesis—consistent with the phenotypes described above.

**Figure 3.**
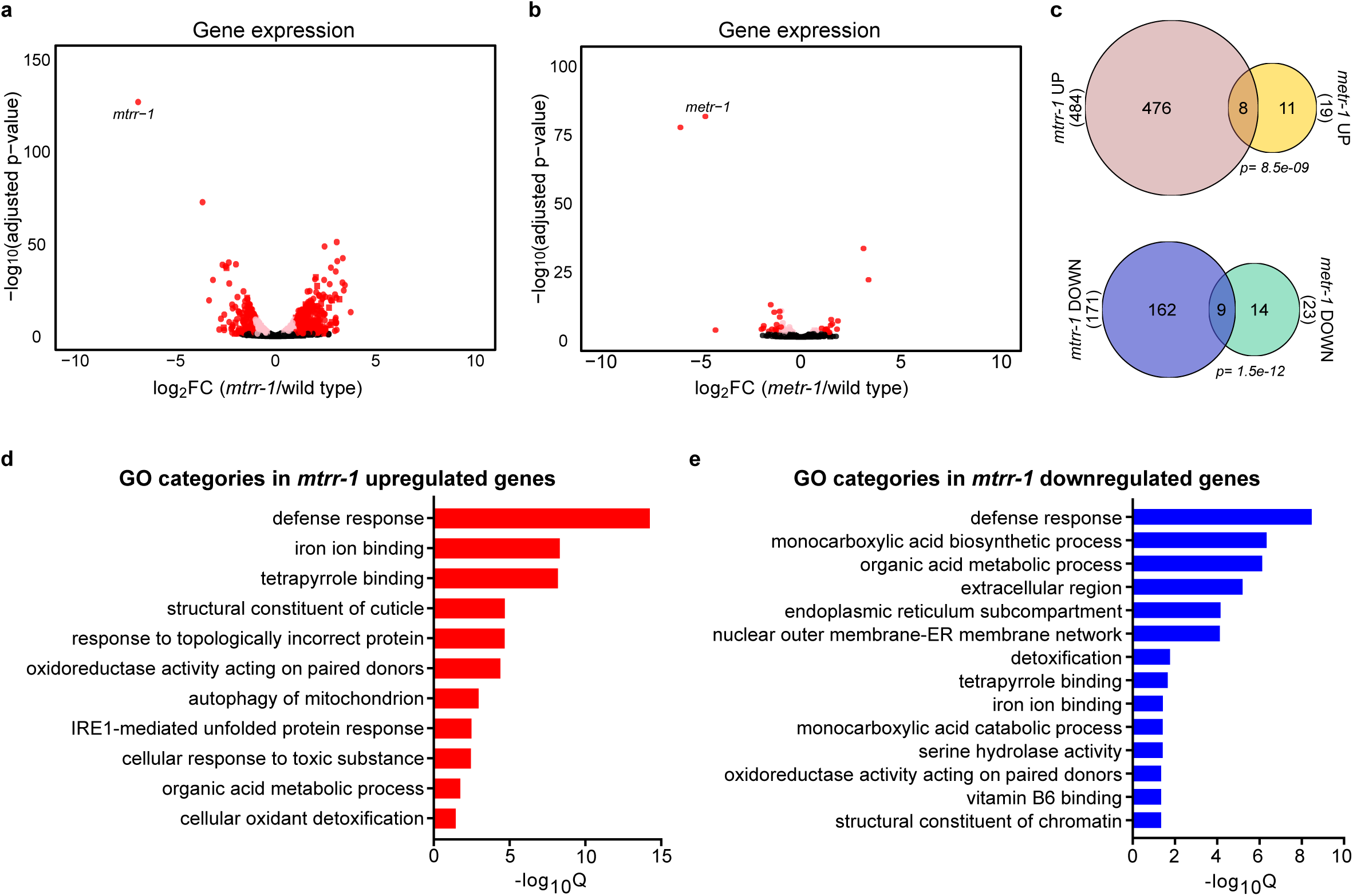
***mtrr-1* regulates gene expression involved in mitochondrial and ER stress responses independently of methionine synthesis. a**. Volcano plot of gene expression in *mtrr-1* mutants compared to wild type (WT). Differential expression analysis was calculated with DESeq2. Significant up- and down-regulated transcripts are highlighted (in red FDR <0.05 and log_2_FC ≥│1│, in pink FDR <0.05 and log_2_FC <│1│, data represent the mean of two independent biological replicates). **b**. as in (a), but in *metr-1* mutants compared wild type (WT). Data represent the mean of three independent biological replicates. **c**. Venn diagrams showing the overlap between upregulated (top) and downregulated (bottom) genes in *mtrr-1* and *metr-1* mutants. Significance of the overlap was determined with a hypergeometric test. **d**. GO terms significantly enriched among upregulated genes in *mtrr-1* mutants. **e**. GO terms significantly enriched among downregulated genes in *mtrr-1* mutants. GO terms were determined with the Enrichment Analysis tool of Wormbase.

Gene Ontology (GO) enrichment analysis revealed that *mtrr-1* mutants exhibit altered regulation of stress-response pathways (Fig. 3d,e). Upregulated genes were enriched for categories such as “defense response” and “response to biotic stimulus”, which are known to be activated during mitochondrial stress^37^’^38^. Categories associated with Endoplasmic Reticulum (ER) stress, such as “responses to topologically incorrect protein” and “unfolded protein response” (UPR), were enriched as well. Correspondingly, we observed upregulation of ER and cytosolic UPR markers *hsp-4* and *Hsp70* genes *(F44E5.4* and *F44E5.5), hsp-16.2, hsp-16.41*^39^*∼*\* respectively. In addition, we observed strong upregulation of *tbb-6,* a marker of the PMK-3/p38 MAPK mitochondrial stress signaling pathway^42^.

Among downregulated genes, in addition to categories such as “defense response”, “ER subcompartment” and “interaction between ER and nucleus”, we observed enrichment for “monocarboxylic acid biosynthetic process” and “monocarboxylic acid catabolic process”, suggesting perturbations in lipid metabolism in *mtrr-1* mutants.

### MTRR-1 deficiency, but not METR-1 loss, causes long-chain acylcarnitine accumulation and alters mitochondrial functionality

To assess metabolic changes in *mtrr-1* and *metr-1* mutants and identify signatures unique to loss of the methionine synthase reductase MTRR-1, we performed untargeted metabolomic profiling. We found that levels of SAM and its byproduct S-adenosylhomocysteine (SAH) were unchanged in both *mtrr-1* and *metr-1* animals (Fig. 4a), consistent with the fact that these mutants obtain methionine primarily from their diet and thus maintain SAM production. Additionally, we detected no differences in the SAM/SAH ratio (Fig. 4b). Because SAH is a potent inhibitor of methyltransferases, including histone methyltransferases, an unchanged SAM/SAH ratio supports the conclusion that global methylation capacity is unaffected in these mutants. Accordingly, knockdown of the SAM synthetase *sams-4* derepressed the heterochromatic reporter additively with *mtrr-1* deletion (Fig. 4c), indicating that *mtrr-1* mutants remain sensitive to reduced SAM levels.

**Figure 4.**
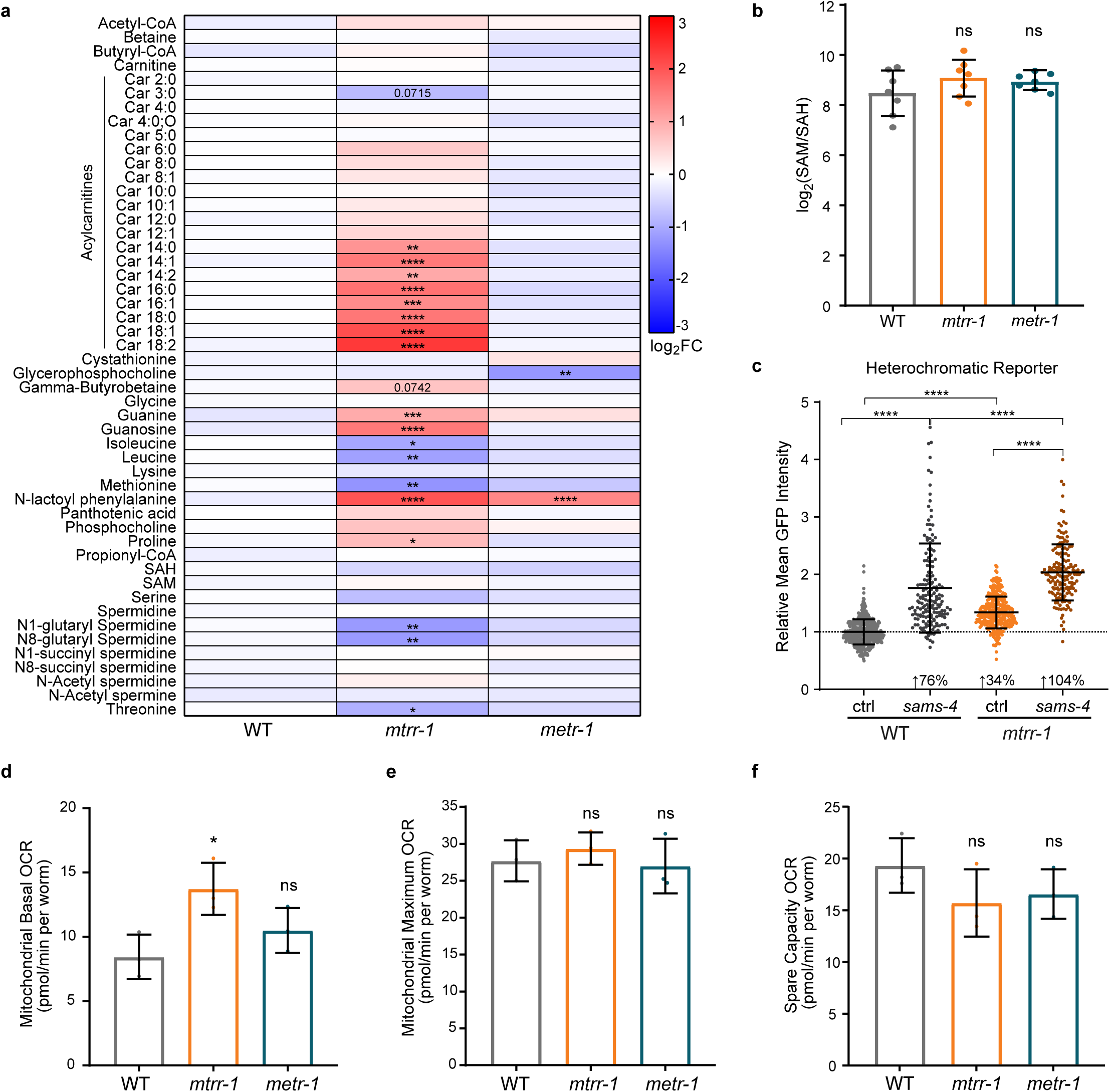
Accumulation of long-chain acylcarnitines and altered mitochondrial respiration are specific to *mtrr-1* mutants. **a.** Heatmap of the relative log₂ fold change (log₂ FC) in the indicated metabolites in *mtrr-1* and *metr-1* mutant L1s relative to wild type (WT). Statistical significance was determined by two-way ANOVA with Dunnett’s Multiple Comparison test. Data represent the mean of seven independent biological replicates. *p<0.05, **p<0.01, ***p<0.001, ****p<0.0001. **b**. Barplot of the log_2_ SAM/SAH ratio in wild type (WT), *mtrr-1* and *metr-1* mutant L1s. Statistical significance was determined by one-way ANOVA with Dunnett’s Multiple Comparison test. Data represent the mean of seven independent biological replicates. ns = not significant. **c**. Beeswarm plot of the quantification of the mean GFP intensity expressed from the heterochromatic reporter outlined in Fig. 2d, under the indicated RNAi conditions, relative to wild type (WT) control (ctrl) RNAi L4440. Statistical significance was determined by Kruskal-Wallis test. Data are from three independent biological replicates, each with more than 20 animals. ****p<0.0001. **d**. Mitochondrial basal Oxygen Consumption Rate (OCR) of day 2 adults in wild type (WT), *mtrr-1* and *metr-1* mutants. Statistical significance was determined by one-way ANOVA with Dunnett’s Multiple Comparison test. Data represent the mean of three independent biological replicates. *p<0.05, ns = not significant. **e**. Mitochondrial maximum OCR of day 2 adults in wild type (WT), *mtrr-1* and *metr-1* mutants. Statistical significance was determined by one-way ANOVA with Dunnett’s Multiple Comparison test. Data represent the mean of three independent biological replicates. ns = not significant. **f**. Spare Capacity OCR of day 2 adults in wild type (WT), *mtrr-1* and *metr-1* mutants. Statistical significance was determined by one-way ANOVA with Dunnett’s Multiple Comparison test. Data represent the mean of three independent biological replicates. ns = not significant. Exact p and n values are in Supplementary Table 2.

We next analyzed additional metabolites. In line with our previous data and earlier studies showing that the main source of methionine is diet, we found that methionine levels were unaffected in *metr-1* mutants (Fig. 4a). In contrast, methionine levels are slightly decreased in *mtrr-1* mutants (Fig. 4a). However, global SAM levels remained unchanged, indicating that methyl donor availability is maintained. This observation suggests that methionine is utilized differently in *mtrr-1* mutants rather than being limiting for SAM production.

Among all metabolites profiled (Figs. 4a and S4a), the most striking »//rr-7-specific change was the marked accumulation of long-chain acylcamitines. These molecules form when fatty acyl-CoAs are conjugated to carnitine by carnitine palmitoyltransferase I at the outer mitochondrial membrane to enable their transport into the mitochondrial matrix for P-oxidation^43^. Inside mitochondria, these fatty acids are oxidized to generate energy and acetyl-CoA, and their accumulation is typically associated with impaired P-oxidation^44^. Despite this, acetyl-CoA and other short-chain acyl-CoAs, including propionyl- and butyryl-CoA, remained unchanged in *mtrr-1* mutants (Fig. 4a). Consistent with a compensatory response, we observed a specific reduction in isoleucine, leucine, and threonine—amino acids that can feed the TCA cycle to support acetyl-CoA production when P-oxidation is compromised^45^.

Another mtrr-7-specific metabolic alteration is the selective reduction of Nl-glutaryl-spermidine and N8-glutaryl-spermidine, while other spermidine derivatives remain unchanged (Fig. 4a). In *C. elegans,* these glutarylated spermidines—and not other spermidine derivatives—are specifically regulated by the mitochondria-localized sirtuin SIR-2.3^46^, further supporting altered mitochondrial physiology in *mtrr-1* mutants.

Together with the activation of transcriptional programs associated with mitochondrial and endoplasmic reticulum (ER) stress (Fig. 3d, e), these results indicate that an impaired mitochondrial function is specific to *mtrr-1* mutants.

To directly assess mitochondrial activity, we performed Seahorse assays in *mtrr-1* and *metr-1* mutants. Mitochondrial basal oxygen consumption rate (OCR) was elevated only in *mtrr-1* mutants (Fig. 4d), whereas maximal OCR (Fig. 4e) and spare respiratory capacity (Fig. 4f) were unchanged in both genotypes. Altogether, these results confirm that mitochondrial function is specifically altered in *mtrr-1* mutants.

### Neither antioxidant treatment nor *metr-1* deletion restores repetitive element silencing in *mtrr-1* mutants

Because mitochondrial dysfunction is frequently associated with oxidative stress, and MTRR-1 functions as an oxidoreductase, we asked whether oxidative stress contributes to repeat derepression in *mtrr-1* mutants. To test this possibility, we first supplemented *mtrr-1* animals with reduced glutathione (GSH) or its precursor N-acetyl-L-cysteine (NAC), which increase intracellular glutathione availability and antioxidant capacity, and quantified by RT-qPCR the expression of four repetitive elements derepressed upon *mtrr-\* loss. Supplementation with either GSH or NAC failed to reduce repeat expression (Fig. 5a), indicating that increasing glutathione availability is not sufficient to restore repeat silencing.

**Figure 5.**
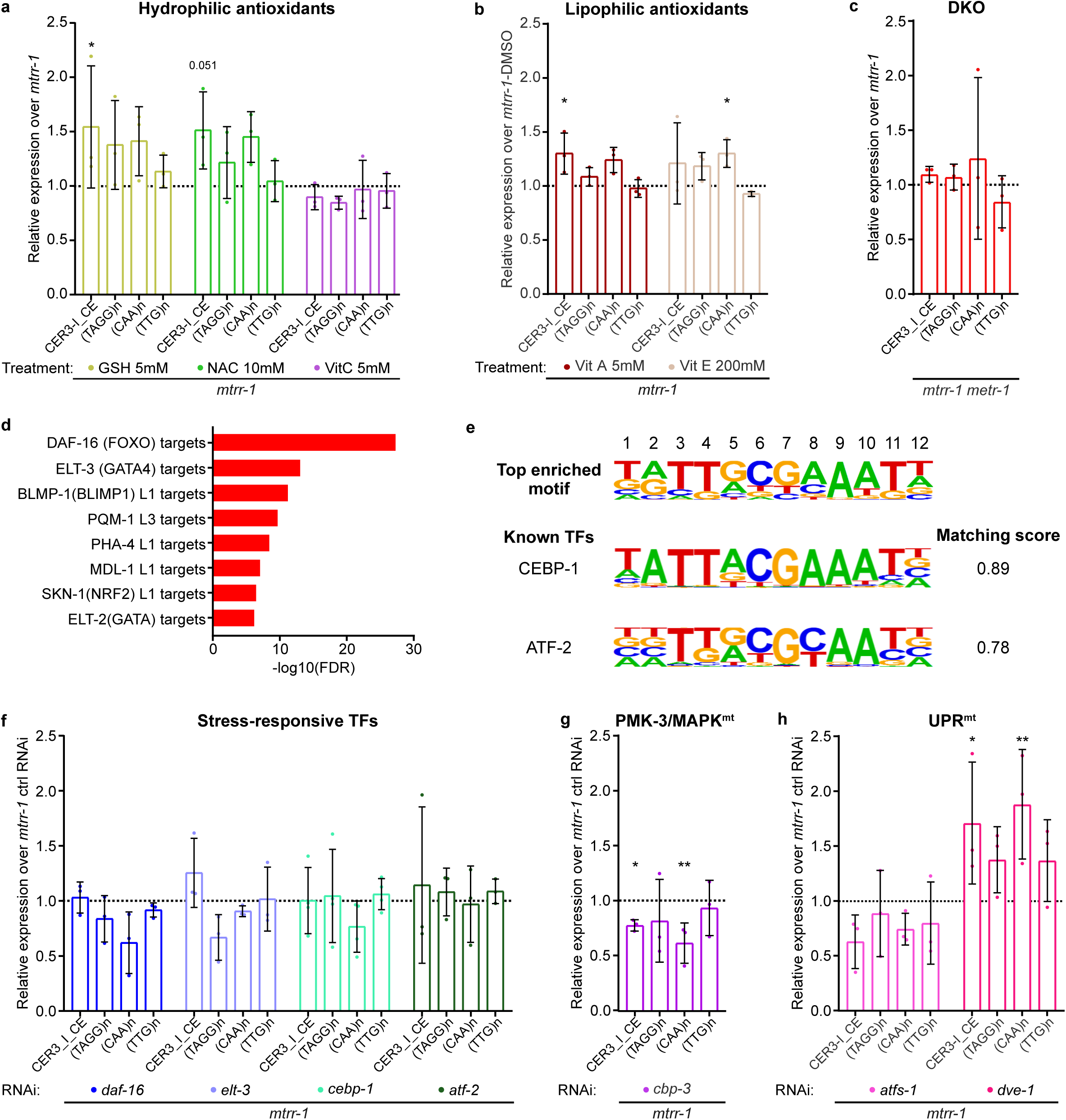
Knock down of the mitochondrial stress responsive factor CBP-3 partially rescues repeats derepression in *mtrr-1* mutants. **a.** Barplot of the expression of the indicated repetitive elements, measured by qPCR, in *mtrr-1* mutants treated with the indicated antioxidant and relative to untreated *mtrr-1*, as control (ctrl). **b.** As in (a) but relative to DMSO-treated *mtrr-1*. **c.** As in (a) but in *mtrr-1 metr-1* double mutants relative to single *mtrr-1* mutants. **d.** Enrichment analysis of upregulated genes in *mtrr-1* mutants among transcription factor (TF) target gene sets. The analysis was performed using WormExp50. **e.** The two highest-scoring TF motif matches for the top *de novo* motif identified in the promoter regions of genes upregulated in *mtrr-1* mutants (see Fig. S5a). Motif similarity scores range from 0 to 1, with higher scores indicating greater similarity between the identified motif and known transcription factor binding motifs.The analysis was performed using HOMER^51^. **f.** As in (a) but in *mtrr-1* mutants treated with RNAi targeting the indicated stress-responsive TFs and relative to ctrl RNAi. **g.** As in (f) but with RNAi targeting the mitochondria surveillance factor *cbp-3,* component of the PMK-3/MAPK mitochondria signaling pathway. **h.** As in (f) but with RNAi targeting components of UPR^mt^ pathway. For (a, b, f, h) Statistical significance was determined by two-way ANOVA with Dunnett’s Multiple Comparison test. For (c) and (g), Statistical significance was determined by multiple t-tests. For (a-c) and (f-h), data represent the mean of at least three independent biological replicates. *p<0.05, **p<0.01, exact p and n values are in Supplementary Table 2.

Both GSH and NAC supplementation increase intracellular sulfur metabolites, and *mtrr-1* mutants are predicted to exhibit increased flux through the transsulfuration pathway as a consequence of homocysteine accumulation^47^’^49^. Thus, we next tested antioxidants that do not directly contribute sulfur-containing metabolites. However, neither supplementation with the hydrophilic antioxidant vitamin C (Fig. 5a) nor with the lipophilic antioxidants vitamins A and E (Fig. 5b) restored repeat silencing. These results further fail to support oxidative stress as the primary cause of heterochromatin derepression in *mtrr-1* mutants.

We next asked whether persistence of oxidized, inactive METR-1 contributes to the heterochromatin phenotype. To directly test this possibility, we generated *mtrr-1 metr-1* double mutants and measured repetitive element expression relative to *mtrr-1* single mutants. Repeat expression was comparable between the two genotypes (Fig. 5c), indicating that removal of METR-1 does not suppress heterochromatin derepression caused by loss of MTRR-1. Consistently, *mtrr-1 metr-1* double mutants were phenotypically indistinguishable from *mtrr-1* single mutants (data not shown).

Together, these findings indicate that neither altered redox homeostasis nor persistent METR-1 inactivation is sufficient to explain the heterochromatin defects observed in *mtrr-1* mutants, supporting the existence of a distinct mechanism linking MTRR-1 to heterochromatin maintenance.

### Depletion of general stress-responsive transcription factors (TFs) has minor effects on repeat expression in *mtrr-1* mutants

Stress response pathways are activated in *mtrr-1* mutants. Therefore, we asked whether stress-responsive transcription factors (TFs) contribute to repeat derepression. To identify candidate regulators, we combined two complementary approaches. First, enrichment analysis of differentially expressed genes using WormExp^50^ revealed significant overrepresentation of targets of DAF-16, ELT-3, SKN-1, BLMP-1, and PQM-1 among genes differentially expressed in *mtrr-1* mutants (Fig. 5d; Fig. S5a). Among these, DAF-16 and ELT-3 were selected for functional analysis because they showed the strongest enrichment. Second, we performed *de novo* motif enrichment analysis of promoters of genes upregulated in *mtrr-1* mutants using HOMER^51^, identifying six significantly enriched motifs (Fig. S5b). The most significantly enriched motif (Motif 1) matched binding sites for the bZIP transcription factors CEBP-1 and ATF-2, which showed the highest motif matching scores (0.89 and 0.78, respectively) (Fig. 5e). Moreover, both *cebp-1* and *atf-2* transcripts were themselves upregulated in *mtrr-1* mutants (log2FC = 1.05 and 1.72, respectively). Thus, we selected CEBP-1 and ATF-2 for functional analyses.

To determine whether these TFs contribute to repeat derepression, we depleted each candidate individually by RNAi in *mtrr-1* mutants and quantified by RT-qPCR the expression of four repetitive elements derepressed in these animals. While depletion of *atf-2* had no detectable effect on repeat expression (Fig. 5f), knockdown of *daf-16, elt-3,* and *cebp-1* produced reductions in the expression of individual repetitive elements. However, none of these effects reached statistical significance. Specifically, *daf-16* and *cebp-1* knockdown reduced (CAA)« expression by approximately 40% and 24%, respectively, whereas *elt-3* knockdown reduced (TAGG)« expression by more than 30%. Together, these findings suggest that no single stress-responsive TF is sufficient to account for repeat derepression in *mtrr-1* mutants.

### Depletion of mitochondrial stress-responsive factors partially rescues repeat derepression in *mtrr-1* mutants

As *mtrr-1* mutants exhibit altered mitochondrial respiration, accumulation of long-chain acylcamitines, and activation of mitochondrial stress-response genes, we next asked whether mitochondrial stress signaling contributes to heterochromatin derepression. We therefore individually depleted key components of mitochondria-to-nucleus signaling pathways. Specifically, we used RNAi to target CBP-3, a component of the PMK-3/MAPK mitochondrial stress signaling pathway^42,52^, and the mitochondrial unfolded protein response (UPR^mt^) regulators ATFS-1 and DVE-1^53,54^. Remarkably, *cbp-3* knockdown partially rescued repeat derepression in *mtrr-1* mutants, significantly reducing the expression of two of the four repetitive elements analyzed. Compared with control RNAi, *cbp-3* knockdown decreased CER3 I CE expression by approximately 22% and (CAA)n expression by approximately 40% (Fig. 5g). Similarly, depletion of *atfs-1* partially reduced expression of three of the four repetitive elements analyzed, including the two repeats rescued by *cbp-3* (CER3 I CE and (CAA)n). Although these reductions did not reach statistical significance, CER3 I CE expression decreased by nearly 40%, whereas (CAA)ra and (TTG)n expression were reduced by more than 20%. In contrast, depletion of *dve-1* further enhanced repeat derepression in *mtrr-1* mutants, significantly increasing the expression of CER3 I CE and (CAA)n (Fig. 5h).

Together, these findings identify mitochondrial stress signaling as a regulator of heterochromatin derepression in *mtrr-1* mutants. While disruption of PMK-3/MAPK-dependent signaling through CBP-3 partially restores repeat silencing, depletion of *atfs-1* produces a similar, albeit non­significant, trend, whereas depletion of *dve-1* exerts the opposite effect. Thus, distinct components of the mitochondrial stress response differentially affect heterochromatin upon MTRR-1 loss.

### Inducing mitochondrial stress is sufficient to impair heterochromatin silencing

Our data indicate that MTRR-1 regulates heterochromatin independently of methionine synthesis and that altered mitochondrial homeostasis is a defining feature of *mtrr-1* mutants. Moreover, genetic perturbation of mitochondrial stress signaling partially restores repeat silencing in these animals. We therefore asked whether mitochondrial stress alone is sufficient to impair heterochromatin function.

To induce mitochondrial stress, we depleted by RNAi either *spg-7,* encoding a mitochondrial metalloprotease whose loss activates mitochondrial stress responses^37^’^38^, or *nuo-1,* encoding a complex I subunit whose depletion induces mild mitochondrial stress without completely disrupting electron transport^55,56^. Supporting the relevance of this approach, analysis of a published dataset from wild-type animals subjected to *spg-7* RNAi^38^ revealed that approximately 15% of the genes upregulated in *mtrr-1* mutants were also induced following *spg-7* depletion (hypergeometric test, P < 0.0001; Fig. 6a). These shared genes were enriched for biological processes including “defense response to other organism” and “autophagy of mitochondrion” (Fig. 6b).

**Figure 6.**
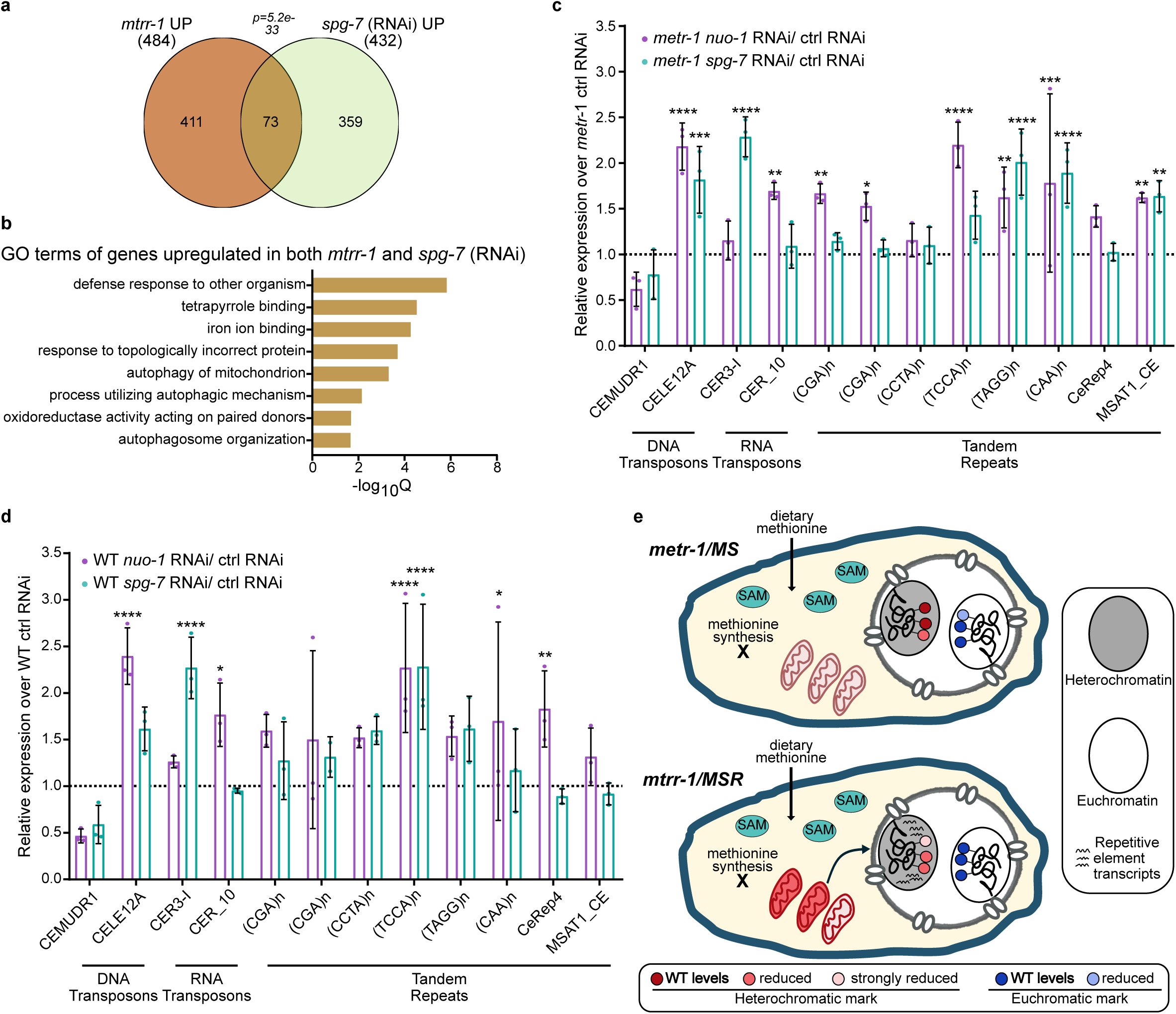
Inducing mitochondrial stress is sufficient to derepress repetitive elements in WT and *metr-1* mutants. **a**. Venn diagram showing the overlap between upregulated (log_2_FC≥ 1) genes in *mtrr-1* mutants and animals treated with *spg-7* RNAi38. Significance was determined using a hypergeometric test. **b**. GO terms significantly enriched among upregulated genes shared between *mtrr-1* mutants and *spg-7* RNAi treated animals. **c**. Barplot of the expression of the indicated repetitive elements, measured by qPCR, in *metr-1* mutants treated with the indicated RNAi and relative to ctrl RNAi (L4440). Statistical significance was determined by two-way ANOVA with Dunnett’s Multiple Comparison test. Data represent the mean of three independent biological replicates. *p<0.05, **p<0.01, ***p<0.001, ****p<0.0001. **d**. As in (c) but in wild type (WT) animals. Data represent the mean of three independent biological replicates. *p<0.05, **p<0.01, ***p<0.001, ****p<0.0001. **e**. Model: deletion of the methionine synthase *metr-1* (top) leaves histone methylation and gene expression largely unchanged, likely because dietary methionine is sufficient to maintain SAM levels. In contrast, loss of the methionine synthase reductase *mtrr-1* (bottom) results in a specific loss of heterochromatic histone marks and impaired heterochromatin silencing, despite unaffected global SAM levels. We propose that this phenotype arises from altered mitochondrial functionality, which is unique to *mtrr-1* mutants. For (c and d), exact p and n values are in Supplementary Table 2.

To determine whether mitochondrial perturbation affects heterochromatin silencing, we quantified the expression of twelve repetitive elements following *spg-7* or *nuo-1* depletion. We initially performed these experiments in a *metr-1* mutant background, reasoning that although impaired methionine synthesis alone is insufficient to disrupt heterochromatin, it might sensitize animals to mitochondrial stress. Despite differences in the magnitude and spectrum of repetitive elements affected, depletion of either *spg-7* or *nuo-1* consistently induced repetitive element derepression in *metr-1* mutants (Fig. 6c). Remarkably, the same mitochondrial perturbations also derepressed repetitive elements in wild-type animals (Fig. 6d), with effects largely comparable to those observed in a *metr-1* background. Together, these findings demonstrate that mitochondrial stress is sufficient to impair heterochromatin silencing, providing independent support for a functional link between mitochondrial homeostasis and heterochromatin maintenance.

## Discussion

Methionine availability is a key determinant of cellular SAM levels, the universal methyl donor required for histone methylation and other epigenetic modifications^1^. In recent years, dietary methionine has emerged as a key modulator of the epigenome through its effects on SAM abundance^6, n>12^. However, whether intracellular methionine synthesis contributes to the regulation of histone methylation—one of the best-studied and functionally relevant chromatin modifications from yeast to humans—has remained largely unclear.

Here, we show that dietary methionine is the principal contributor to cellular SAM pools, as deletion of methionine synthase (MS), METR-1 in C. *elegans,* leaves global SAM levels unchanged. Unexpectedly, we also uncover a specific role for MTRR-1, the methionine synthase reductase (MSR) in *C. elegans,* in regulating heterochromatin state and function independently of methionine synthesis. Unlike *metr-l/MS* mutants, *mtrr-1 /MSR* mutants display severe developmental defects and multiple signatures of altered mitochondrial functionality. Strikingly, genetic impairment of mitochondrial-to-nucleus signaling partially restores heterochromatin silencing in *mtrr-1* mutants, whereas experimental induction of mitochondrial stress is sufficient to compromise heterochromatin silencing in otherwise wild-type animals. We therefore propose that loss of *mtrr-1* alters heterochromatin by disrupting mitochondrial homeostasis rather than by impairing methionine synthesis (Fig.6e).

Loss of the methionine synthase METR-1 results in only mild changes in histone methylation, at both euchromatic and heterochromatic marks, along with minor effects on growth and gene expression. This is consistent with the fact that *C. elegans* obtains most of its methionine from the diet, making endogenous methionine synthesis largely dispensable under nutrient-replete conditions. Although global SAM and SAH levels remain unchanged in *metr-1* mutants, we cannot rule out changes in nuclear SAM availability that may contribute to the modest reductions in histone methylation observed. However, given the very modest effects observed in *metr-1* mutants, these histone methylation changes are likely insufficient to affect gene or repeat expression. This is consistent with work suggesting that histones can act as methyl sinks^57,58^ and that changes in histone methylation do not always translate into functional genomic consequences.

In contrast, loss of MTRR-1 causes derepression of repetitive elements, broad gene expression changes and selective reductions in histone methylation at multiple heterochromatic marks. The most strongly affected modification is H3K23me3, recently identified as a heterochromatin-associated modification in *C. elegans^13^’*^25,26^. A role for H3K23me3 in protecting heterochromatin from DNA damage has been reported in *Tetrahymena* thermophila^27^, and in mammals, H3K23 has been identified as a substrate of H3K9 methyltransferases^28^, thereby reinforcing its association with heterochromatin. H3K9me2, a conserved hallmark of constitutive heterochromatin with established repressive functions^31^, is also reduced in *mtrr-1* mutants, as are H3K9mel, H3K23me2 and H3K27me3. Although we cannot pinpoint which modification(s) drive the loss of silencing, we propose that reduced levels of one or more heterochromatic marks contribute to the derepression of repetitive elements.

Our findings in *C. elegans* resemble aspects of Mtrr deficiency in mice. Consistent with our metabolomics data (Fig. 4a), *Mtrr* hypomorphic mice exhibit no reduction in SAM levels or in the SAM/SAH ratio in most tissues^47^. Although Mtrr deficiency was initially reported to cause global DNA hypomethylation^16^, a more recent study demonstrated that this effect is locus-specific and may reflect characteristics of the genetic approach used in the particular mouse model studied^17^, leaving the role of MTRR in DNA methylation unclear. Importantly, our study is the first to reveal a histone methylation defect resulting from loss of methionine synthase reductase, highlighting an additional layer through which this enzyme influences genome regulation.

While the precise mitochondrial alteration remains to be determined, transcriptomic and metabolomic data indicate that mitochondrial homeostasis is specifically disrupted following loss of the methionine-cycle enzyme MTRR-1, but not its cognate methionine synthase METR-l/MS, as evidenced by long-chain acylcarnitine accumulation and transcriptional activation of mitochondrial and ER stress pathways. Consistent with these findings, *mtrr-1* mutants also exhibit increased basal mitochondrial oxygen consumption (OCR), whereas *metr-1* mutants do not. Although mitochondrial stress typically reduces basal OCR^59^, the increase in mitochondrial basal OCR observed in *mtrr-1* mutants does not necessarily indicate higher ATP production. Increased basal OCR can also result from enhanced proton leak or increased mitochondrial mass^60^. While mitochondrial O2 consumption primarily occurs via the electron transport chain to drive ATP synthesis, mitochondrial membrane-associated cytochrome P450 enzymes, O2-dependent monooxygenases, also consume O2 during detoxification reactions. Notably, 16 of the ∼75 cytochrome P450 genes in *C. elegans^61^’*^62^ are upregulated in *mtrr-1* knockout animals, suggesting increased cytochrome P450 activity. Thus, the elevated oxygen consumption in *mtrr-1* mutants may reflect enhanced cytochrome P450 activity rather than increased mitochondrial ATP production.

Female mice expressing an Mtrr hypomorph exhibit increased citrate synthase activity— indicative of elevated mitochondrial content^63^—but reduced P-oxidation when normalized to mitochondrial mass^64^. Although mitochondrial respiration and long-chain acylcamitine levels were unchanged in these mice, these findings suggest that mitochondrial homeostasis may also be perturbed by Mtrr deficiency in mammals. Supporting this, both *mtrr-1* mutant worms (this work) and Mtrr hypomorphic mice exhibit developmental defects^16^’^64^.

Methionine synthase reductase is a FAD-containing oxidoreductase that uses NADPH as its electron donor to reduce the cobalamin cofactor of methionine synthase (METR-1). Thus, an explanation for the mitochondrial and chromatin defects observed in *mtrr-1* mutants would be the accumulation of oxidized, inactive METR-1, which may itself promote oxidative stress, ER stress, and indirectly affect mitochondrial function. However, the absence of phenotypic or molecular rescue in *mtrr-\ metr-1* double mutants indicates that accumulation of inactive METR-1 is unlikely to be responsible for heterochromatin defects. Instead, these findings suggest that loss of MTRR-1 affects chromatin regulation through mechanisms independent of inactive METR-1 accumulation.

An alternative possibility is that reduced consumption of NADPH following MTRR-1 loss increases NADPH availability for other NADPH-dependent enzymes^65^’^66^, such as those operating in the endoplasmic reticulum or in oxidative detoxification pathways, thereby perturbing cellular redox homeostasis and indirectly contributing to the mitochondrial alterations observed in *mtrr-1* mutants. The inability of multiple antioxidant treatments to restore heterochromatin silencing argues against oxidative stress being the principal driver of heterochromatin defects. Nonetheless, these findings do not exclude a broader contribution of altered redox homeostasis to the phenotypes observed in *mtrr-1* mutants. Indeed, the modest increase in repeat derepression following antioxidant supplementation raises the possibility that further perturbation of an already altered redox homeostasis may exacerbate the phenotype.

Our genetic and functional experiments identify mitochondrial stress as a key contributor to heterochromatin regulation. Indeed, depletion of *cbp-3,* a component of the PMK-3/MAPK mitochondrial stress signaling pathway^42^, was the only intervention that significantly restored silencing of more than one repetitive element in *mtrr-1* mutants. Moreover, induction of mitochondrial stress in otherwise wild-type animals was sufficient to induce repetitive element derepression, demonstrating that mitochondrial perturbation done can compromise heterochromatin state.

Interestingly, depletion of *atfs-1,* a key regulator of the mitochondrial unfolded protein response (UPR^mt^), produced a similar, albeit non-significant, trend, whereas depletion of *dve-1,* another UPR^mt^-associated TF, further exacerbated repeat derepression. These findings indicate that distinct branches of the mitochondrial retrograde response differentially influence heterochromatin in *mtrr-1* mutants.

The incomplete rescue following depletion of individual signaling components may reflect incomplete RNAi-mediated depletion or functional redundancy among mitochondrial stress signaling pathways. Such redundancy is consistent with previous observations. For example, depletion of *alfs-1* under conditions of mitochondrial stress enhances *cbp-3* expression^52^, whereas loss of *atfs-1* also increases expression of *lin-65,* a component of the DVE-1-dependent UPR^mt^ pathway^67^, highlighting extensive crosstalk among mitochondrial retrograde signaling pathways. In addition, although depletion of individual stress-responsive TF did not significantly rescue repeat derepression, the modest reductions observed following *daf-16, elt-3* and *cebp-1* knockdown raise the possibility that these factors provide an additional layer of redundancy. Interestingly, the most enriched motif among genes upregulated in *mtrr-1* mutants is recognized by bZIP transcription factors, a family that includes ATFS-1 and CEBP-1 and whose members have previously been implicated in heterochromatin regulation^68^’^71^.

Together, these observations support crosstalk among mitochondrial retrograde signaling pathways and potentially other stress-responsive TFs in regulating heterochromatin.

Our findings are consistent with emerging evidence linking mitochondrial function to heterochromatin regulation. For example, in fission yeast, heterochromatin-mediated epimutations silence nuclear genes encoding mitochondrial proteins, thereby inducing mitochondrial dysfunction and conferring antifungal resistance^72^. Likewise, mitochondrial stress regulates heterochromatin in a mechanosensitive cellular model^73^. In C. *elegans,* the H3K9me2 methyltransferase MET-2 contributes to the transcriptional response to electron transport chain impairment^67^ together with DVE-1 and LIN-65, in parallel with ATFS-1. Furthermore, activation of the UPR^mt^ following *spg-7* knockdown partially depends on the H3K27me2/3 demethylases JMJD-1.2 and JMJD-3.1^74^ . Mitochondrial stress has also been shown to induce a switch in linker histone expression—from *his-24* (H 1.1) to *hil-1* (H1.0)—which is thought to influence chromatin compaction^75^. Notably, we observe this same switch in *mtrr-1* mutants, with downregulation of *his-24* (log2FC = −1.14, P = 0.0001) and upregulation of *hil-1* (log2FC = 1.02, P = 0.0003), further supporting a shared chromatin response between MTRR-1 deficiency and mitochondrial perturbation.

Clinically, mutations in MS and MSR are often treated as functionally equivalent as both lead to the detrimental accumulation of homocysteine and are managed with similar therapeutic strategies^76^’^77^. However, our findings show that MTRR-1 loss produces metabolic and chromatin phenotypes not seen in METR-1 loss, revealing additional roles for MTRR-1 in mitochondrial homeostasis and genome regulation. These results suggest that MSR deficiency in humans may be mechanistically distinct from MS deficiency.

Together, our findings reveal an unexpected, methionine-synthesis-independent role for methionine synthase reductase in maintaining mitochondrial homeostasis and heterochromatin integrity. Collectively, our results support a model in which disruption of mitochondrial homeostasis promotes heterochromatin derepression through mitochondrial stress signaling. This work broadens the functional scope of the methionine cycle and identifies mitochondrial physiology as a regulator of heterochromatin state, with broad implications for development, stress responses, genome stability, and disease.

## Acknowledgements

We thank the lab of William Kelly (Emory University), the Caenorhabditis Genetics Center, funded by the National Institutes of Health Office of Research Infrastructure Programs (grant no. P40 OD010440CGC); and the C. *elegans* Gene Knockout Project at the Oklahoma Medical Research Foundation for sharing strains. We thank SunyBiotech for support in generating the *mtrr-1* null allele. We thank Ann-Christine Konig and the Metabolomics and Proteomics Core Facility for performing LC-MS/MS on histones. We thank the lab of Merly Vogt (Helmholtz Munich) for support in performing the Seahorse experiments. V.F. is supported by the EpiCrossBorders PhD program. Work in the Scialdone lab is supported by the Helmholtz Association. Funding in the Cabianca lab was provided by the German Research Foundation (Deutsche Forschungsgemeinschaft) project numbers: 459488476, 507935196 and 559907625. D.S.C. thanks Helmholtz Munich for support.

## Author Contributions

Conceptualization: FRP, DSC

Methodology: FRP, AT, AS, AA, MW

Investigation: FRP, AT, IE, VF, PC, AA, MW

Visualization: FRP, DSC

Funding acquisition: DSC, TB, AS, RS

Supervision: DSC

Writing - original draft: FRP, DSC

Writing - review & editing: FRP, MW, DSC

## Materials and methods

### *C. elegans* maintenance and strains

Nematodes were grown on nematode growth medium (NGM) agar plates seeded with *Escherichia coli* OP50 at 20 °C, except where otherwise stated. All strains used in this study are listed in Supplementary Table 1.

Bacterial cultures were grown in LB for 16-18 h at 37 °C with shaking at 200 rpm. Cultures of KEIO collection mutants were supplemented with kanamycin at 25 pg/mL, whereas RNAi clones were cultured with ampicillin at 100 pg/mL.

### Constructs and strains

A complete deletion of *metr-1 (dca3* allele) was generated by CRISPR-Cas9. Briefly, two sgRNAs (sgRNAl: GAGTAGTCTTTTCGAGGAGT; sgRNA2:

GATCGCTCAACTTCTTCCTT) were used to introduce double-strand breaks flanking the *metr- 1* ORF. An ssDNA repair template (tttccagaaaATGACTCGAAGTAGTCTTTTCGAGtGAAGAAGTTGAGCGATGGCTATCACC AATTCTTGG) was supplied to generate a complete deletion of the gene. The strain was outcrossed twice prior to experimentation.

A complete deletion of the *mtrr-1* open reading frame (ORF) was generated by CRISPR-Cas9 by SunyBiotech. The strain was outcrossed twice prior to experimentation.

### Worm size measurements

For assessment of development in mutant and WT strains under standard conditions, synchronized Lis were grown on NGM plates seeded with OP50 bacteria. For assessing the development of *metr-1* and WT animals under methionine-reduced conditions, synchronized Lis were seeded on NGM plates seeded with KEIO collection clones carrying single deletions of *motB,* as control, or *metE,* to reduce the methionine levels.

After 72 h, animals were imaged in bright field using a Leica Ml 65 FC fluorescence stereo microscope equipped with a Leica K5 camera and controlled by LAS X software, at 2.5x magnification. The freehand tool in Cell-ACDC^78^ was used to outline each worm and measure its area. Automatic thresholding was applied using Gaussian filter sigma of 2.00, and Thresholding Algorithm Li to determine the mask for each worm, which was then used to calculate worm area.

### Antioxidants supplementation

Antioxidant supplementation was performed by adding the indicated compounds to nematode growth medium (NGM) plates at the following final concentrations: 5 mM reduced glutathione (GSH), 10 mM N-acetyl-L-cysteine (NAC), 5 mM vitamin C, 5 pM vitamin A, and 200 pM vitamin E. Because vitamins A and E were dissolved in DMSO, NGM plates containing 0.02% DMSO served as controls.

### Collection of Lis

Gravid adults grown at 20 °C on peptone-enriched plates seeded with a 2x concentrated overnight culture of *E. coli* OP50 were subjected to standard hypochlorite treatment to isolate embryos, which were then hatched overnight in M9 buffer on a roller at 20 °C for 16-18 h to obtain synchronized LI larvae. Lis were then fed for 2 h 45 min to 3 h on peptone-rich plates, seeded with a 2x concentrated overnight culture of *E. coli* OP50, and subsequently washed three times with M9 to remove residual bacteria. After washing, Lis were snap-frozen in liquid nitrogen for metabolomics and histone mass spectrometry analyses. For RNA extraction, samples were snap-frozen in Trizol (15596026, Thermo Fisher).

For *nuo-1* and *spg-7* RNAi treated Lis, worms were grown on RNAi plates seeded with the corresponding 2x concentrated RNAi bacteria. Synchronized Lis were initially plated on L4440 control plates, then after 48 h L4 animals were transferred to L4440, *nuo-1,* or *spg-7* RNAi plates. After ∼30 h of feeding on the respective RNAi bacteria, embryos were obtained from hypochlorite treatment of gravid adults and were allowed to hatch overnight in M9 (16-18 h). The resulting synchronized Lis were fed for 3 h on the respective RNAi plates, washed three times with M9 to remove bacteria, and then snap-frozen in liquid N2 with Trizol (15596026, Thermo Fisher) for RNA extraction.

### Isolation of Histones and Mass Spectrometry

For histone extraction, approximately 300,000 Lis per replicate were lysed using 18 cycles of bead-beater lysis with 200 pL zirconia/silica beads (0.5 mm diameter) and 1 mL of Lysis Buffer (LB) (100 mM HEPES, pH 7.5; 50 mM NaCl; 1% Triton X-100; 0.1% NaDeoxycholate; 1 mM EDTA; 10 mM NaButyrate; protease inhibitors).

To remove soluble proteins, the lysate was centrifuged at high speed (13,000 rpm, 10 min) and the supernatant discarded. The insoluble pellet was resuspended in 300 pL of LB and sonicated to shear DNA and solubilize chromatin. Sonication was performed with 12 cycles on a QSonica instrument (30 s on, 30 s off) at 80% amplitude. After sonication, samples were centrifuged for 10 min at 4 °C and 13,000 rpm, and the supernatant was collected for analysis.

Histone proteins were prepared for LC-MS analysis using a hybrid chemical derivatization protocol^79^ adopted for in-gel sample preparation as described previously^80^. In brief, proteins were resolved on 4—20% polyacrylamide gels (Novex Wedge Well Tris-Glycin-Minigel, Invitrogen) followed by Coomassie staining. Histone protein bands were excised from the gel and destained in a destaining buffer (100 mM triethylammonium bicarbonate in 50% acetonitrile). After destaining, the gel pieces were dehydrated with 200 pi of 100% acetonitrile for 10 min at room temperature after which acetonitrile was discarded. Propionylation solution was prepared by mixing 50 mM TEAB (pH 8.5) and freshly prepared 1% (v/v) propionic anhydride solution in water at a 100:1 ratio. Immediately after preparation, 100 pi of propionylation solution was added to the dehydrated gel pieces followed by 10 min incubation at room temperature. The propionylation reaction was quenched by the addition of 10 pi of 80 mM hydroxylamine and subsequent incubation for 20 min at room temperature. The propionylation solution was discarded and gel pieces were dehydrated with 200 pi of 100% acetonitrile for 10 min at room temperature. After this, the acetonitrile solution was discarded and 20 pi of 50 ng pi— 1 trypsin solution in 100 mM TEAB (pH 8.5) was added. Trypsin digestion was performed overnight at 37 °C. The next day, 50 pi of 100 mM TEAB (pH 8.5) solution was added to each sample followed by 30 min incubation in a thermo shaker (37 °C, 1,500 rpm). A 1% (v/v) solution of phenyl isocyanate in acetonitrile was freshly prepared and 15 pi added to each sample and incubated for 60 min at 37 °C. The samples were acidified by the addition of 24 pi 1% trifluoroacetic acid. Peptides were desalted with Cl 8 spin columns (Thermo Fisher Scientific) according to the manufacturer’s instructions, dried in a speed-vac, resuspended in 50 pi 0.1% trifluoroacetic acid and subsequently used for LC-MS analysis.

The acidified histone peptide digests were analyzed on a Q Exactive HF mass spectrometer (Thermo Fisher Scientific) coupled to a nano-RSLC system (Ultimate 3000, Dionex). Samples were automatically loaded onto a pre-column (300 pm inner diameter * 5 mm, packed with Acclaim PepMaplOO Cl 8, 5 pm, 100 A; LC Packings) and subsequently eluted and separated by reversed-phase chromatography on an analytical column (HSS T3 M-class column, 25 cm, Waters). Peptides were separated using the following gradient at a flow rate of 250 nl min^-1^: solvent B (95% acetonitrile, 0.1% formic acid) was increased from 1% to 25% over 40 min, from 25% to 40% over 20 min, and from 40% to 85% over 5 min. Full-scan MS spectra and MS/MS fragment spectra were acquired in the Orbitrap at resolutions of 120,000 and 15,000, respectively, with maximum injection times of 50 ms for both MS and MS/MS. Up to 10 of the most intense precursor ions were selected for higher-energy collisional dissociation, depending on signal intensity. Dynamic exclusion was disabled.

For the quantification of histone PTMs, MS raw data files were manually analyzed using Skyline software. Briefly, a list of unmodified and differentially modified histone peptides was manually compiled and used to assess histone modification states in each sample. All lysine residues not bearing acetylation or methylation were assumed to be propionylated, and all peptide N-termini were assumed to be modified with phenyl isocyanate. For each modified histone peptide, the relative abundance was estimated by dividing its peak area by the sum of the areas corresponding to all of the observed forms of that peptide (that is, all peptides sharing the same amino acid sequence).

For Figure lc, the relative levels of unmodified and modified H3K27 and H3K36 were calculated by combining both canonical H3 and H3.3 peptides, according to the relative abundance of the two histone H3 variants.

### Total RNA extraction

Trizol (15596026, Thermo Fisher)-collected, liquid nitrogen snap frozen LI larvae were thawed and frozen by five successive transfers between liquid nitrogen and a water bath at 42 °C, followed by vigorous shaking for 2.5 min with 5 cycles of 30 s on and 30 s off at room temperature. Total RNA was subsequently extracted using the RNeasy Mini Kit (74104, Qiagen) according to the manufacturer’s instructions, including an on-column DNase digestion (79254, Qiagen).

### Repetitive elements and gene expression analysis by RNA-seq

RNA quality was assessed using an RNA Pico Chip on a 2100 Bioanalyzer (Agilent Technologies). NGS libraries were prepared with 500 ng of input RNA using the TruSeq Stranded Total RNA Library Prep Kit, following the manufacturer’s instructions. Sequencing was performed on a NovaSeq 6000 or NovaSeq X Plus sequencer with 150-cycle paired-end reads. Sequencing reads were aligned to the C. *elegans* reference genome (cell) using STAR, with multimapping reads retained (options —winAnchorMultimapNmax 200 and — outFilterMultimapNmax 100).

Quantification of repetitive elements was performed using TElocal, a locus-level extension of TEtranscripts^32^, with the *C. elegans* transposable element annotation file provided by the Hammel Lab (https://www.mghlab.org/software/telocal). For differential expression analysis, lowly expressed repeats were filtered out by excluding elements with fewer than 5 reads in all sample. Differential expression analysis was then performed using edgeR^81^.

Gene expression quantification was performed using featureCounts^82^ with uniquely mapped reads. Lowly expressed genes were filtered out by requiring a minimum of 10 reads in each sample, for all samples. Differential gene expression analysis between WT and mutant strains was conducted using DESeq2^83^. Differentially expressed repeats and genes were defined using a threshold of |log2FC| > 1 and FDR < 0.05. Overlap and comparisons between different genotypes were performed indirectly by comparing the lists of differentially expressed genes obtained for each condition relative to WT control. Enrichment analysis for up- and downregulated genes of *mtrr-1* mutants was performed using the WormBase Enrichment Analysis tool.

Identification of gene targets of transcription factors was conducted separately for up- and downregulated genes using WormExp (https://wormexp.zoologie.uni-kiel.de/). In addition, *de novo* motif discovery was performed using HOMER^51^. Promoter regions (-lOOObp and +250bp relative to the transcription start site) of the upregulated genes in *mtrr-1* mutants were analyzed using findMotifsGenome.pl.

### Quantitative PCR with reverse transcription

Complementary DNA (cDNA) was synthesized using either 500 or 1,000 ng of total RNA as template with the Maxima H Minus cDNA Synthesis Master Mix (Ml 661, Thermo Scientific). Expression analysis was performed by real-time PCR (LightCycler480 Roche) using the PowerUp SYBR Green Master Mix (A25742, Life Technologies). Relative quantification was performed by normalizing to the housekeeping gene *cdc-42,* calculating the ACt for each target within the sample. Fold change relative to the control genotype or condition was calculated as - 2^A^AACt. All experiments were repeated two or three times. Primer sequences are listed in Supplementary Table 3.

### Microscopy

Microscopy was performed using a live-cell imaging system (Confocal Spinning Disk Microscope, Visitron Systems GmbH) equipped with a Nikon Eclipse Ti2 microscope, Plan Apo X lOOx/1.45 oil, Plan Apo X 60x/1.40 oil, and Plan Apo X 10x objectives, a Yokogawa CSU-W1 confocal scanner unit, VS-Homogenizer, an Electron Multiplying CCD camera (Andor iXon Series), and VisiView software for acquisition.

For image acquisition, plates were washed with M9 buffer to remove adults and larvae, leaving only embryos, which were allowed to hatch on the plate for 3.5—4 h. Immediately before imaging, freshly hatched Lis were collected in M9. Live imaging was performed on 2% agarose pads supplemented with 0.15% sodium azide (Interchim) to immobilize the worms, as previously described^84^. Images for fluorescence intensity analysis were acquired using the Plan Apo X 10x objective, consisting of 60 z-stacks with a z-spacing of 1 pm.

### Image analysis and GFP Quantification

Z-stack images were sum-projected prior to analysis. Next, animals were selected using the freehand tool in Cell-ACDC^78^ using automatic thresholding with Gaussian filter sigma of 2.00, and Thresholding Algorithm Li to determine the mask for each worm. The mask was then used to calculate worm area, and maximum and mean GFP intensities across the whole body. Background intensity of each image was subtracted for animals’ GFP intensity measurements. Animals were filtered by area to select a population of similar sizes, minimizing confounding effects of size on GFP intensity measurements.

### RNAi experiments

RNAi was performed by feeding on plates^85^. RNAi clones were obtained from the Ahringer library (Source BioScience Ltd.), except for the *metr-1* clone, which was obtained from ORFeome RNAi Library (Vidal library). All clones, derived from single colonies, were sequenced to confirm target specificity. Experiments were conducted at 20 °C on NGM agar plates supplemented with 100 pg/mL carbenicillin and 100 pM isopropyl-P-d-thiogalactoside and seeded with bacteria producing double-stranded RNA, grown overnight (16-18h) at 37 °C. As a negative control, the RNAi clone containing the empty L4440 vector (Fire vector library) was used. In every experiment, *let-607* RNAi was performed in parallel to verify the efficiency of each batch of RNAi plates, as this treatment reliably produces a phenotype of developmental arrest.

### Liquid Chromatography-Mass Spectrometry-based metabolite analysis

All solvents and chemicals were of analytical grade or higher. For LC-MS, only LC-MS grade solvents have been used. C. *elegans* samples were extracted as previously described^86^. Normalization was performed using read-outs from a Hoechst assay as described previously^87^. For the HILIC separation, the LC-HRMS/MS measurements were performed using an Agilent 1290 Infinity II BioLC (Agilent Technologies, Waldbronn, Germany) coupled to a Sciex ZenoTOF 7600 tandem mass spectrometer (MS/MS) equipped with an OptiFlow Turbo V electrospray ionization (ESI) source (Sciex, Darmstadt, Germany). Prior to LC-MS/MS measurement, two aliquots of dried sample extracts were reconstituted with LC-compatible solvents that contained chemical standards at fixed concentrations. One aliquot was analyzed in positive ionization mode. Metabolite separation was performed using an Agilent Infinity Poroshell 120 HILIC-Z column (100 mm x 2.1 mm ID, 2.7 pm particle size, PEEK-lined) with a gradient from eluent A (100% H2O + 10 mM ammonium formate / 0.1% formic acid) to eluent B (10% H2O / 90% ACN +10 mM ammonium formate / 0.1% formic acid). Flow rate was set to 0.5 mL/min and the column temperature was set to 40°C. Injection volume was 5 pL. Flow rate was set to 0.5 mL/min and column temperature was set to 40°C. The second aliquot was analzyed in negative ionization mode and metabolite separation was achieved using a Waters Atlantis Premier BEH Z-HILIC (100 mm x 2.1 mm ID, 1.7 pm particle size) and a gradient from eluent A (100% water +10 mM ammonium acetate / ammonium hydroxide (pH9)) to eluent B (10% water / 90% acetonitrile +10 mM ammonium acetate/ ammonium hydroxide (pH9)). Flow rate was set to 0.5 mL/min and column temperature was set to 40°C. Injection volume was 5 pL in both ionization modes. Detailed LC and MS parameters for both methods can be found in^88^. Initial QC of the data contained checking for internal standards regarding their RT, m/z and intensity. All internal standards have been detected with 5 ppm or less in mass error. Data analysis was performed in a targeted fashion for identified metabolites. Metabolites were identified by comparison to in-house reference standards, publicly available reference spectra, and manual interpretation of fragmentation spectra. Data analysis was performed in Sciex OS 4.0 (Sciex, Darmstadt, Germany). Peaks for all metabolites indicated were integrated with an XIC width of 0.02 Da and a Gaussian smooth width of 3 points using the MQ4 peak-picking algorithm. Peak areas were exported to a .txt file and normalized according to the Hoechst read of the respective sample.

### Seahorse

Synchronized LI worms were grown at 20 °C on NGM plates seeded with OP50 bacterial cultures, to obtain synchronized Day 2 adults for all strains. As *mtrr-1* mutants grow slower, they were bleached and seeded as Lis one day earlier than WT and *metr-1* mutants, *mtrr-1* mutants were allowed to grow for approximately 4 days, while *metr-1* and WT animals were grown for 3 days. The developmental stage of the animals was confirmed before starting Seahorse experiments, ensuring that worms were of similar size and had laid comparable numbers of embryos on the plates.

Worm oxygen consumption was measured as previously described^89^, using the Agilent Seahorse XFp Analyzer. Day 2 adult worms (8-17 per well) were collected and washed three times with M9 buffer. Working solutions were prepared at the following final concentrations: of FCCP (carbonyl cyanide p-trifluoromethoxyphenylhydrazone) 100 pM in M9 andNaN3 (Sigma- Aldrich) 400 mM in dH2O. Oxygen consumption rate (OCR) measurements were performed 5 times under basal conditions, 9 times after FCCP injection to measure maximum OCR and spare respiratory capacity, and 4 times after NaN3 injection to determine non-mitochondrial OCR. Each cycle consisted of 1 min mixing, 2 min waiting, and 3 min measurement. Three independent assays were conducted, and values were normalized per worm for each condition.

For basal OCR, the lowest and highest values were excluded, keeping the three median values to calculate the average basal OCR per well. For maximum OCR, the highest 6 out of 9 values were used. Non-mitochondrial OCR was calculated by excluding the lowest and highest values keeping the two median values to calculate the average per well.

We subtracted the non-mitochondrial OCR values for each well from the corresponding basal OCR and maximum OCR to determine mitochondrial basal and maximum respiration, respectively. Spare mitochondrial respiratory capacity was calculated by subtracting the based OCR from the maximum OCR.

### Statistics and reproducibility

Statistics analysis and data visualization were performed in Excel (Version 2308), R (4.3.1), or GraphPad Prism (8.0.1). All statistical tests, exact P values, n and number of biological replicas are listed in Supplementary Table 2.

### Data reporting

No statistical method was used to predetermine sample size. Sample sizes were chosen based on previous literature and what is common practice in the field. The experiments were not randomized. Investigators were not blinded to group allocation of samples (genotype/treatment) during data collection and analysis.

## Supplementary Material for

**Figure S1. Related to Figure 1.**
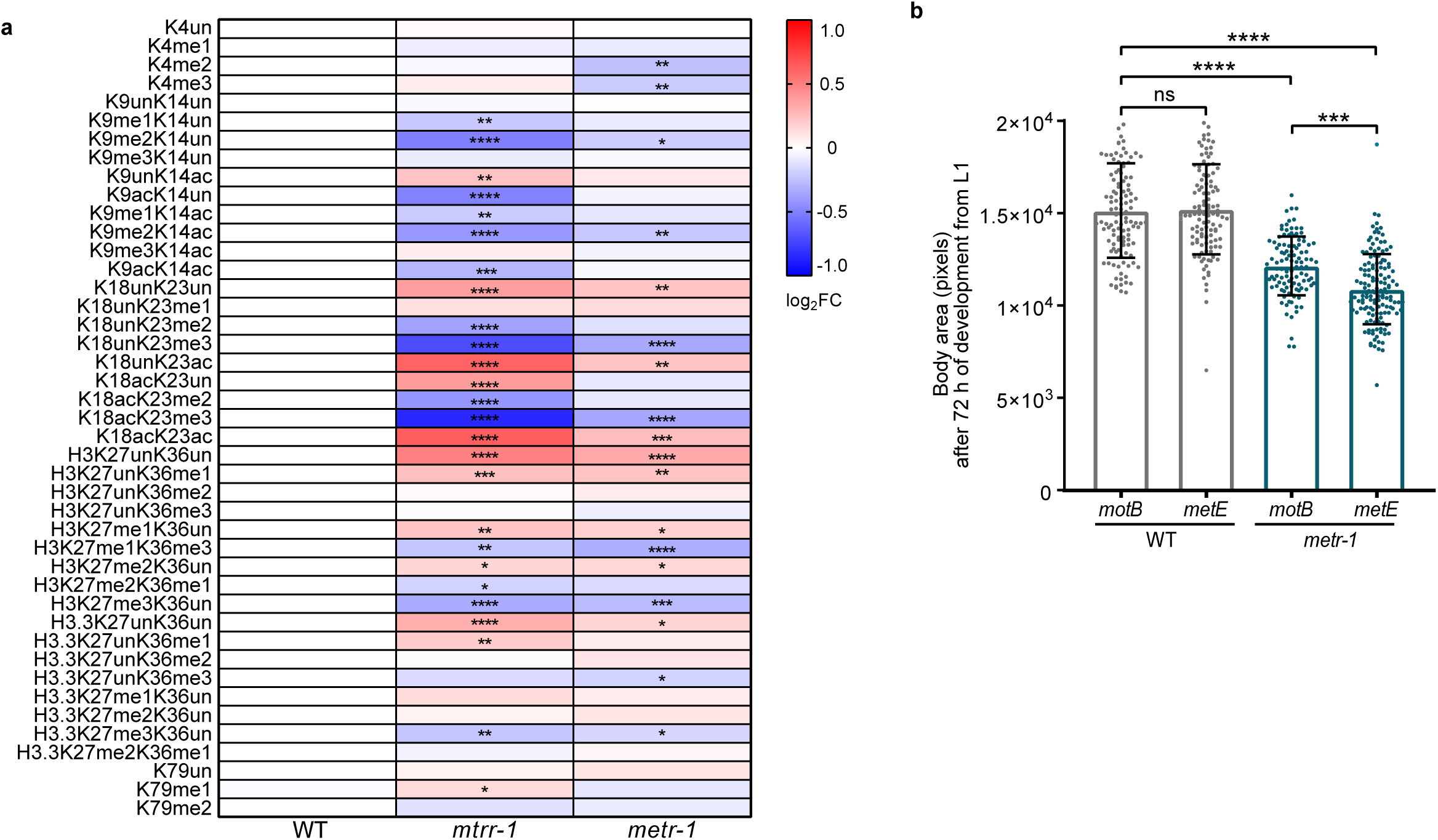
a. Heatmap of the relative log_2_ fold change (log_2_ FC) in the indicated histone H3 modifications in *mtrr-1* and *metr-1* mutant L1s, relative to WT worms. Statistical significance was determined by two-way ANOVA with Dunnett’s Multiple Comparison test. Data represent the mean of four independent biological replicates. *p<0.05, **p<0.01, ***p<0.001, ****p<0.0001. **b.** Barplot of the quantification of the body area (pixels) of wild type (WT) and *metr-1*(CRISPR) animals grown on *motB* (control) or *metE* (reduced methionine) *E. coli* bacterial mutants on NGM plates and measured after 72 h from the onset of the L1 stage. Statistical significance was determined by Kruskal-Wallis test. Data represent the mean of three independent biological replicates, each including at least 20 animals. ***p<0.001, ****p<0.0001; ns = not significant. Exact p and n values are in Supplementary Table 2.

**Figure S2. Related to Figure 2.**
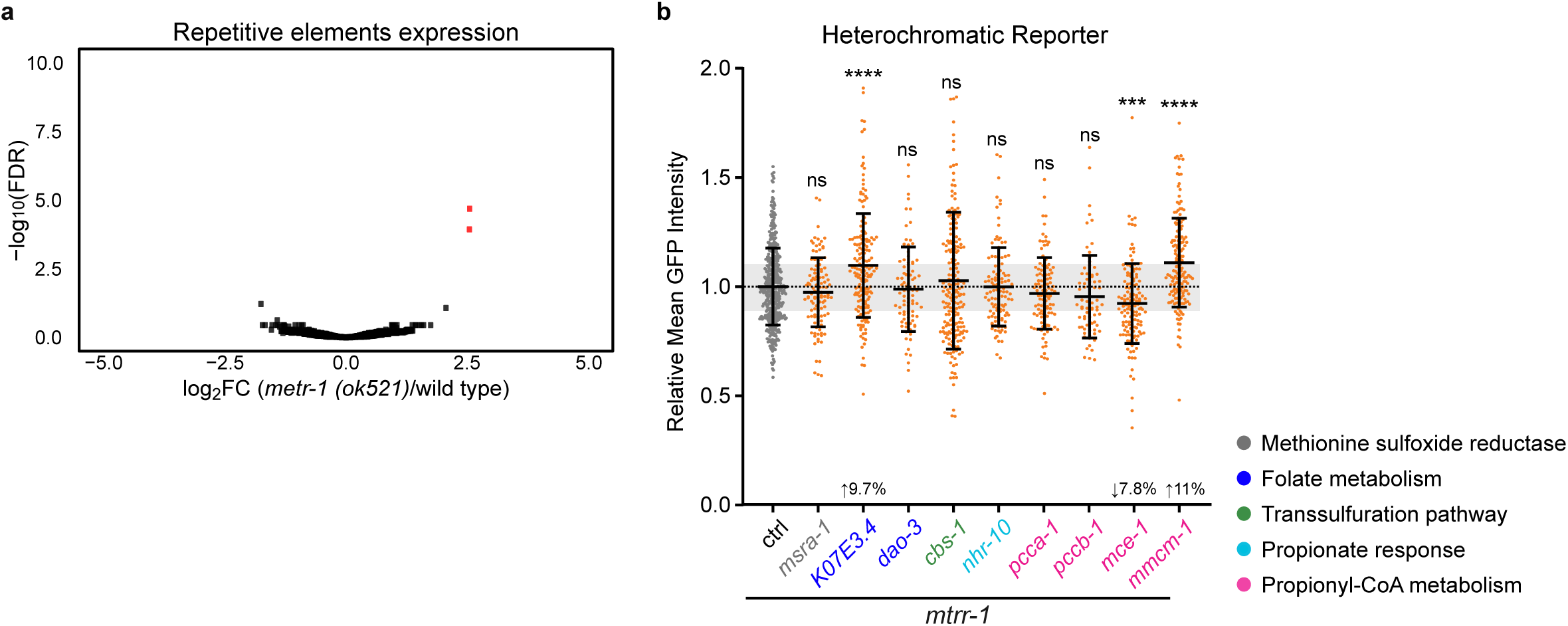
**a**. Volcano plot of repetitive elements in *metr-1 (ok521)* mutants compared to wild type (WT). Significant up- and down-regulated transcripts are highlighted (in red FDR <0.05 and log_2_FC s|l|, data represent the mean of three independent biological replicates). **b**. Beeswarm plot of the quantification of the mean GFP intensity expressed from the heterochromatic reporter (Fig. 2d) in *mtrr-1* L1s treated with the indicated RNAi, relative to the control (ctrl) RNAi L4440. The shaded area denotes a ±10% range, which we considered as cutoff for defining meaningful changes. Statistical significance was determined by Kruskal-Wallis test. Data are from at least two independent biological replicates, each with at least 20 animals. ***p<0.001, ****p<0.0001, ns= not significant. For b, exact p and n values are in Supplementary Table 2.

**Figure S3. Related to Figure 3.**
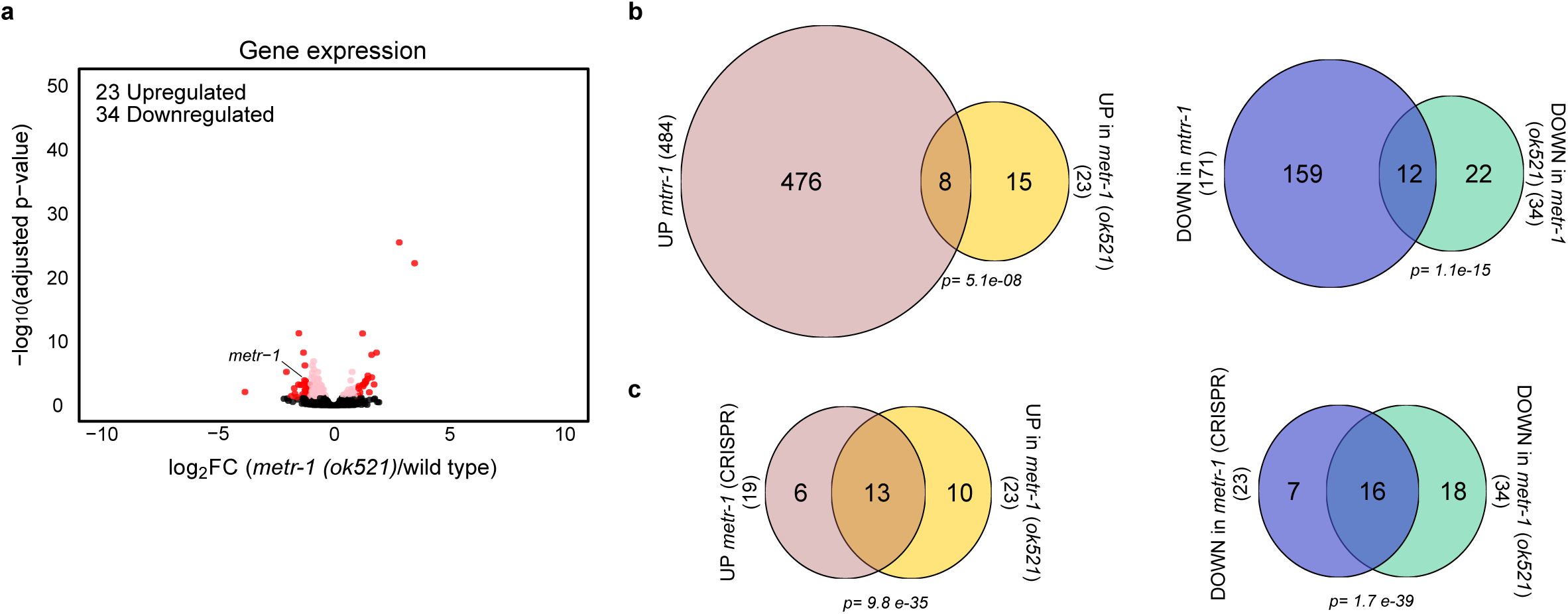
**a**. Volcano plot of gene expression in *metr-1 (ok521)* mutants compared to wild type (WT). Significant up- and downregulated transcripts are highlighted (in red FDR <0.05 and log_2_FC s|l|, in pink FDR <0.05 and log_2_FC <|1|, data represent the mean of three independent biological replicates). **b**. Venn diagram showing overlap between upregulated (left) and downregulated (right) genes in *metr-1(ok521)* and *mtrr-1* mutants. **c**. As in (b) but showing the overlap between upregulated (left) and downregulated (right) genes between *metr-1(ok521)* and *metr-1*(CRISPR) mutants.

**Figure S4. Related to Figure 4.**
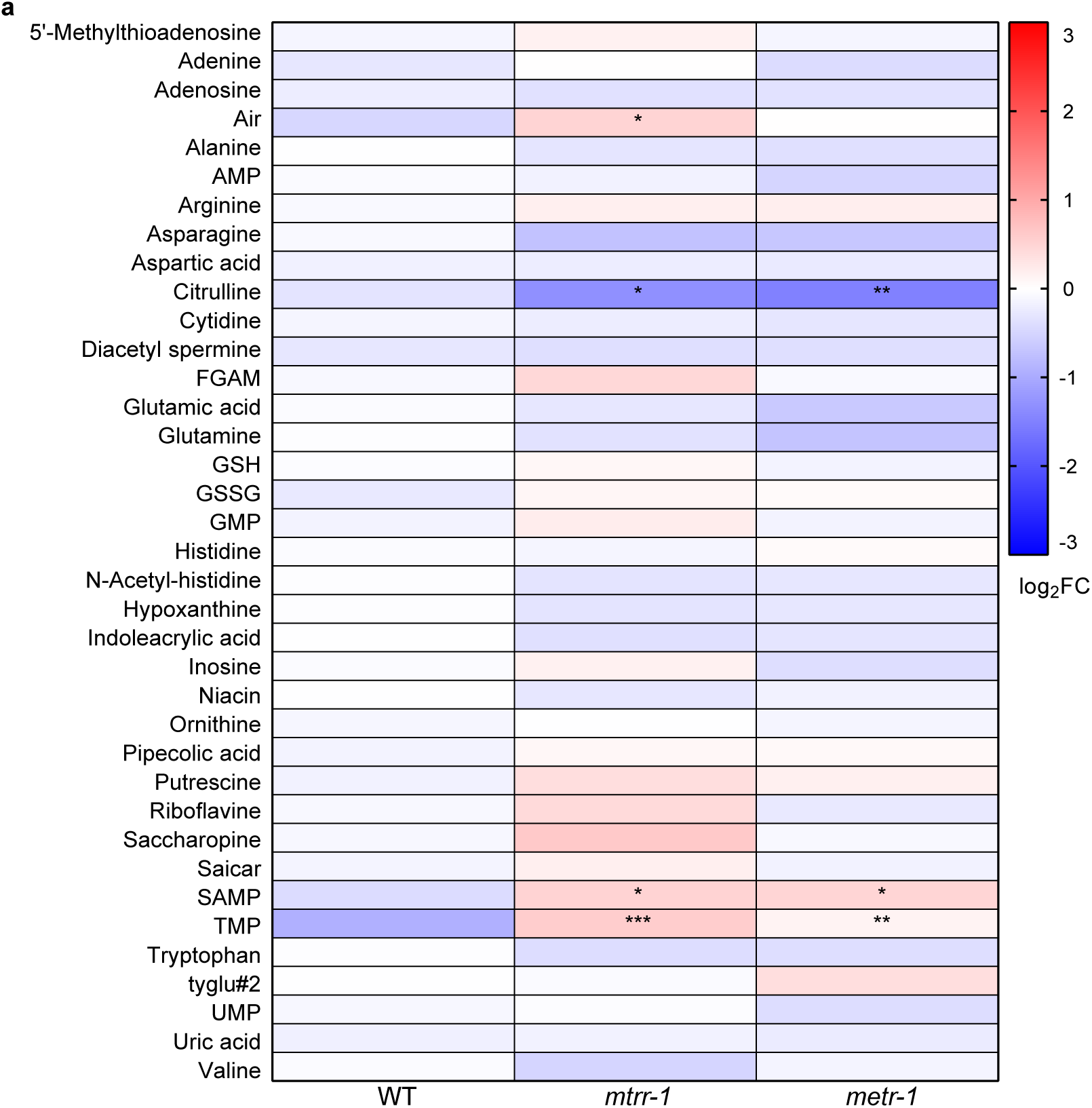
**a**. Heatmap of log_2_ Fold Change (FC) of the indicated metabolites in *mtrr-1* and *metr-1* mutants, relative to wild type (WT) worms. Statistical significance was determined by 2-Way ANOVA with Dunnett’s Multiple Comparison test. Data represent the mean of seven independent biological replicates. *p<0.05, **p<0.01, ***p<0.001. Exact p and n values are in Supplementary Table 2.

**Figure S5. Related to Figure 5.**
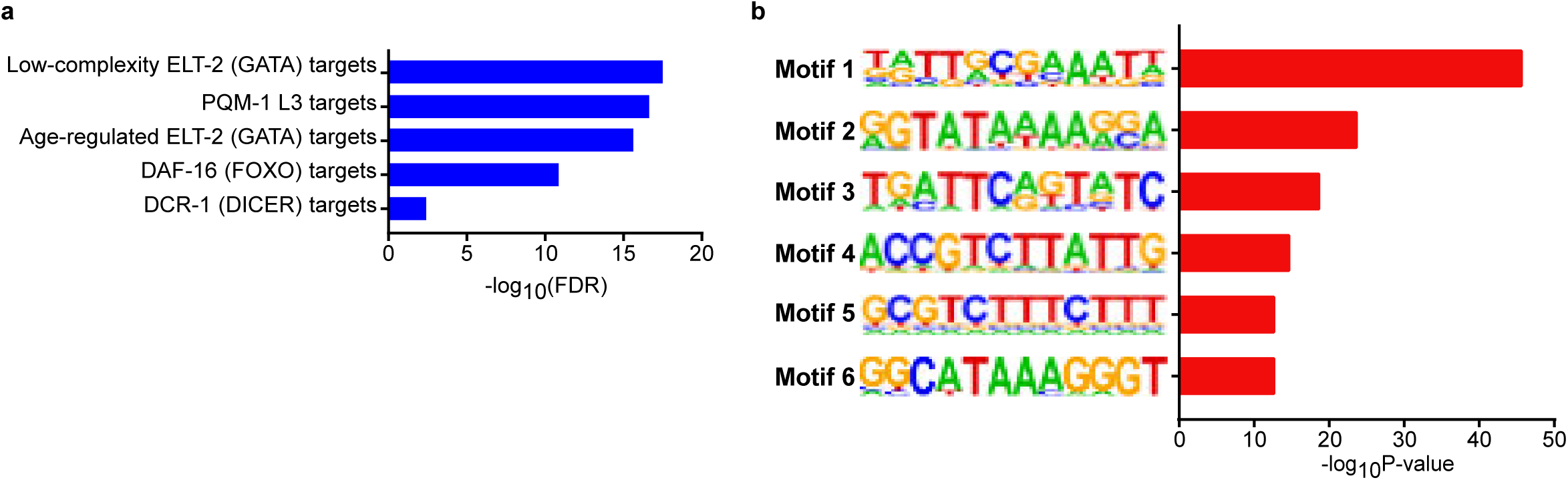
**a**. Enrichment analysis of downregulated genes in *mtrr-1* mutants among transcription factor (TF) targets. The analysis was performed using WormExp^50^ (https://wormexp.zoologie.uni-kiel.de/wormexp/). **b.** Significantly enriched motifs in the promoter regions of upregulated genes in *mtrr-1* mutants, identified by *de novo* motif analysis using HOMER^51^.

**Supplementary Table 1:**
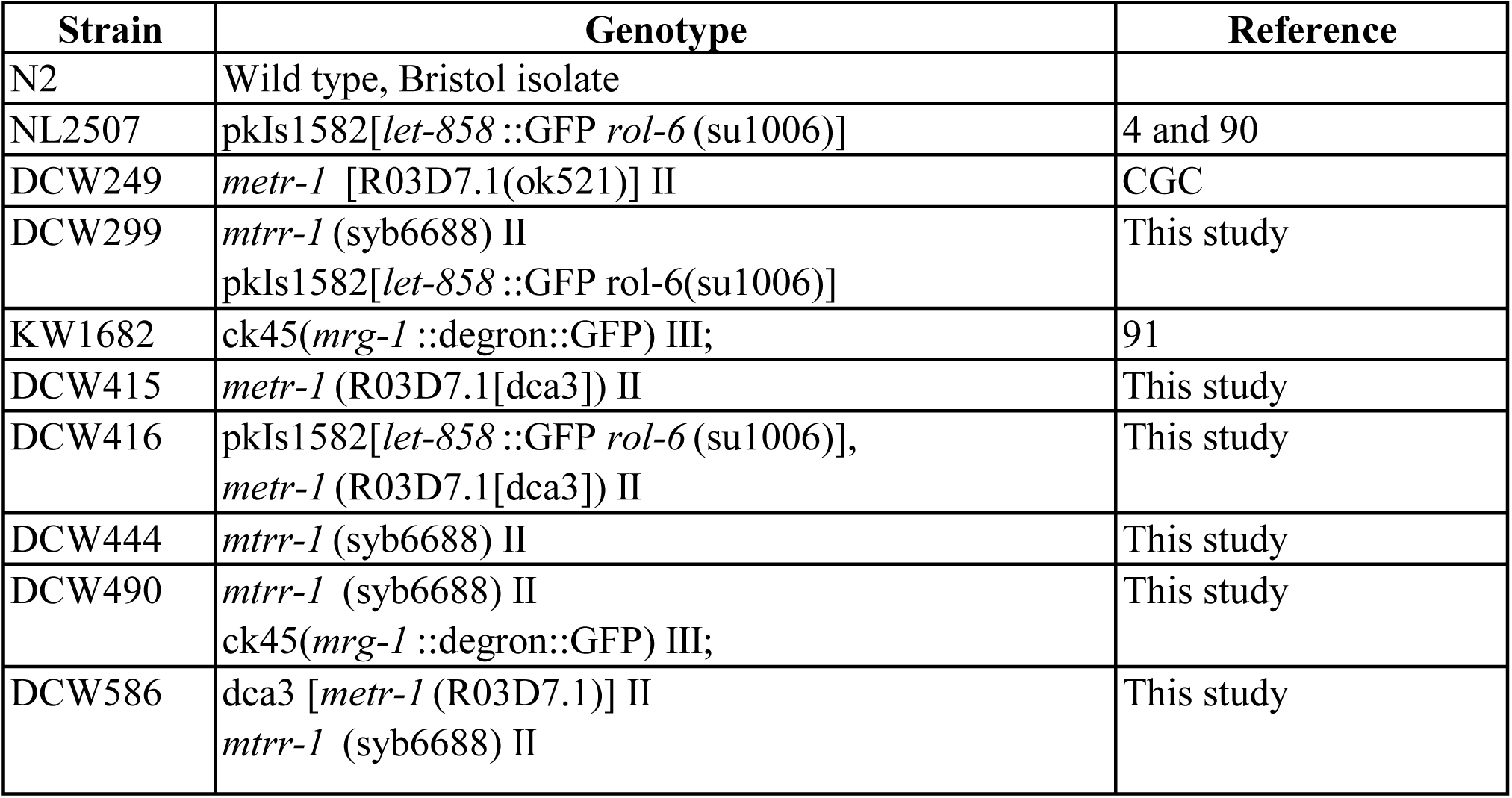
List of worm strains used in this study.

| **Strain** | **Genotype** | **Reference** |
| --- | --- | --- |
| N2 | Wild type, Bristol isolate |  |
| NL2507 | pkIs1582[*let-858* ::GFP *rol-6* (su1006)] | 4 and 90 |
| DCW249 | *metr-1* [R03D7.1(ok521)] II | CGC |
| DCW299 | *mtrr-1* (syb6688) II<br>pkIs1582[*let-858* ::GFP *rol-6* (su1006)] | This study |
| KW1682 | ck45(*mrg-1* ::degron::GFP) III; | 91 |
| DCW415 | *metr-1* (R03D7.1[dca3]) II | This study |
| DCW416 | pkIs1582[*let-858* ::GFP *rol-6* (su1006)],<br>*metr-1* (R03D7.1[dca3]) II | This study |
| DCW444 | *mtrr-1* (syb6688) II | This study |
| DCW490 | *mtrr-1* (syb6688) II<br>ck45(*mrg-1* ::degron::GFP) III; | This study |
| DCW586 | dca3 [*metr-1* (R03D7.1)] II<br>*mtrr-1* (syb6688) II | This study |

**Supplementary Table 2:**
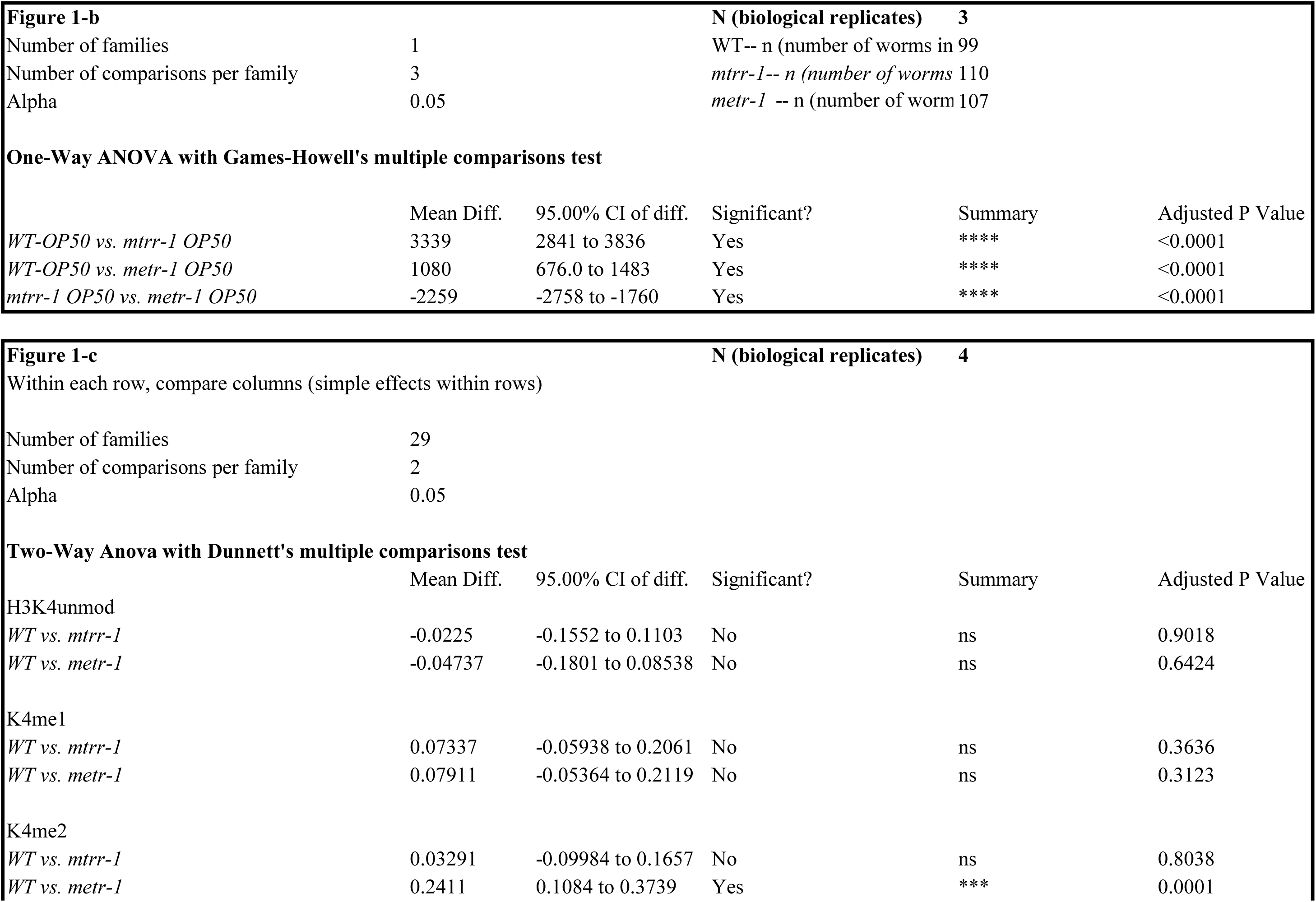
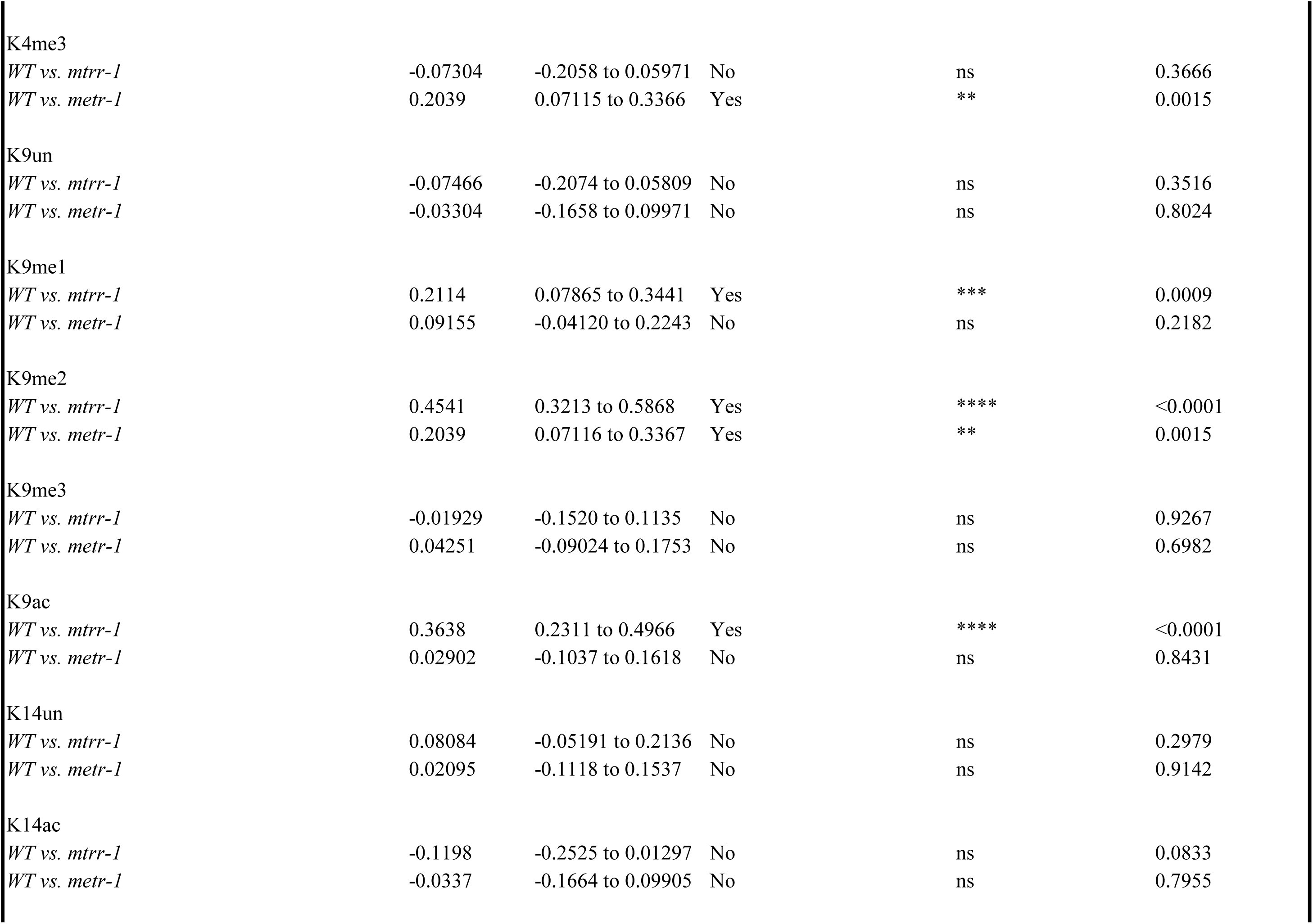
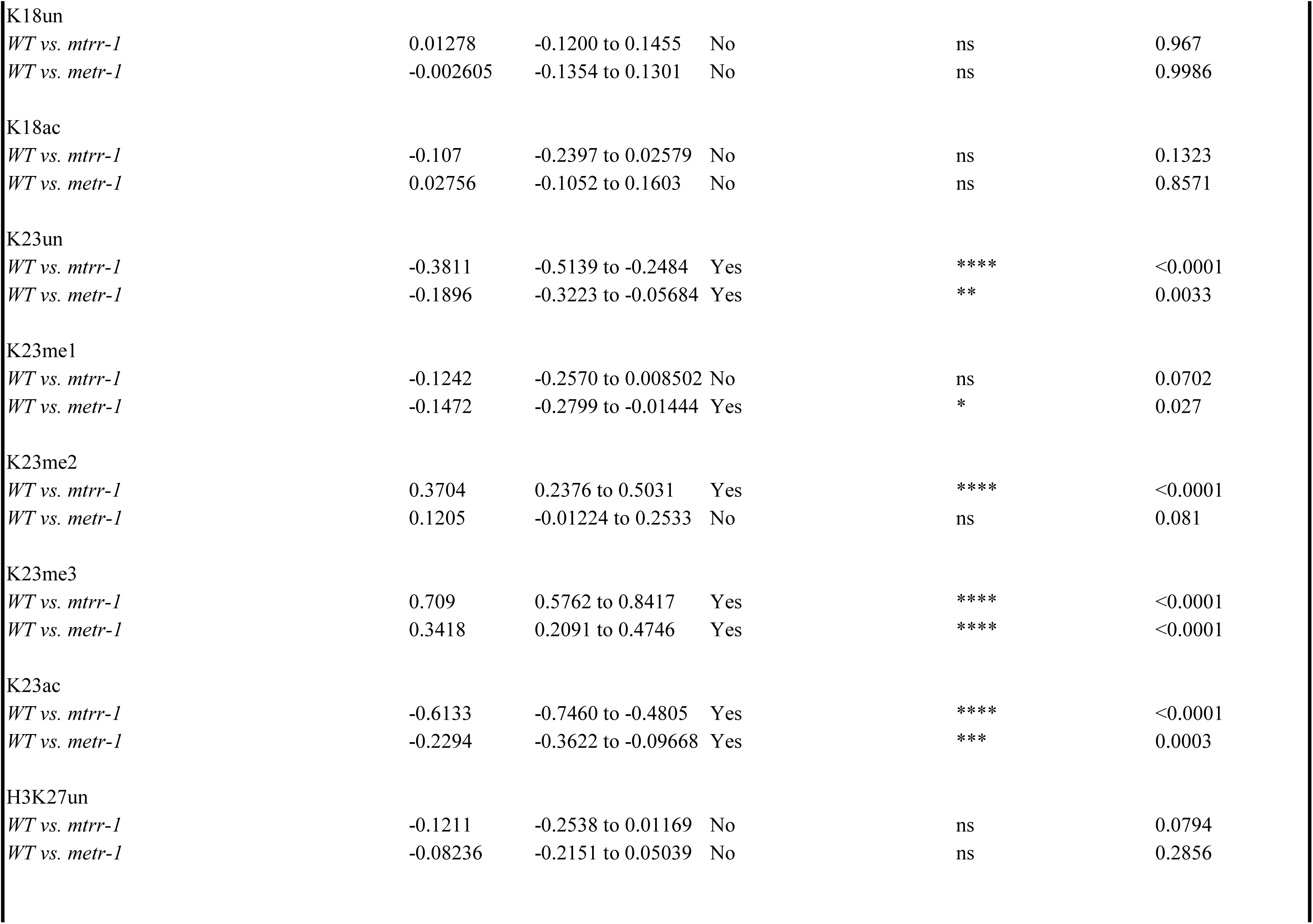
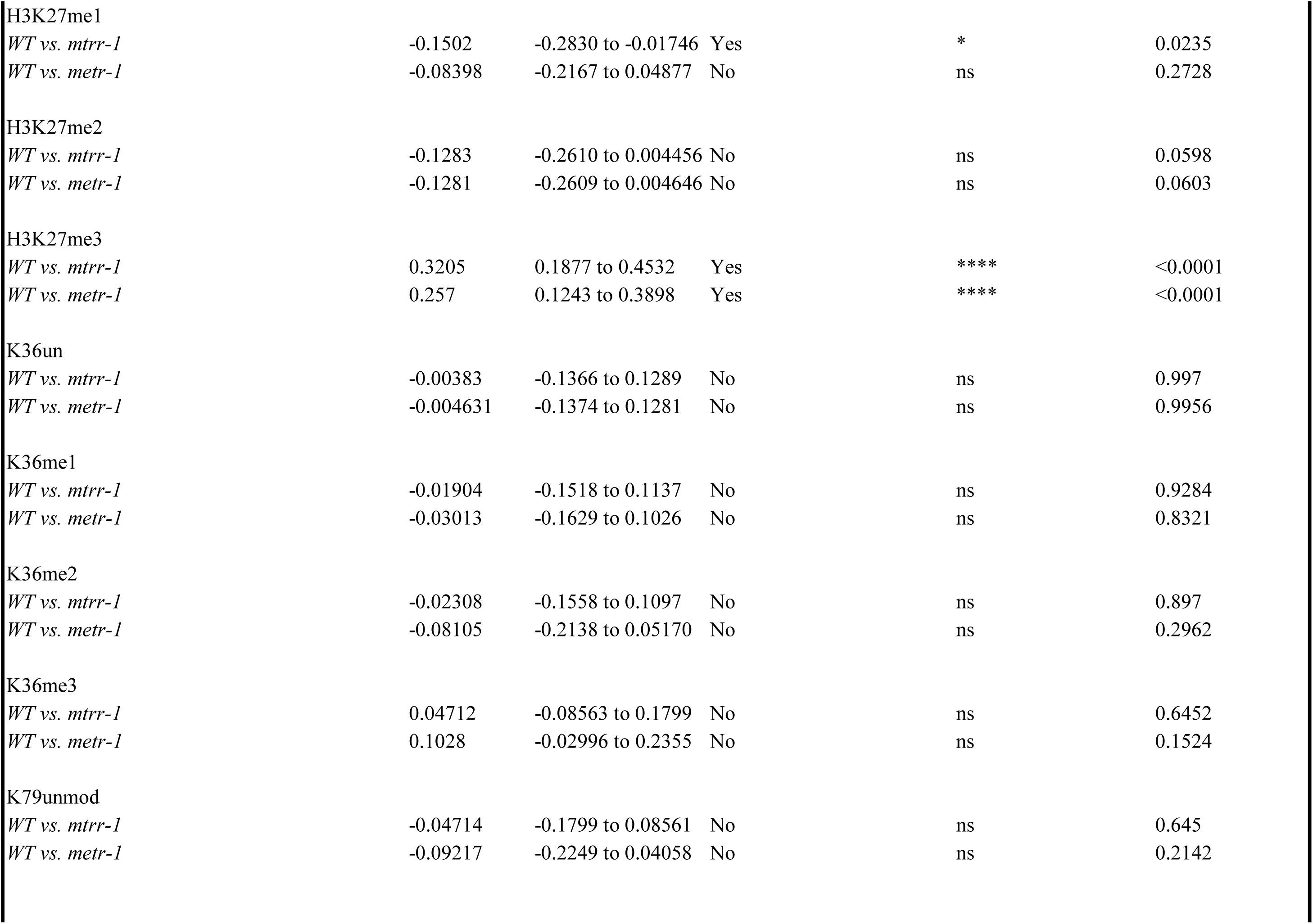
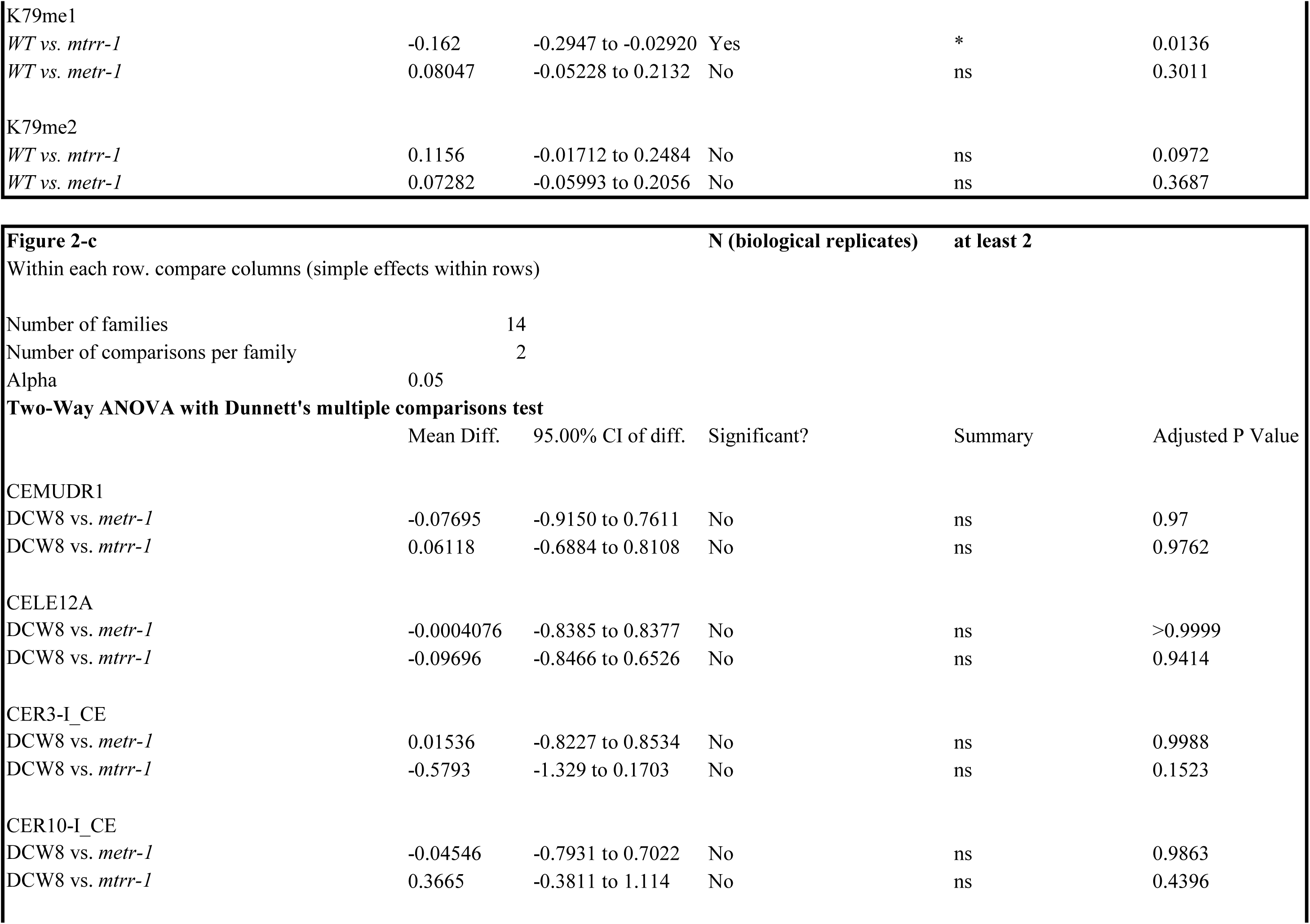
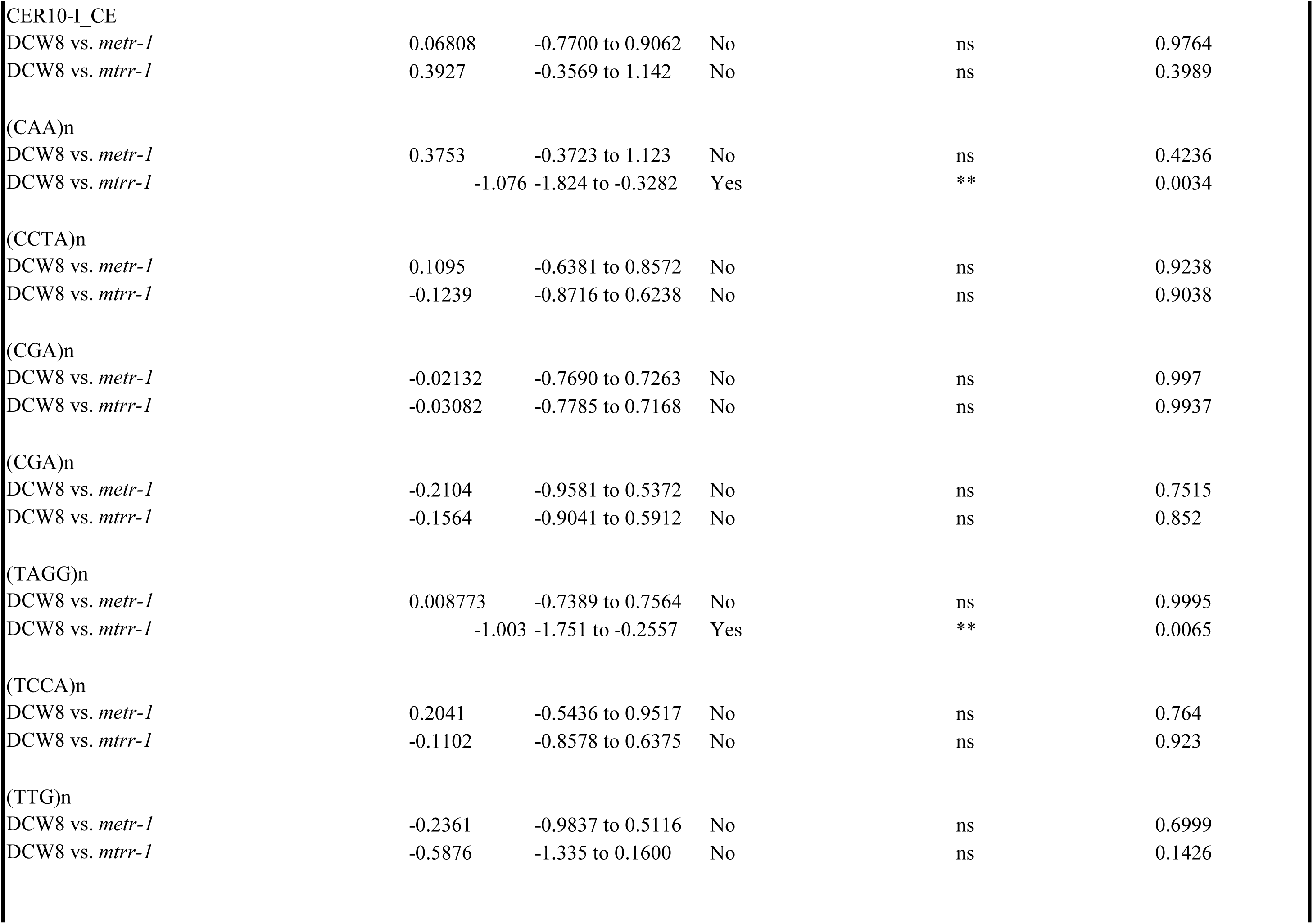
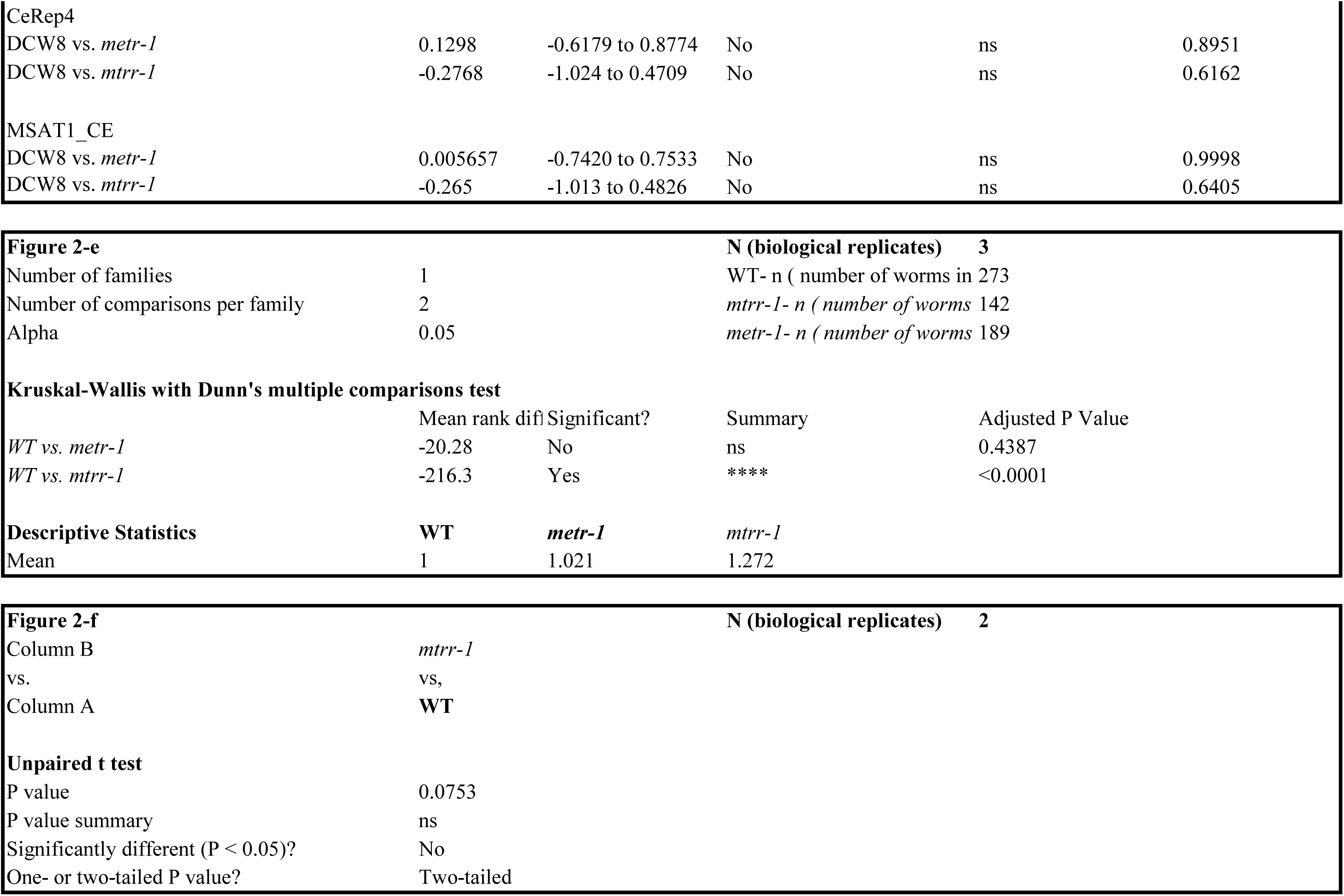
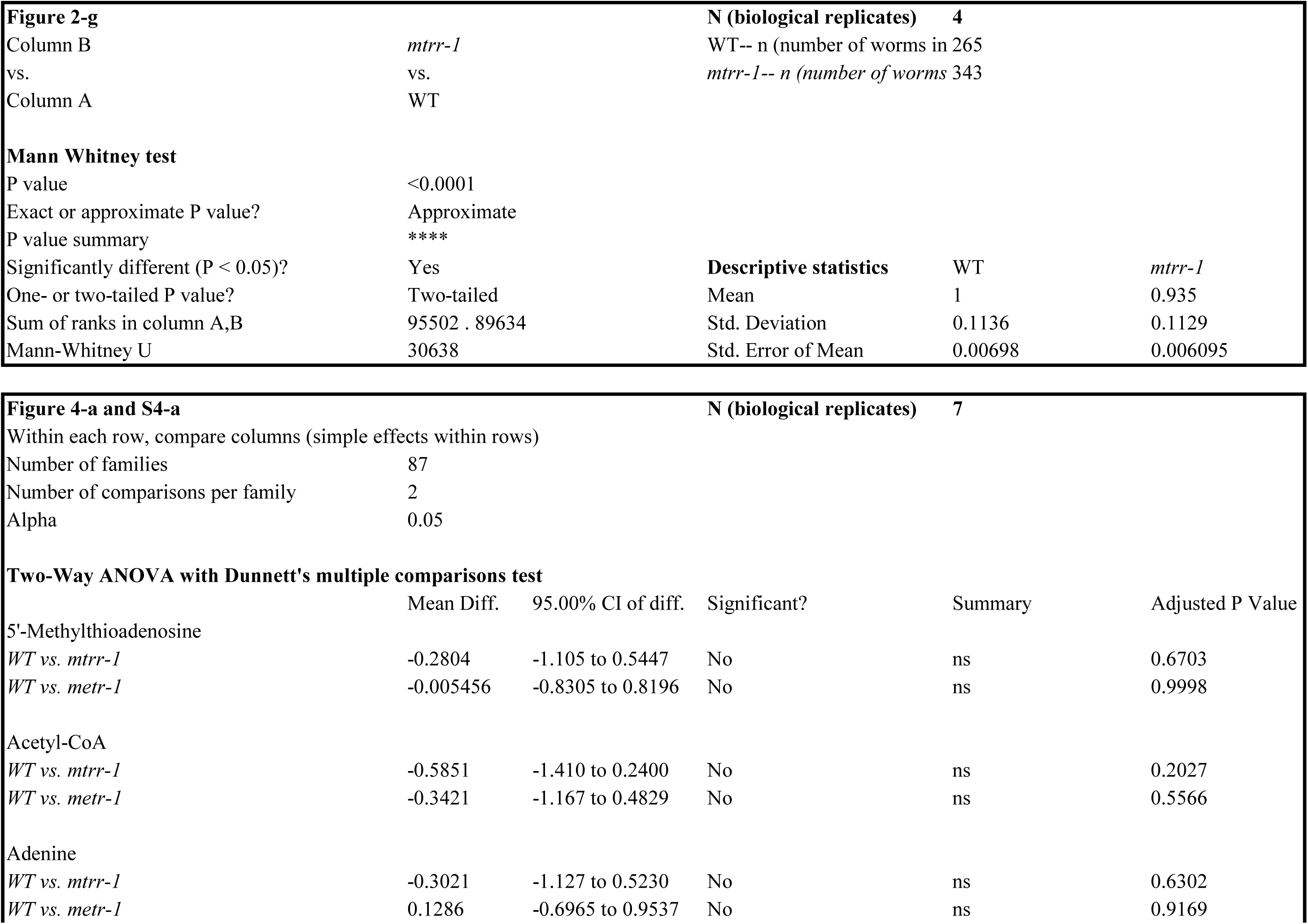
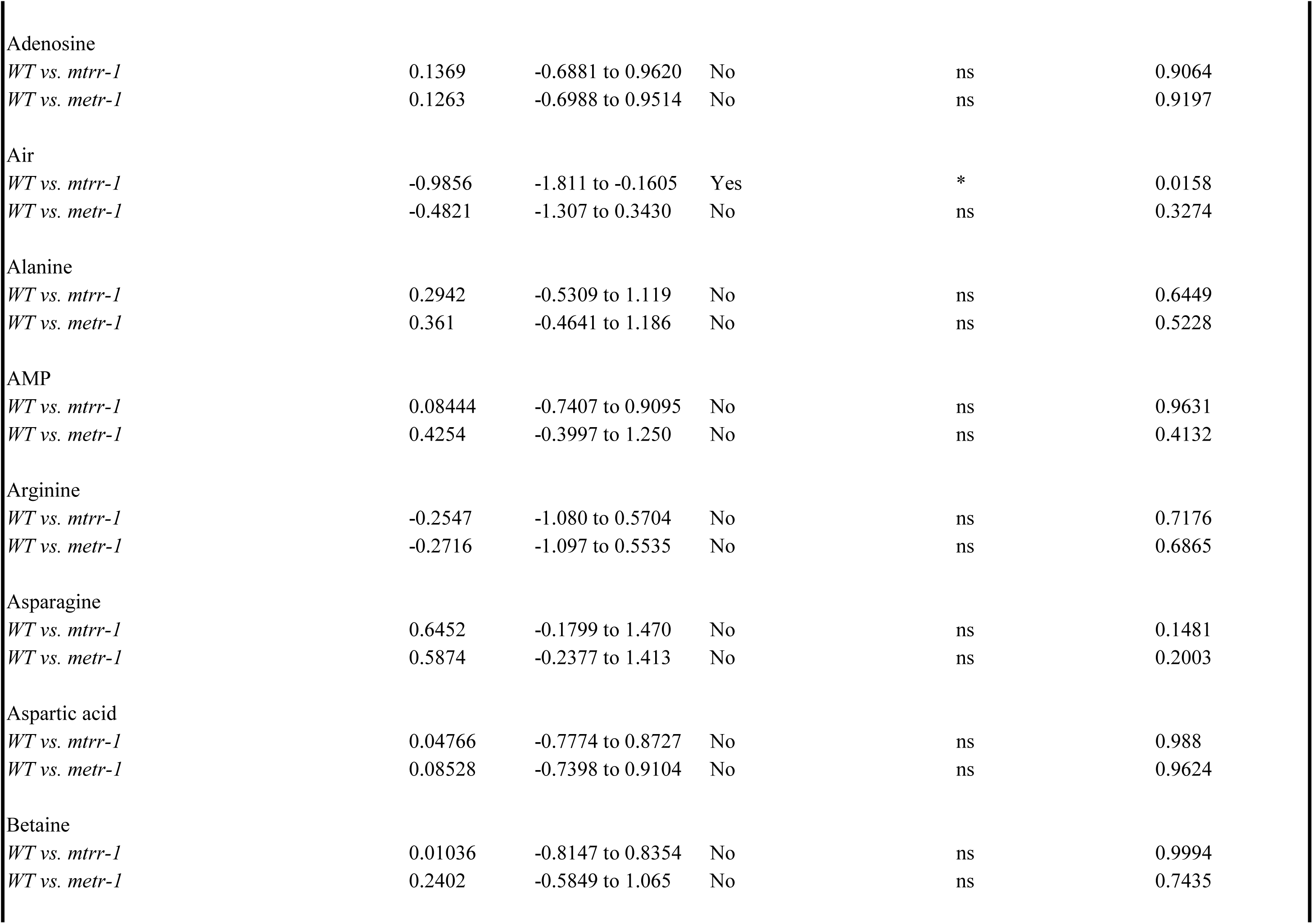
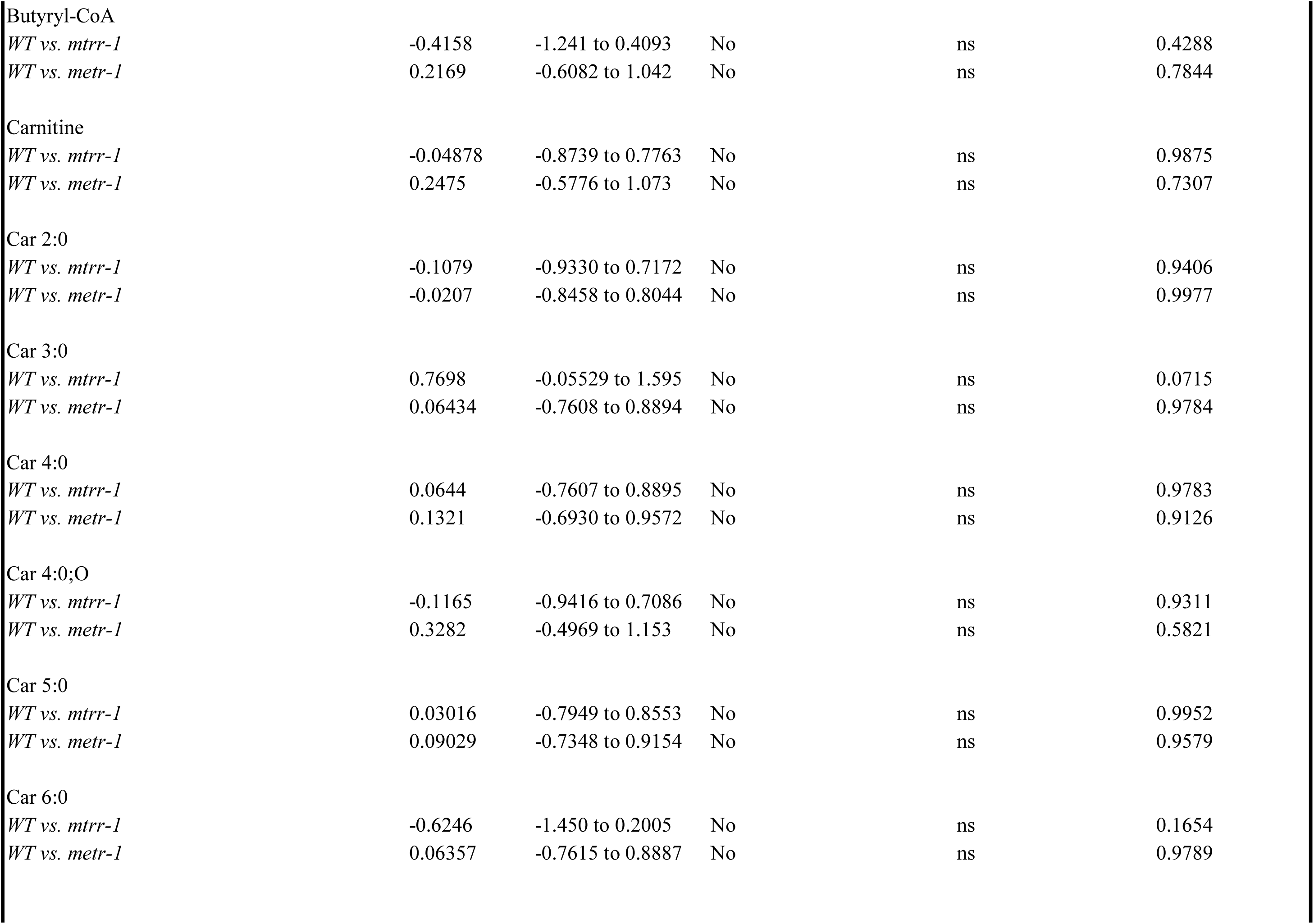
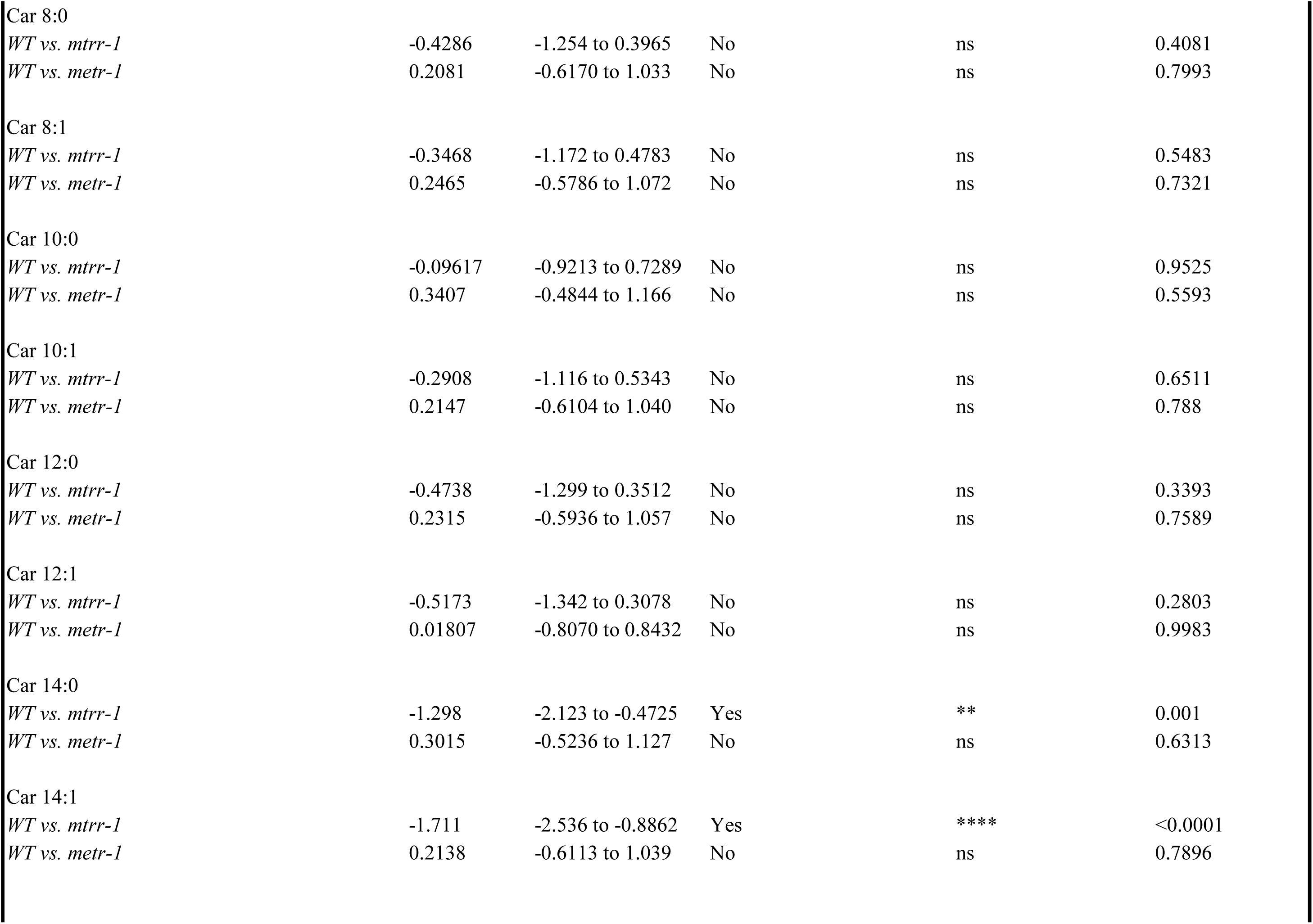
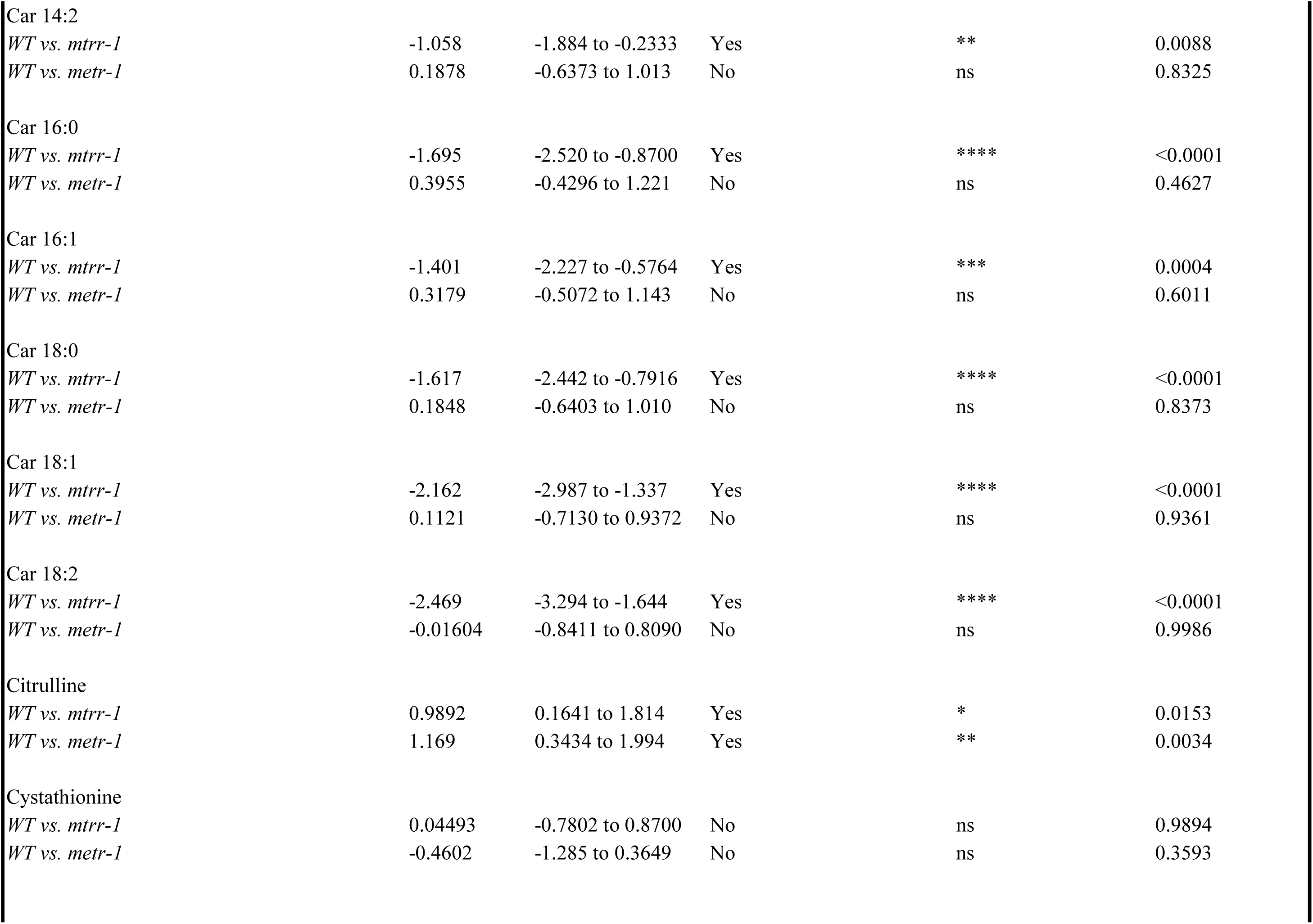
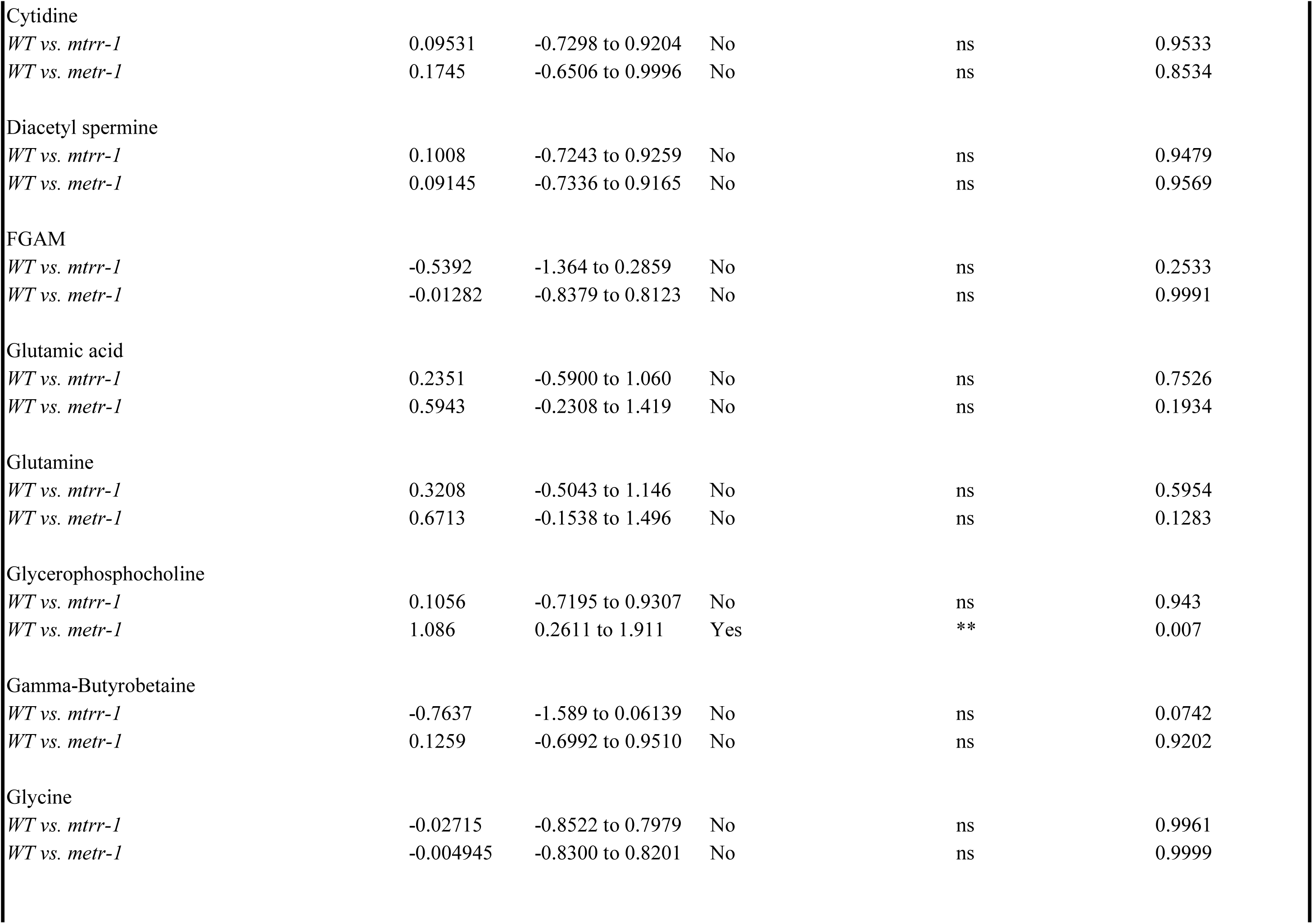
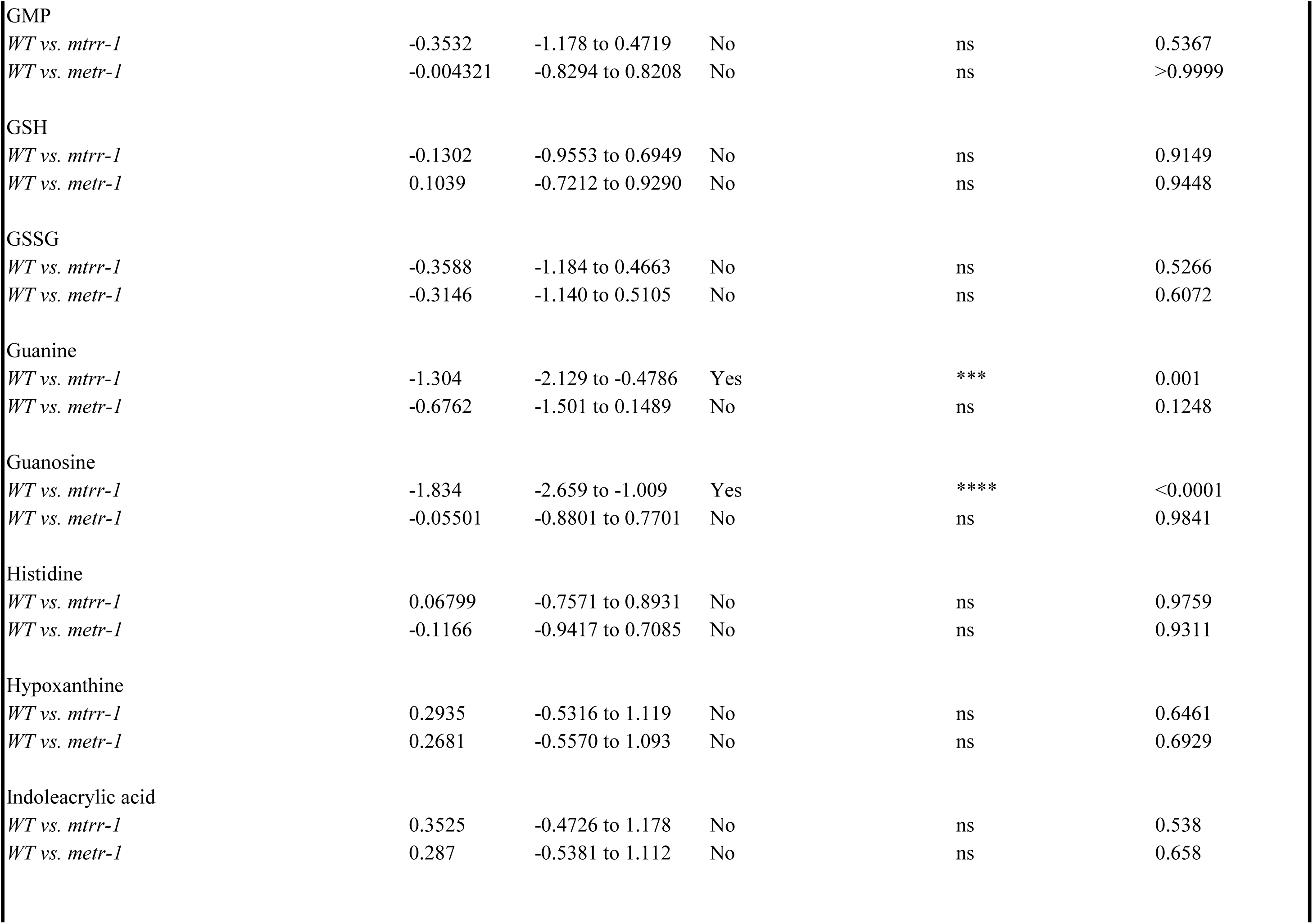
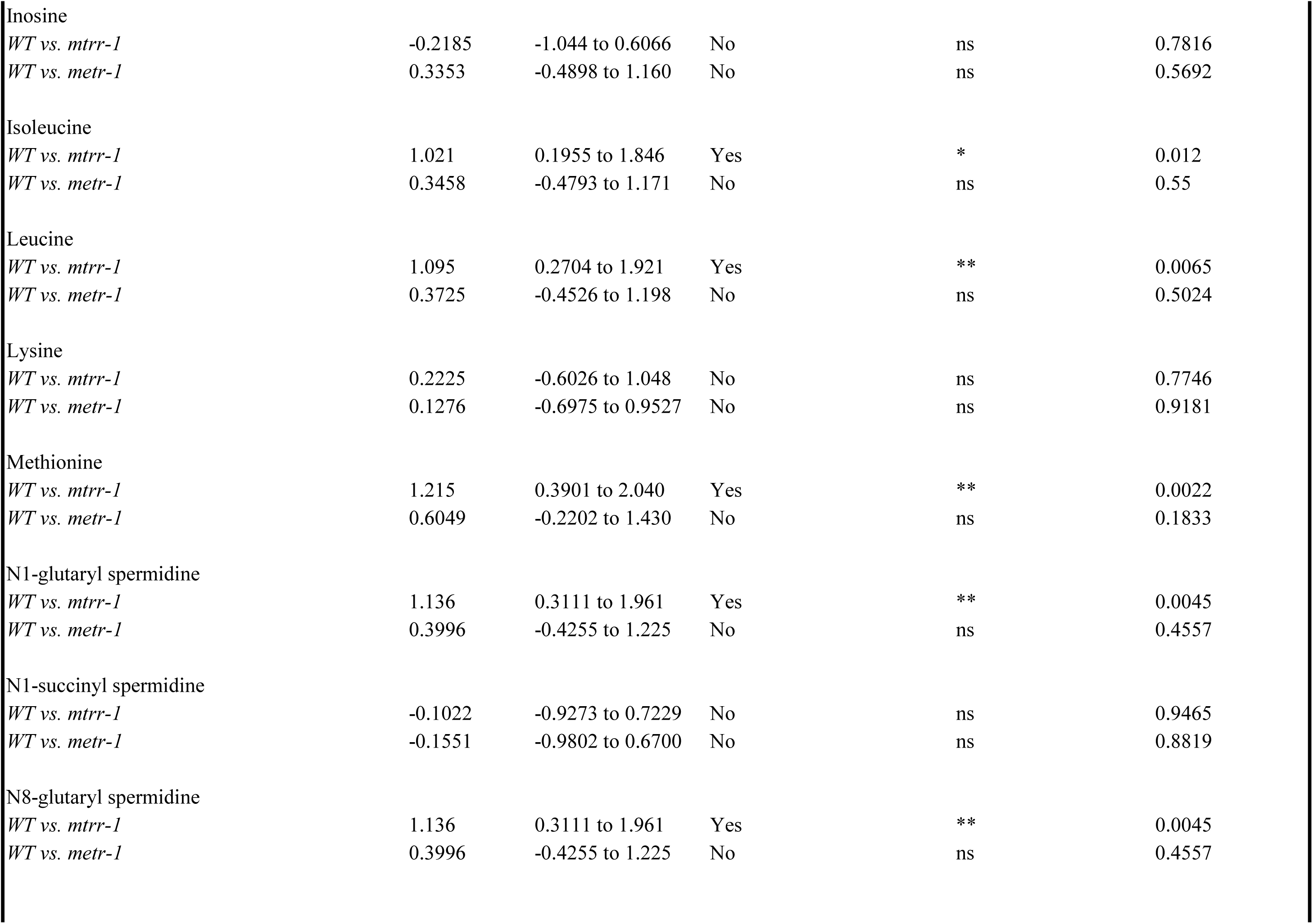
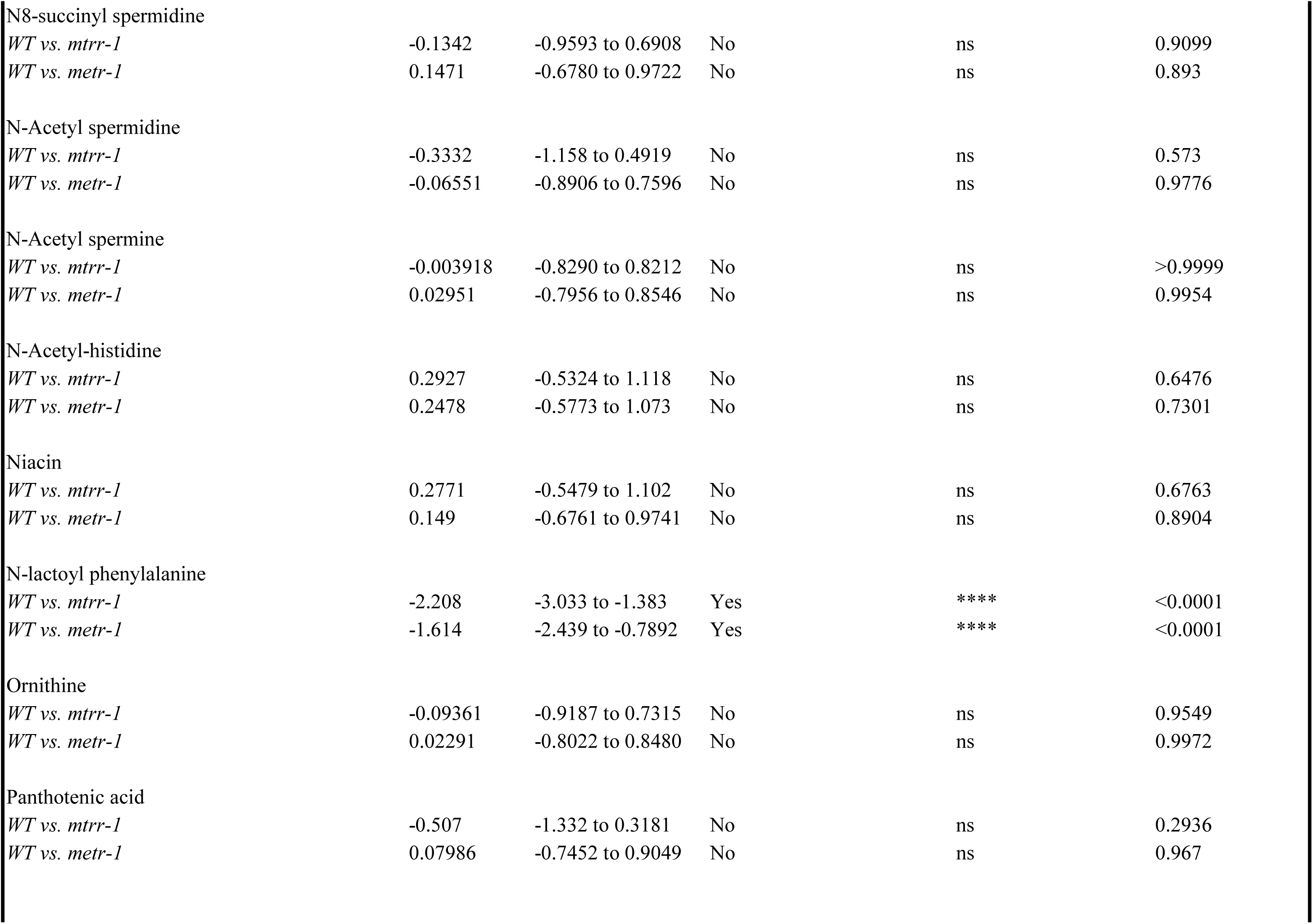
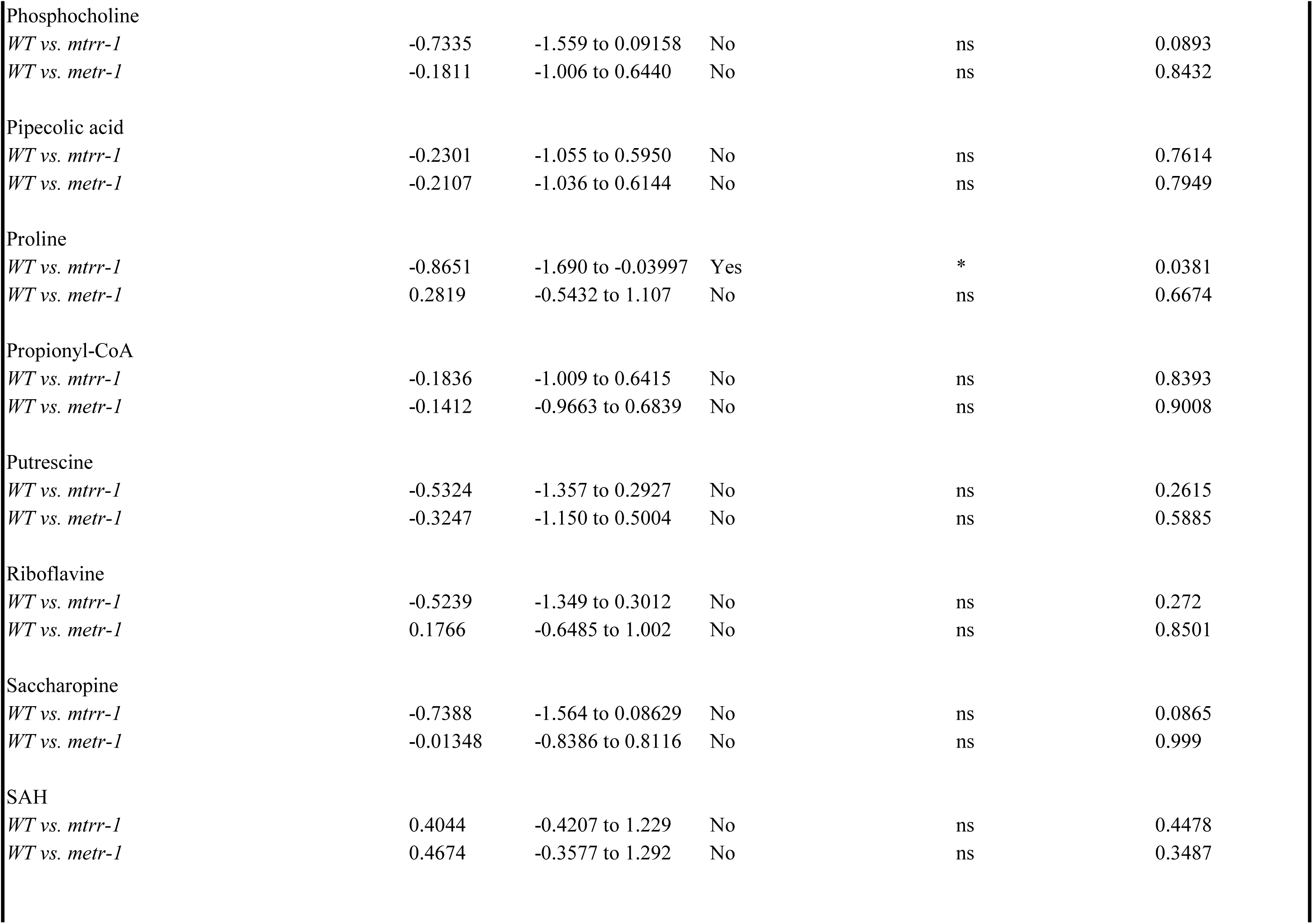
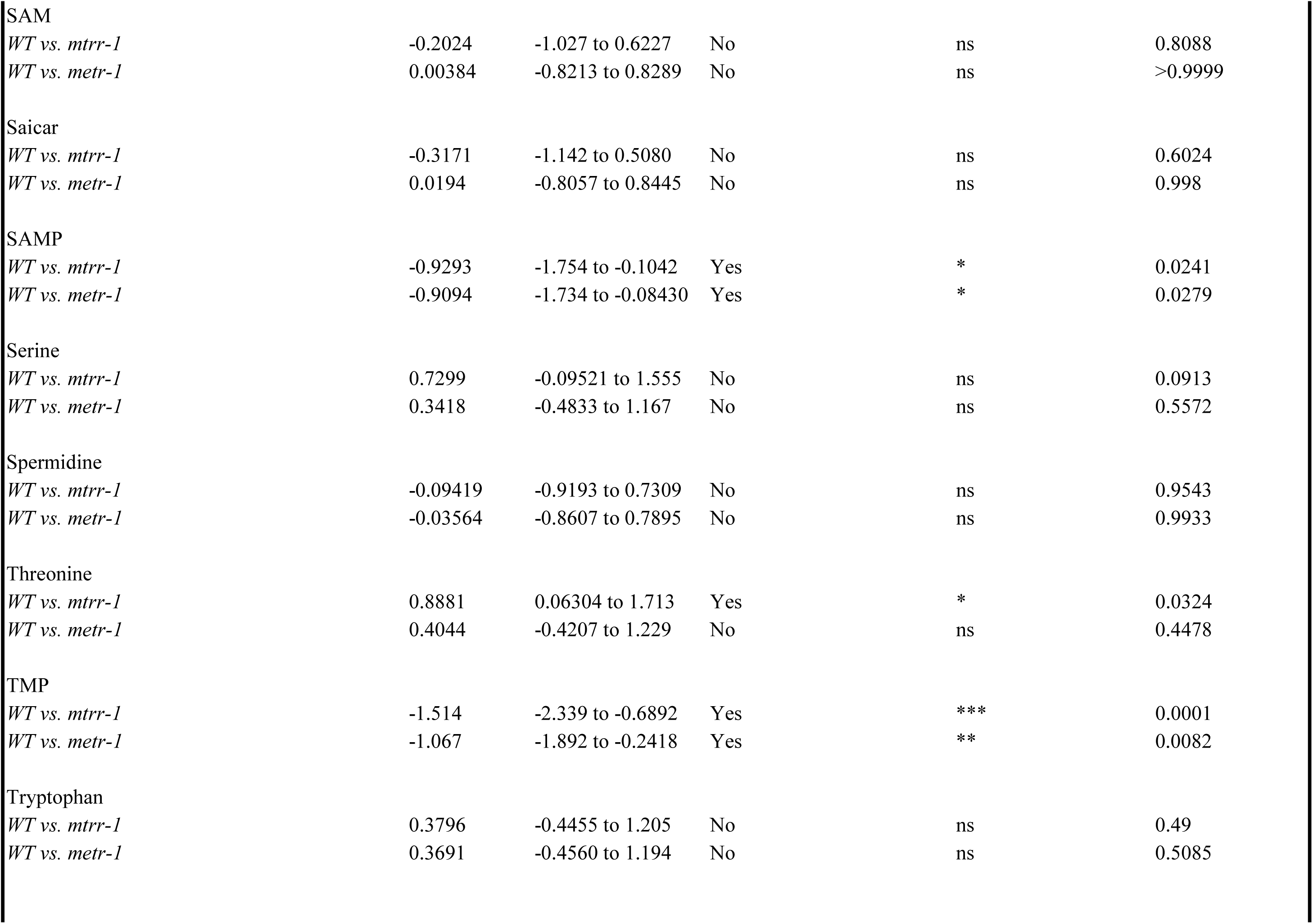
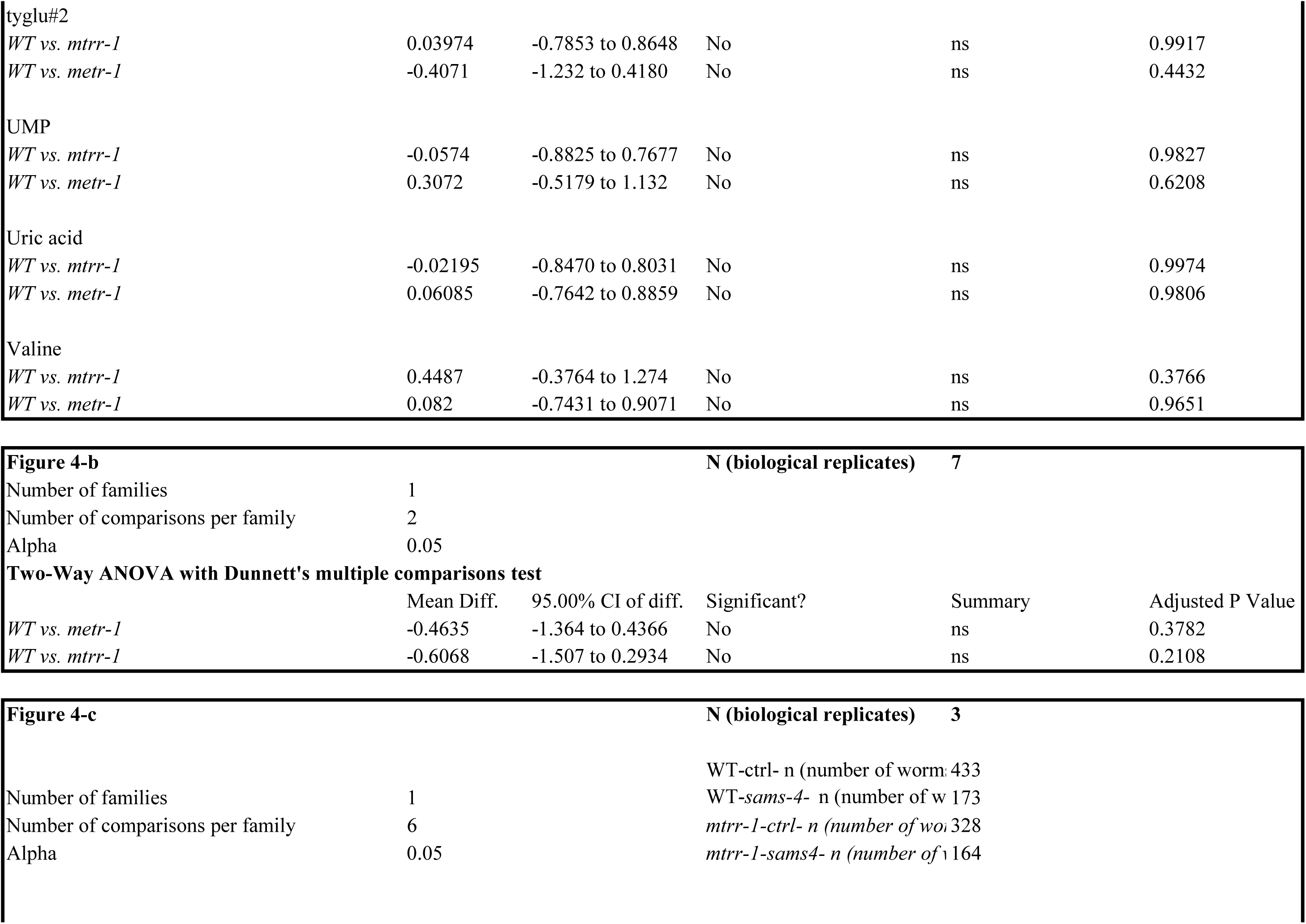
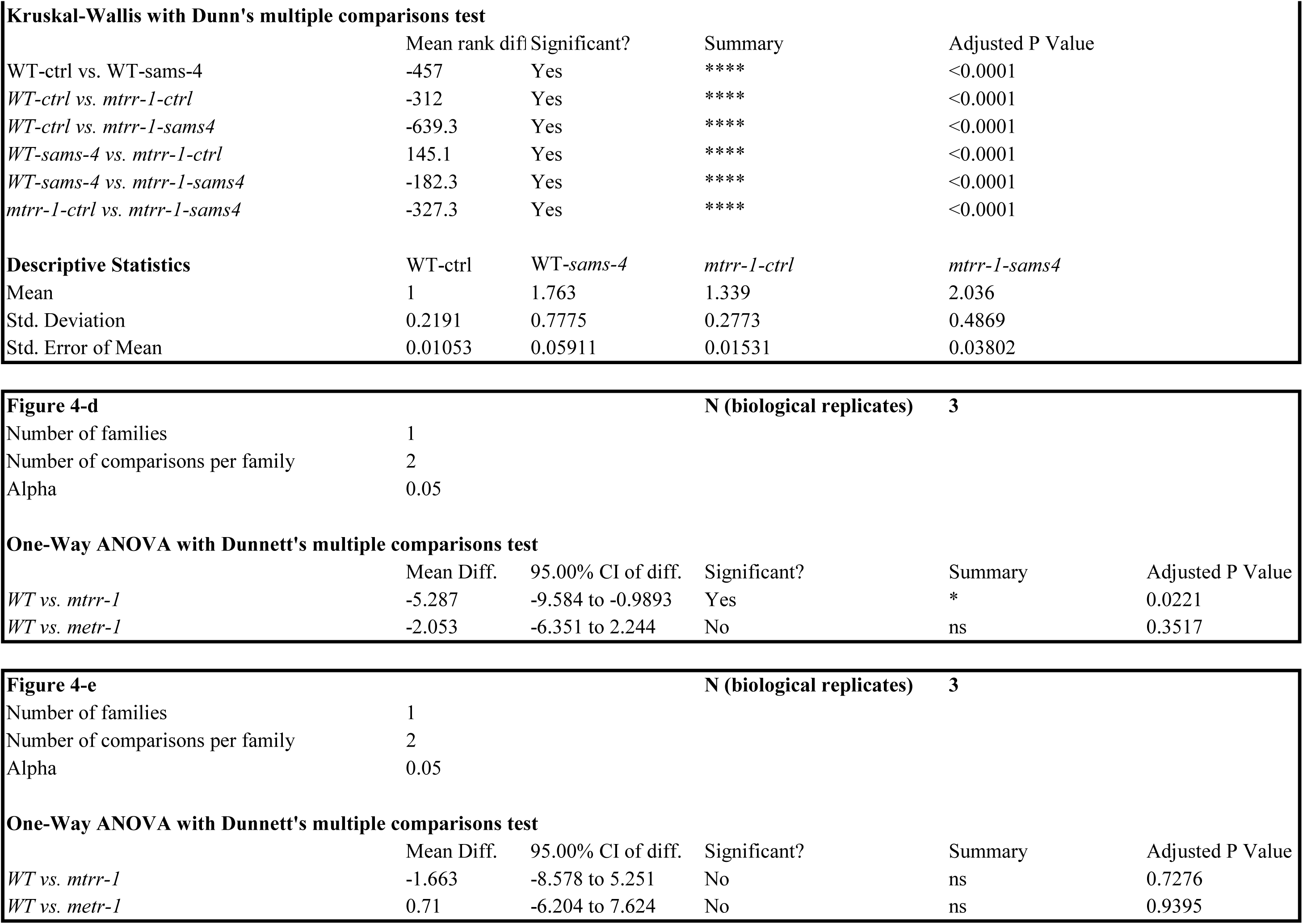
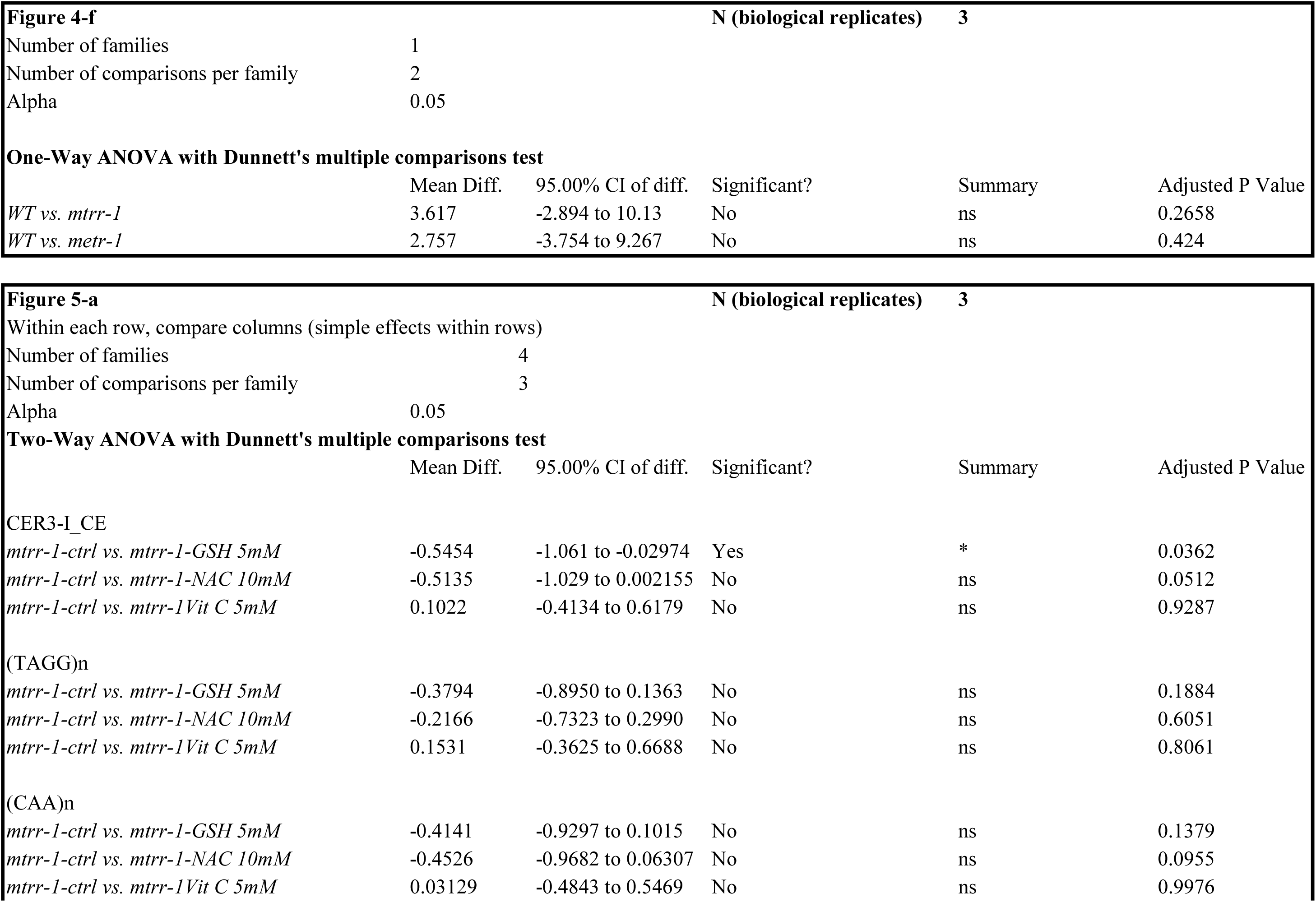
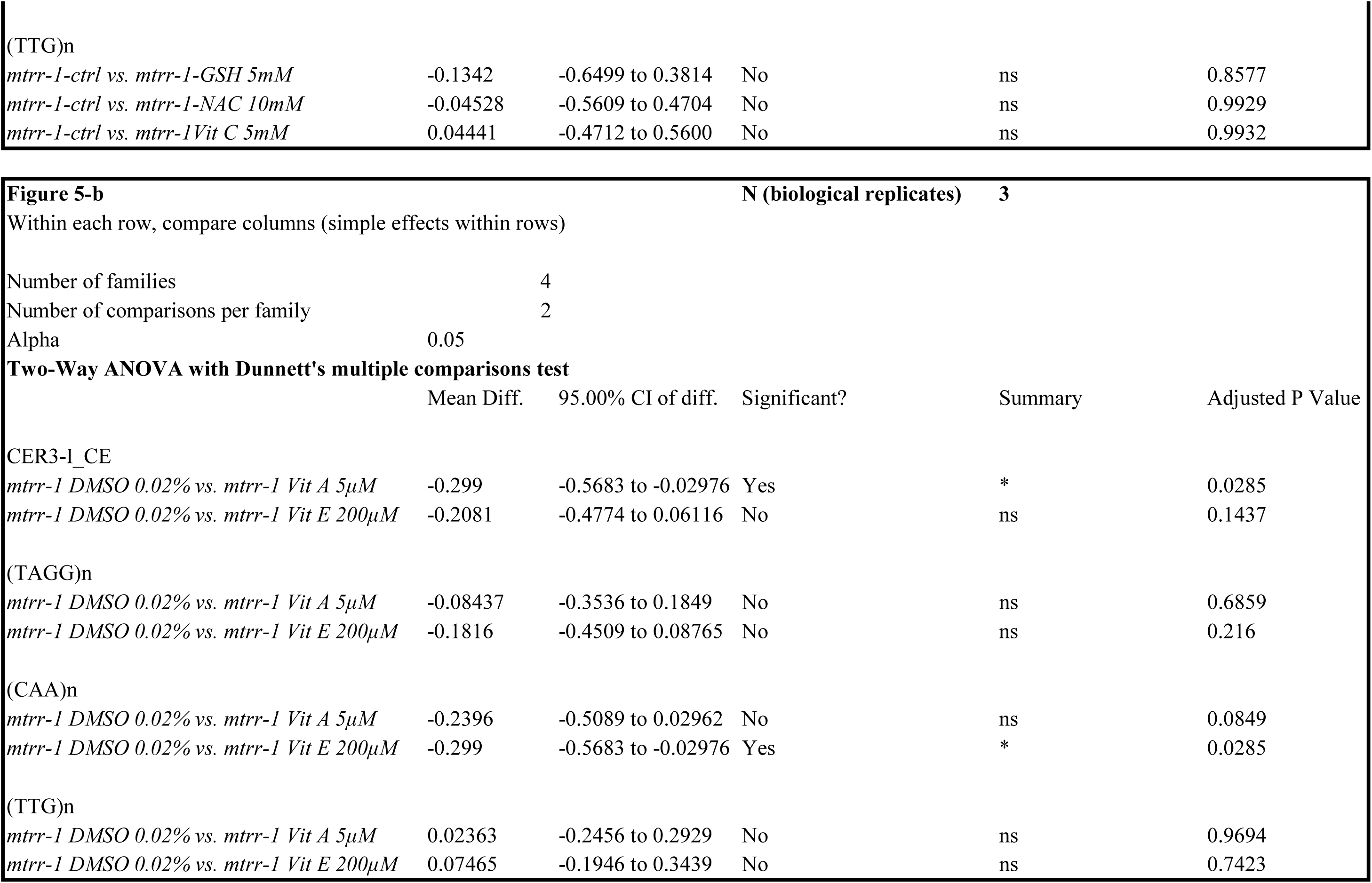
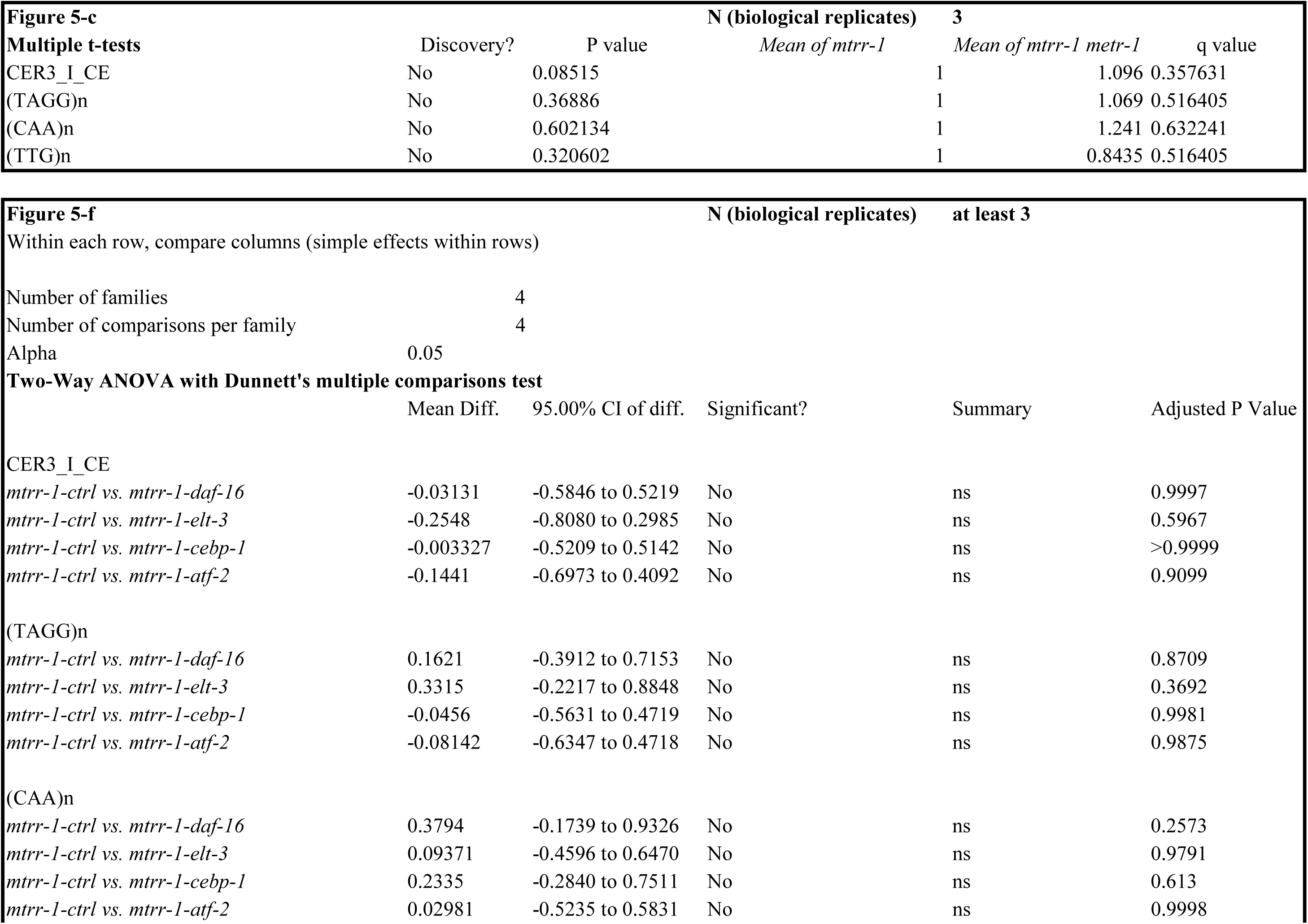
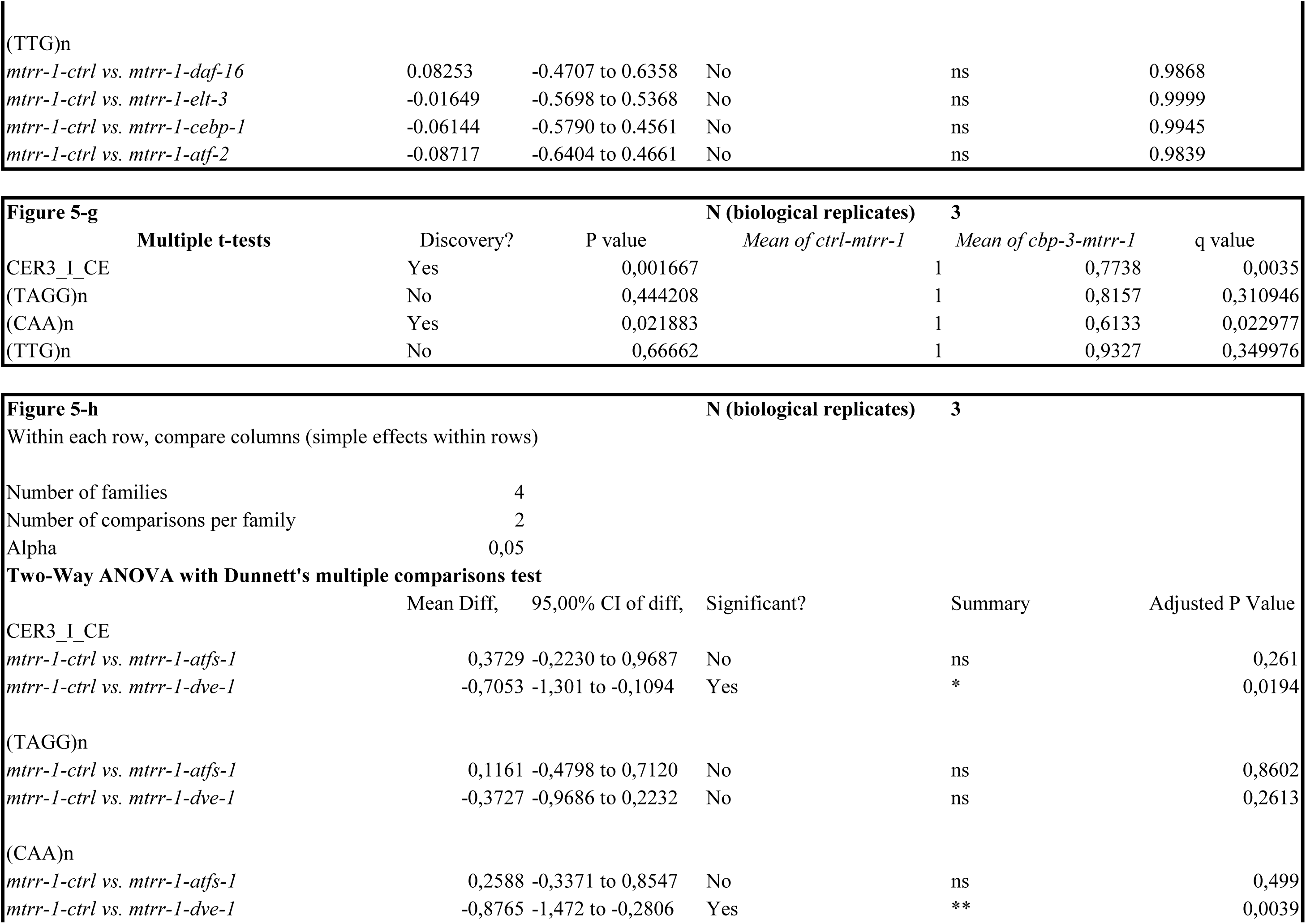
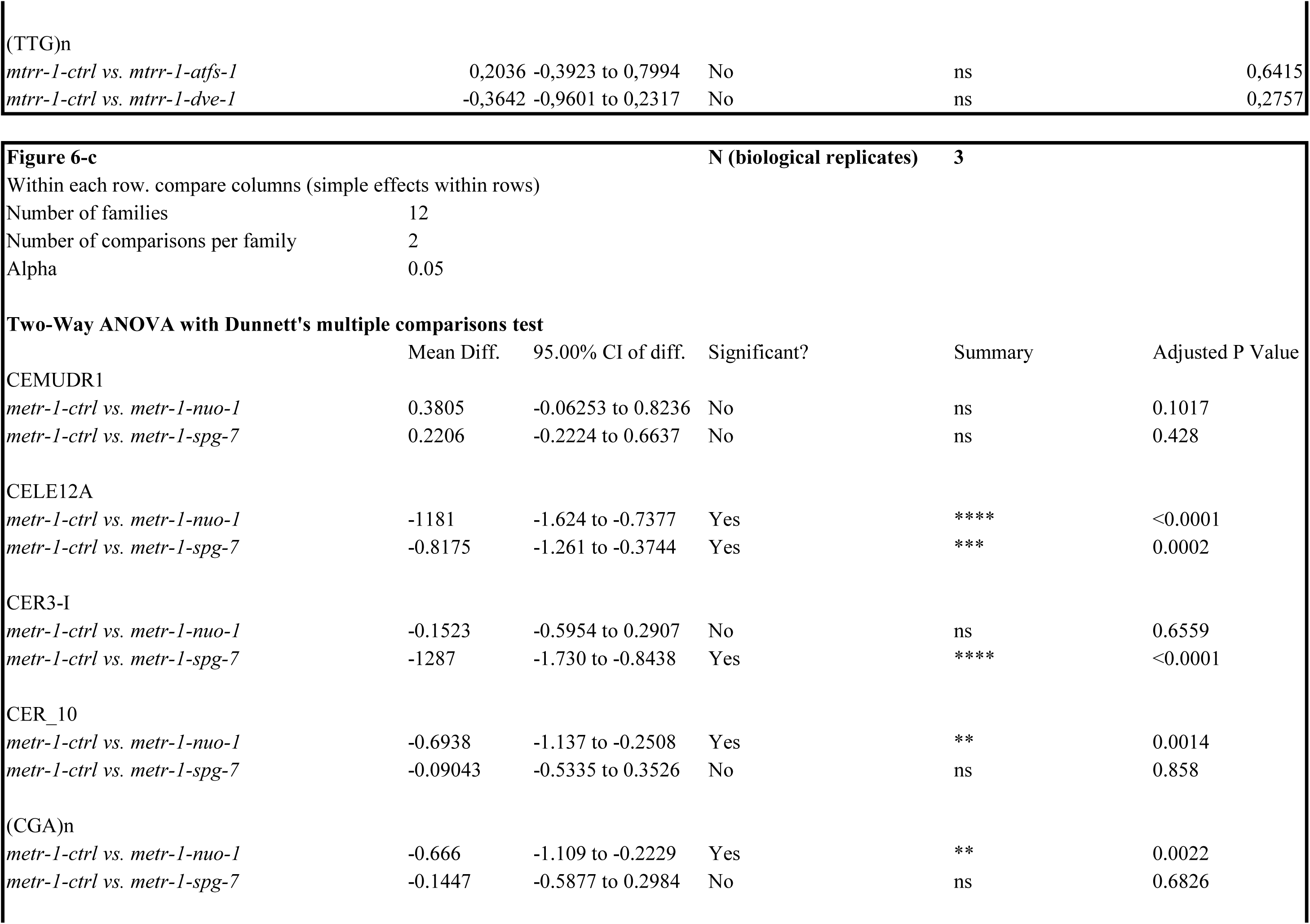
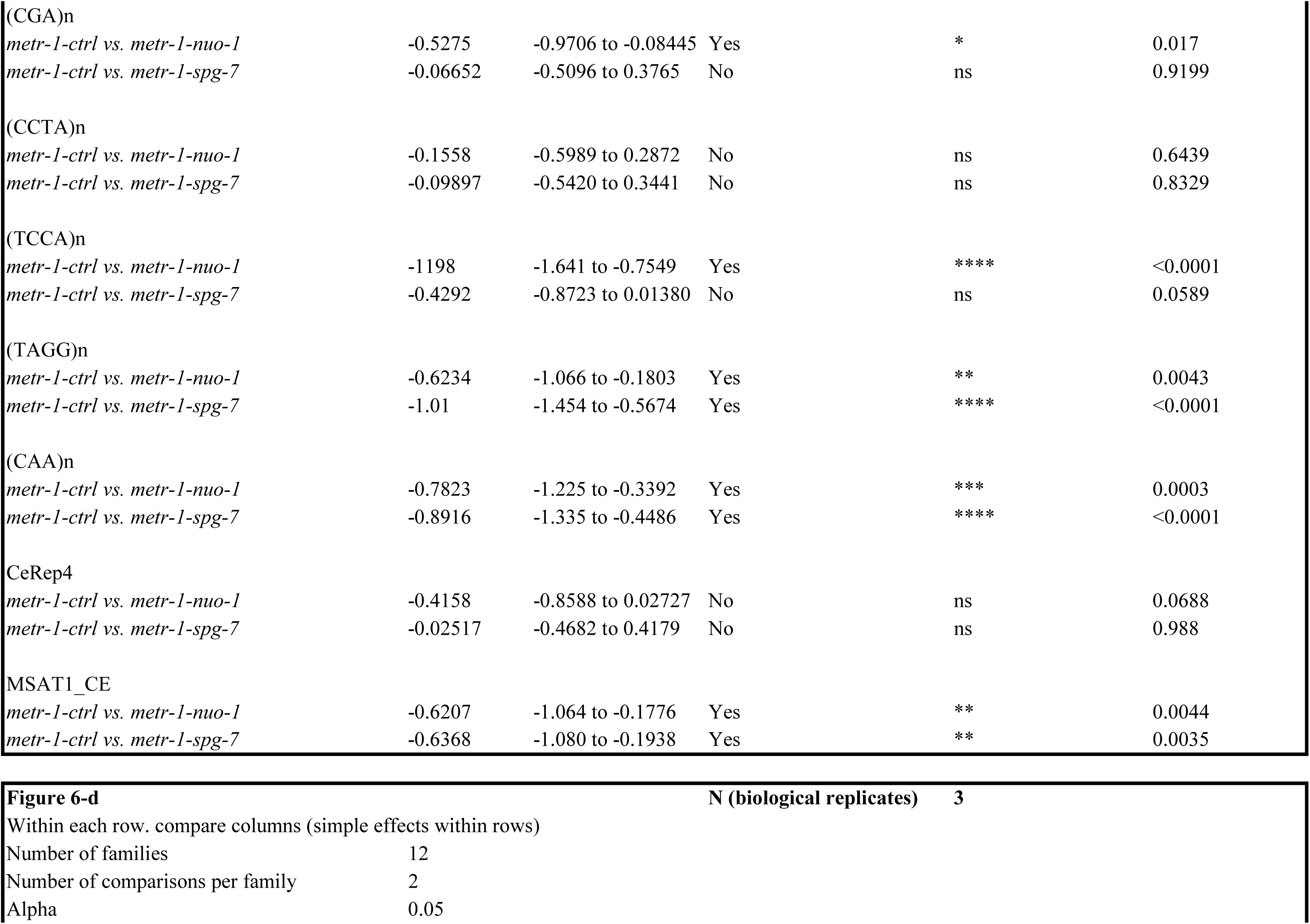
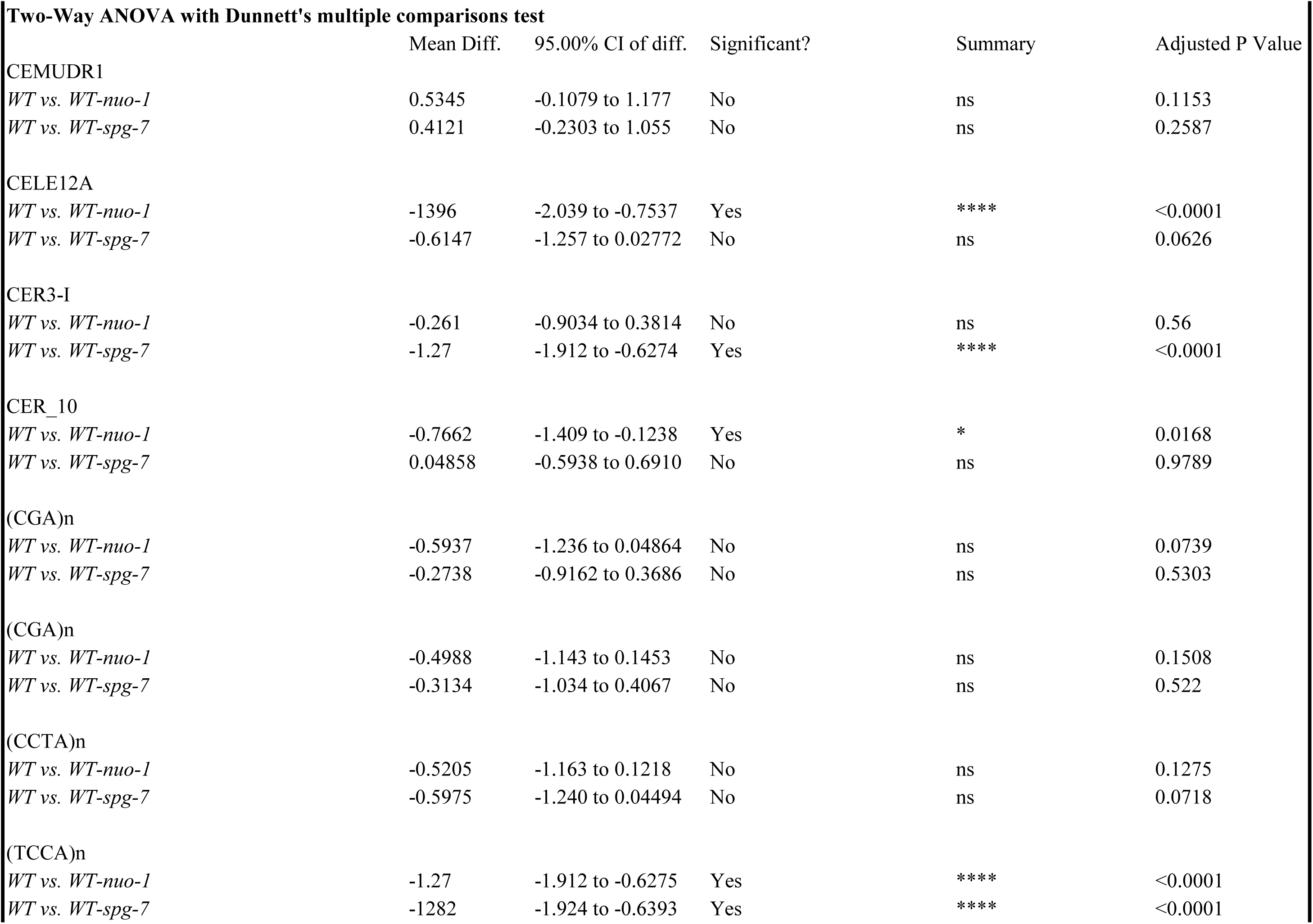
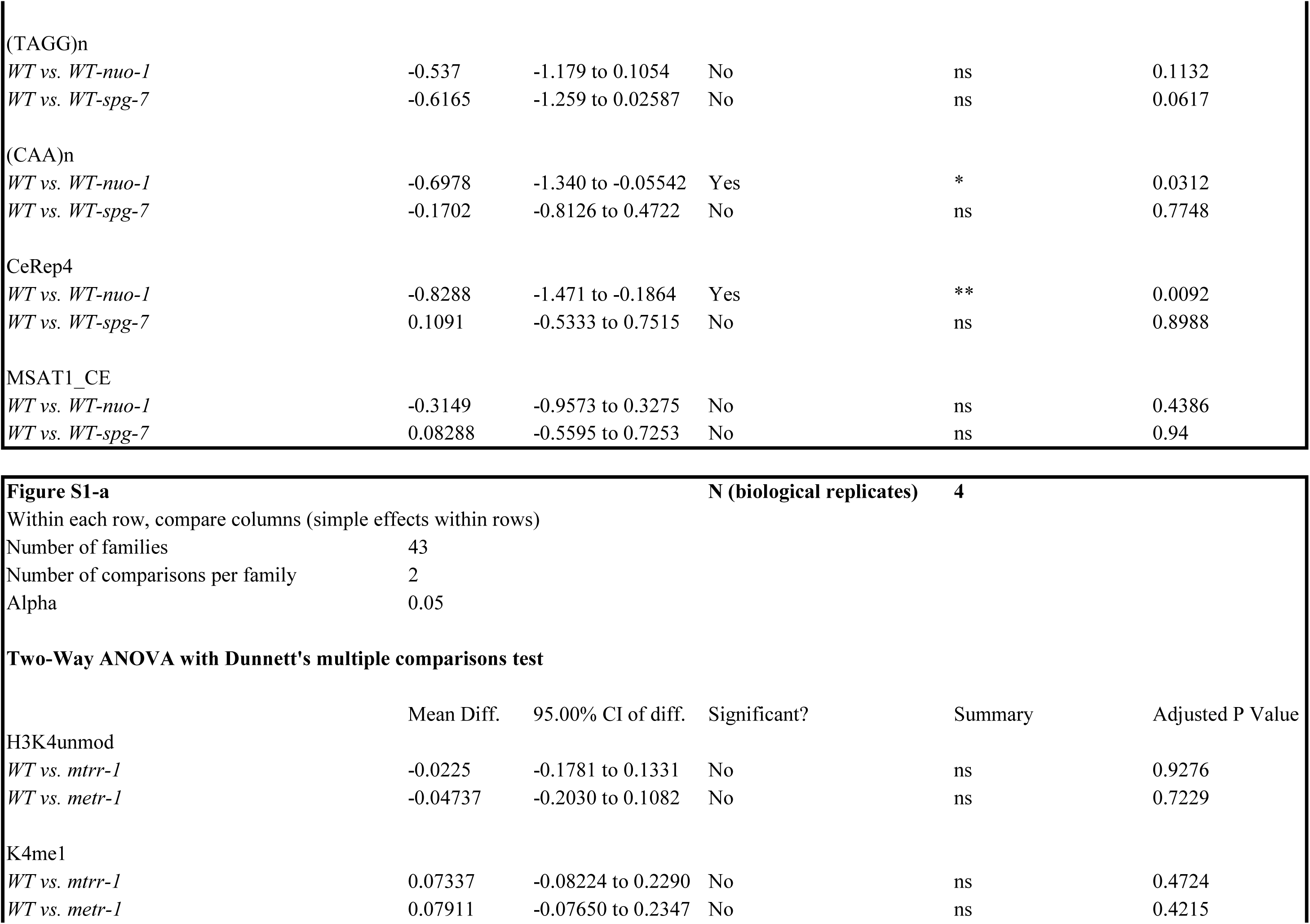
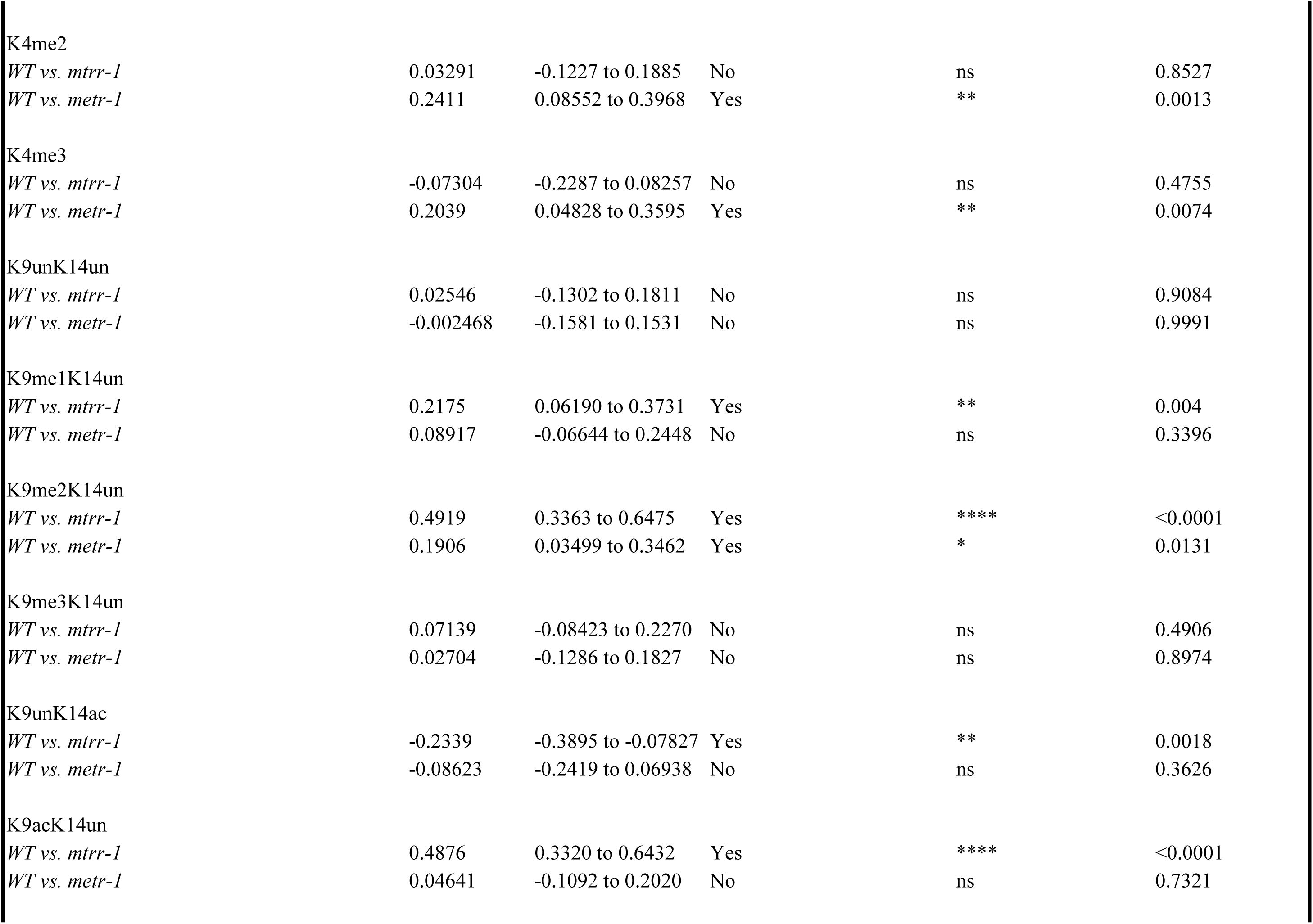
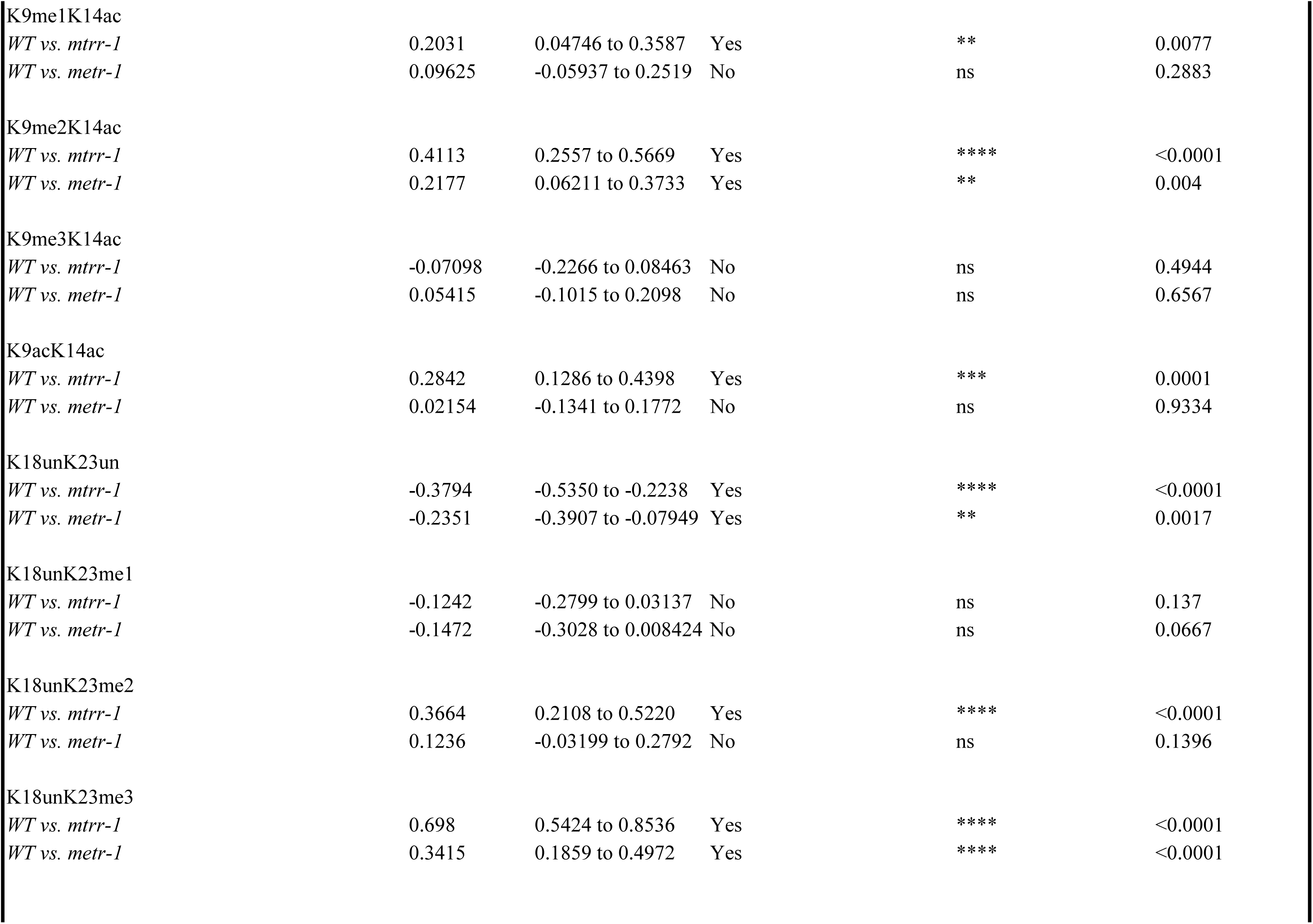
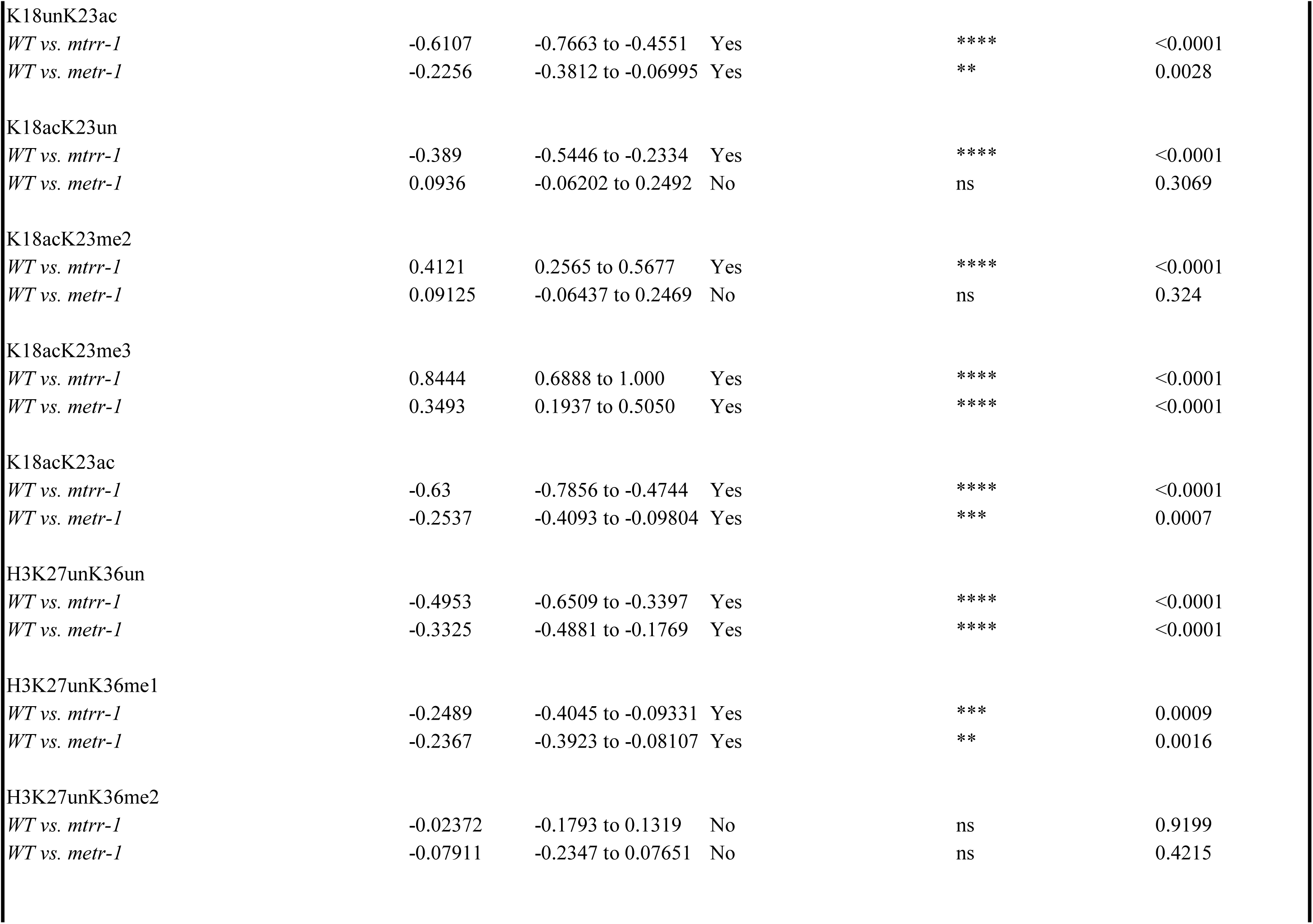
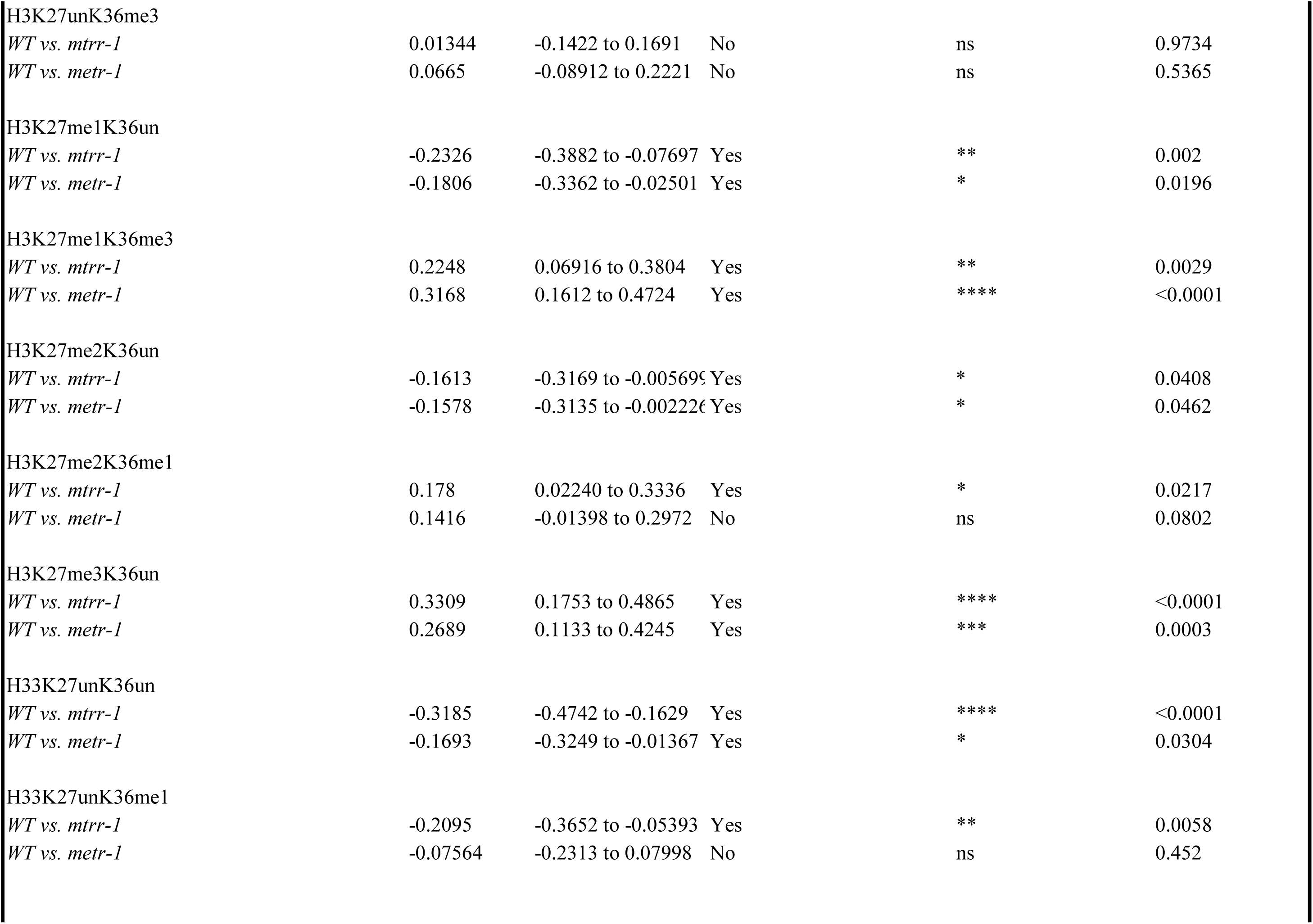
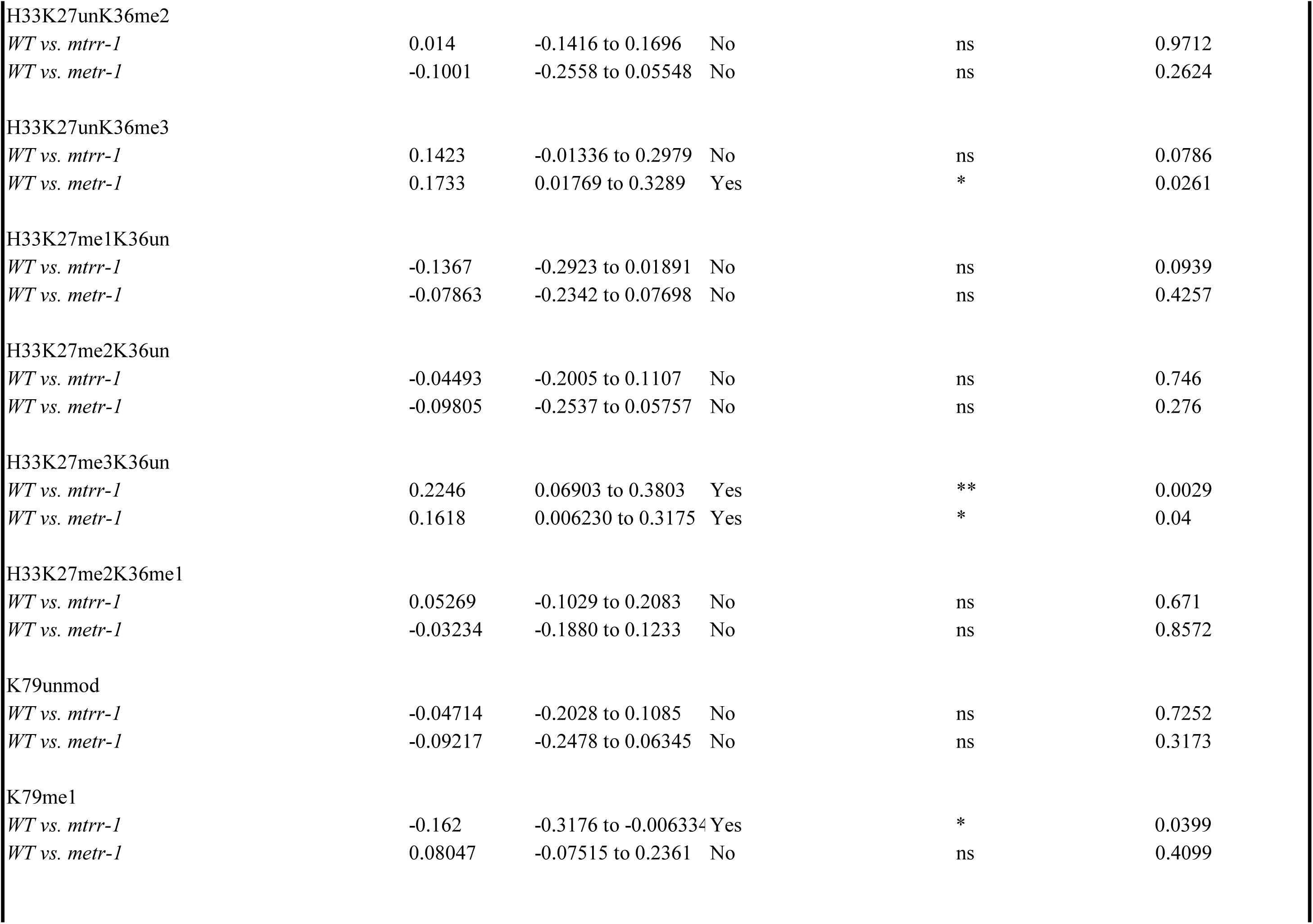
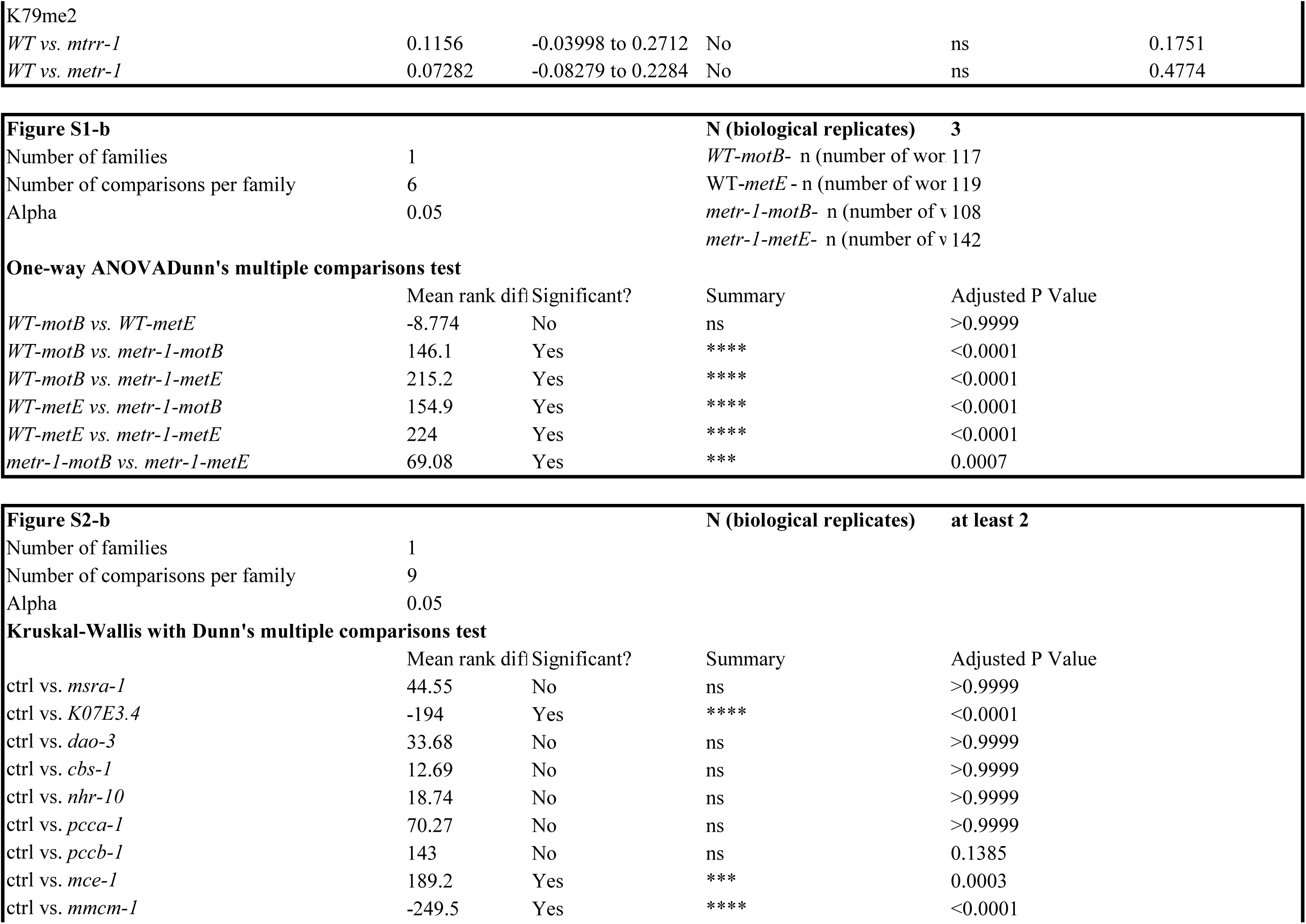
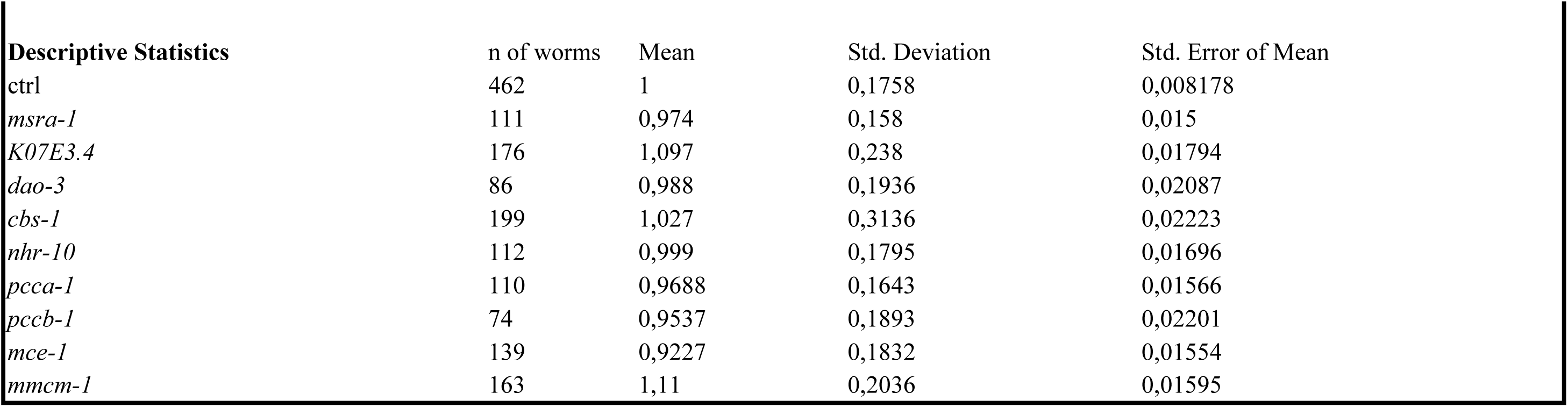
N and p-values of the presented experiments.

**Supplementary Table 3:**
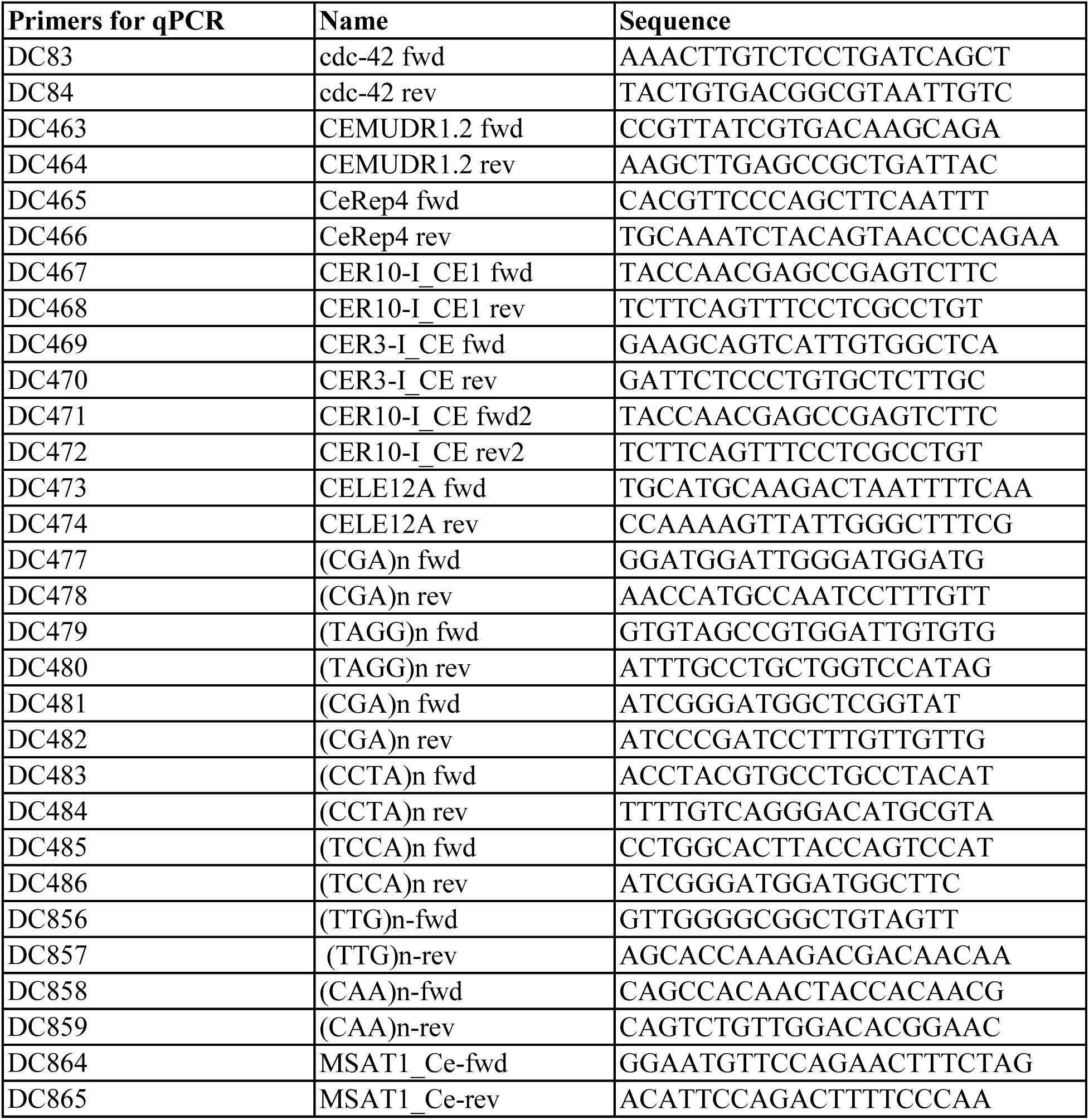
List of primers used in this study.

## References

1. Dai, Z., Ramesh, V. & Locasale, J.W. The evolving metabolic landscape of chromatin biology and epigenetics. Nat Rev Genet 21, 737–753 (2020).

2. Jambhekar, A., Dhall, A. & Shi, Y. Roles and regulation of histone methylation in animal development. Nat Rev Mol Cell Biol 20, 625–641 (2019).

3. Michalak, E.M., Burr, M.L., Bannister, A. J. & Dawson, M.A. The roles of DNA, RNA and histone methylation in ageing and cancer. Nat Rev Mol Cell Biol 20, 573–589 (2019).

4. Towbin, B.D. et al. Step-wise methylation of histone H3K9 positions heterochromatin at the nuclear periphery. Cell 150, 934–947 (2012).

5. Meng, J. et al. METHIONINE ADENOSYLTRANSFERASE4 Mediates DNA and Histone Methylation. Plant Physiol 177, 652–670 (2018).

6. Haws, S.A. et al. Methyl-Metabolite Depletion Elicits Adaptive Responses to Support Heterochromatin Stability and Epigenetic Persistence. Mol Cell 78, 210–223 e218 (2020).

7. Godbole, A.A. et al. S-adenosylmethionine synthases specify distinct H3K4me3 populations and gene expression patterns during heat stress. Elife 12 (2023).

8. Li, W., Han, Y., Tao, F. & Chong, K. Knockdown of SAMS genes encoding S-adenosyl-1- methionine synthetases causes methylation alterations of DNAs and histones and leads to late flowering in rice. J Plant Physiol 168, 1837–1843 (2011).

9. Ducker, G.S. & Rabinowitz, J.D. One-Carbon Metabolism in Health and Disease. Cell Metab 25, 27–42 (2017).

10. Sadhu, M.J. et al. Nutritional control of epigenetic processes in yeast and human cells. Genetics 195, 831–844 (2013).

11. Shiraki, N. et al. Methionine metabolism regulates maintenance and differentiation of human pluripotent stem cells. Cell Metab 19, 780–794 (2014).

12. Mentch, S.J. et al. Histone Methylation Dynamics and Gene Regulation Occur through the Sensing of One-Carbon Metabolism. Cell Metab 22, 861–873 (2015).

13. Shane, B. & Stokstad, E.L. Vitamin B12-folate interrelationships. Annu Rev Nutr 5, 115–141 (1985).

14. Leclerc, D. et al. Cloning and mapping of a cDNA for methionine synthase reductase, a flavoprotein defective in patients with homocystinuria. Proc Natl Acad Sci U S A 95, 3059–3064 (1998).

15. Lupu, D.S. et al. Altered methylation of specific DNA loci in the liver of Bhmt-null mice results in repression of Iqgap2 and F2rl2 and is associated with development of preneoplastic foci. FASEB J 31, 2090–2103 (2017).

16. Padmanabhan, N. et al. Mutation in folate metabolism causes epigenetic instability and transgenerational effects on development. Cell 155, 81–93 (2013).

17. Bertozzi, T.M. et al. Variably methylated retrotransposons are refractory to a range of environmental perturbations. Nat Genet 53,1233–1242 (2021).

18. Wasmuth, J., Schmid, R., Hedley, A. & Blaxter, M. On the extent and origins of genic novelty in the phylum Nematoda. PLoS Negl Trop Dis 2, e258 (2008).

19. Simpson, V.J., Johnson, T.E. & Hammen, R.F. Caenorhabditis elegans DNA does not contain 5-methylcytosine at any time during development or aging. Nucleic Acids Res 14, 6711–6719 (1986).

20. Wolthers, K.R. & Scrutton, N.S. Cobalamin uptake and reactivation occurs through specific protein interactions in the methionine synthase-methionine synthase reductase complex. FEBS J276, 1942–1951 (2009).

21. Olteanu, H. & Banerjee, R. Human methionine synthase reductase, a soluble P-450 reductase-like dual flavoprotein, is sufficient for NADPH-dependent methionine synthase activation. J Biol Chem 276, 35558–35563 (2001).

22. Yamada, K., Gravel, R.A., Toraya, T. & Matthews, R.G. Human methionine synthase reductase is a molecular chaperone for human methionine synthase. Proc Natl Acad Sci U S A 103, 9476–9481 (2006).

23. Sidoli, S., Vandamme, J., Salcini, A.E. & Jensen, O.N. Dynamic changes of histone H3 marks during Caenorhabditis elegans lifecycle revealed by middle-down proteomics. Proteomics 16, 459–464 (2016).

24. Solovei, I., Thanisch, K. & Feodorova, Y. How to rule the nucleus: divide et impera. Curr Opin Cell Biol 40, 47–59 (2016).

25. Vandamme, J. et al. H3K23me2 is a new heterochromatic mark in Caenorhabditis elegans. Nucleic Acids Res 43, 9694–9710 (2015).

26. Schwartz-Orbach, L. et al. Caenorhabditis elegans nuclear RNAi factor SET-32 deposits the transgenerational histone modification, H3K23me3. Elife 9 (2020).

27. Papazyan, R. et al. Methylation of histone H3K23 blocks DNA damage in pericentric heterochromatin during meiosis. Elife 3, e02996 (2014).

28. Vinson, D.A. et al. De novo methylation of histone H3K23 by the methyltransferases EHMT1/GLP and EHMT2/G9a. Epigenetics Chromatin 15, 36 (2022).

29. Zecic, A., Dhondt, I. & Braeckman, B.P. The nutritional requirements of Caenorhabditis elegans. Genes Nutr 14, 15 (2019).

30. Townsend, D.M., Tew, K.D. & Tapiero, H. Sulfur containing amino acids and human disease. Biomed Pharmacother 58, 47–55 (2004).

31. Padeken, J., Methot, S.P. & Gasser, S.M. Establishment of H3K9-methylated heterochromatin and its functions in tissue differentiation and maintenance. Nat Rev Mol Cell Biol 23, 623–640 (2022).

32. Jin, Y., Tam, O.H., Paniagua, E. & Hammell, M. TEtranscripts: a package for including transposable elements in differential expression analysis of RNA-seq datasets. Bioinformatics 31, 3593–3599 (2015).

33. Cabreiro, F. et al. Metformin retards aging in C. elegans by altering microbial folate and methionine metabolism. Cell 153, 228–239 (2013).

34. Liu, Y.J. et al. Glycine promotes longevity in Caenorhabditis elegans in a methionine cycle- dependent fashion. PLoS Genet 15, el007633 (2019).

35. Padeken, J. et al. Synergistic lethality between BRCA1 and H3K9me2 loss reflects satellite derepression. Genes Dev 33, 436–451 (2019).

36. Giese, G.E. et al. Caenorhabditis elegans methionine/S-adenosylmethionine cycle activity is sensed and adjusted by a nuclear hormone receptor. Elife 9 (2020).

37. Pellegrino, M.W. et al. Mitochondrial UPR-regulated innate immunity provides resistance to pathogen infection. Nature 516, 414–417 (2014).

38. Nargund, A.M., Pellegrino, M.W., Fiorese, C.J., Baker, B.M. & Haynes, C.M. Mitochondrial import efficiency of ATFS-1 regulates mitochondrial UPR activation. Science 337, 587–590 (2012).

39. Calfon, M. et al. IRE1 couples endoplasmic reticulum load to secretory capacity by processing the XBP-1 mRNA. Nature 415, 92–96 (2002).

40. Link, C.D., Cypser, J.R., Johnson, C.J. & Johnson, T.E. Direct observation of stress response in Caenorhabditis elegans using a reporter transgene. Cell Stress Chaperones 4, 235–242 (1999).

41. Bacaj, T. & Shaham, S. Temporal control of cell-specific transgene expression in Caenorhabditis elegans. Genetics 176, 2651–2655 (2007).

42. Munkacsy, E. et al. DLK-1, SEK-3 and PMK-3 Are Required for the Life Extension Induced by Mitochondrial Bioenergetic Disruption in C. elegans. PLoS Genet 12, el006133 (2016).

43. McGarry, J.D. & Brown, N.F. The mitochondrial carnitine palmitoyltransferase system. From concept to molecular analysis. Eur J Biochem 244, 1–14 (1997).

44. McCoin, C.S. et al. Unique plasma metabolomic signatures of individuals with inherited disorders of long-chain fatty acid oxidation. J Inherit Metab Dis 39, 399–408 (2016).

45. Mann, G., Mora, S., Madu, G. & Adegoke, O.A.J. Branched-chain Amino Acids: Catabolism in Skeletal Muscle and Implications for Muscle and Whole-body Metabolism. Front Physiol 12, 702826 (2021).

46. Zhang, B. et al. Acylspermidines are conserved mitochondrial sirtuin-dependent metabolites. Nat Chem Biol 20, 812–822 (2024).

47. Elmore, C.L. et al. Metabolic derangement of methionine and folate metabolism in mice deficient in methionine synthase reductase. Mol Genet Metab 91, 85–97 (2007).

48. Gaughan, D.J. et al. The methionine synthase reductase (MTRR) A66G polymorphism is a novel genetic determinant of plasma homocysteine concentrations. Atherosclerosis 157, 451–456 (2001).

49. Ferguson, G.D. & Bridge, W.J. The glutathione system and the related thiol network in Caenorhabditis elegans. Redox Biol 24, 101171 (2019).

50. Yang, W., Dierking, K. & Schulenburg, H. WormExp: a web-based application for a Caenorhabditis elegans-specific gene expression enrichment analysis. Bioinformatics 32, 943–945 (2016).

51. Heinz, S. et al. Simple combinations of lineage-determining transcription factors prime cis- regulatory elements required for macrophage and B cell identities. Mol Cell 38, 576–589 (2010).

52. Taouktsi, E. et al. Mitochondrial p38 Mitogen-Activated Protein Kinase: Insights into Its Regulation of and Role in LONP1-Deficient Nematodes. Int J Mol Sci 24 (2023).

53. Haynes, C.M., Petrova, K., Benedetti, C., Yang, Y. & Ron, D. ClpP mediates activation of a mitochondrial unfolded protein response in C. elegans. Dev Cell 13, 467–480 (2007).

54. Haynes, C.M., Yang, Y., Blais, S.P., Neubert, T.A. & Ron, D. The matrix peptide exporter HAF-1 signals a mitochondrial UPR by activating the transcription factor ZC376.7 in C. elegans. Mol Cell 37, 529–540 (2010).

55. Rea, S.L., Ventura, N. & Johnson, T.E. Relationship between mitochondrial electron transport chain dysfunction, development, and life extension in Caenorhabditis elegans. PLoS Biol 5, e259 (2007).

56. Grad, L.I. & Lemire, B.D. Mitochondrial complex I mutations in Caenorhabditis elegans produce cytochrome c oxidase deficiency, oxidative stress and vitamin-responsive lactic acidosis. Hum Mol Genet 13, 303–314 (2004).

57. Ye, C., Sutter, B.M., Wang, Y., Kuang, Z. & Tu, B.P. A Metabolic Function for Phospholipid and Histone Methylation. Mol Cell 66, 180–193 el 88 (2017).

58. Perez, M.F. & Sarkies, P. Histone methyltransferase activity affects metabolism in human cells independently of transcriptional regulation. PLoS Biol 21, e3002354 (2023).

59. Quiros, P.M. et al. Multi-omics analysis identifies ATF4 as a key regulator of the mitochondrial stress response in mammals. J Cell Biol 216, 2027–2045 (2017).

60. Hill, B.G. et al. Integration of cellular bioenergetics with mitochondrial quality control and autophagy. Biol Chem 393, 1485–1512 (2012).

61. Schafer, P, Muller, M., Kruger, A., Steinberg, C.E. & Menzel, R. Cytochrome P450- dependent metabolism of PCB52 in the nematode Caenorhabditis elegans. Arch Biochem Biophys 488, 60–68 (2009).

62. Larigot, L., Mansuy, D., Borowski, I., Coumoul, X. & Dairou, J. Cytochromes P450 of Caenorhabditis elegans: Implication in Biological Functions and Metabolism of Xenobiotics. Biomolecules 12 (2022).

63. Larsen, S. et al. Biomarkers of mitochondrial content in skeletal muscle of healthy young human subjects. J Physiol 590, 3349–3360 (2012).

64. Sowton, A.P. et al. Mtrr hypomorphic mutation alters liver morphology, metabolism and fuel storage in mice. Mol Genet Metab Rep 23, 100580 (2020).

65. Ding, N., Yuan, Z., Sun, L. & Yin, L. Dynamic and Static Regulation of Nicotinamide Adenine Dinucleotide Phosphate: Strategies, Challenges, and Future Directions in Metabolic Engineering. Molecules 29 (2024).

66. Chandel, N.S. NADPH-The Forgotten Reducing Equivalent. Cold Spring Harb Perspect Biol 13 (2021).

67. Tian, Y. et al. Mitochondrial Stress Induces Chromatin Reorganization to Promote Longevity andUPR(mt). Cell 165, 1197–1208 (2016).

68. Piwien Pilipuk, G., Galigniana, M.D. & Schwartz, J. Subnuclear localization of C/EBP beta is regulated by growth hormone and dependent on MAPK. J Biol Chem 278, 35668–35677 (2003).

69. Liu, X., Wu, B., Szary, J., Kofoed, E.M. & Schaufele, F. Functional sequestration of transcription factor activity by repetitive DNA. J Biol Chem 282, 20868–20876 (2007).

70. Seong, K.H., Li, D., Shimizu, H., Nakamura, R. & Ishii, S. Inheritance of stress-induced, ATF-2-dependent epigenetic change. Cell 145, 1049–1061 (2011).

71. Cabianca, D.S. et al. Active chromatin marks drive spatial sequestration of heterochromatin in C. elegans nuclei. Nature 569, 734–739 (2019).

72. Fellas, A. et al. Heterochromatin epimutations impose mitochondrial dysfunction to confer antifungal resistance. EMBO J (2025).

73. Ghose, R. et al. Mitochondria-derived nuclear ATP surge protects against confinement- induced proliferation defects. Nat Commun 16, 6613 (2025).

74. Merkwirth, C. et al. Two Conserved Histone Demethylases Regulate Mitochondrial Stress- Induced Longevity. Cell 165, 1209–1223 (2016).

75. Matilainen, O., Sleiman, M.S.B., Quiros, P.M., Garcia, S. & Auwerx, J. The chromatin remodeling factor ISW-1 integrates organismal responses against nuclear and mitochondrial stress. Nat Commun 8, 1818 (2017).

76. Kumar, T., Sharma, G.S. & Singh, L.R. Homocystinuria: Therapeutic approach. Clin Chim Acta 458, 55–62(2016).

77. Huemer, M. et al. Guidelines for diagnosis and management of the cobalamin-related remethylation disorders cblC, cblD, cblE, cblF, cblG, cblJ and MTHFR deficiency. J Inherit MetabDis 40,21–48(2017).

## Method and tables references

78. Padovani, F., Mairhormann, B., Falter-Braun, P., Lengefeld, J. & Schmoller, K.M. Segmentation, tracking and cell cycle analysis of live-cell imaging data with Cell-ACDC. BMC Biol 20, 174 (2022).

79. Maile, T.M. et al. Mass spectrometric quantification of histone post-translational modifications by a hybrid chemical labeling method. Mol Cell Proteomics 14, 1148–1158 (2015).

80. Lukauskas, S. et al. Decoding chromatin states by proteomic profiling of nucleosome readers. Nature 627, 671–679 (2024).

81. Robinson, M.D., McCarthy, D.J. & Smyth, G.K. edgeR: a Bioconductor package for differential expression analysis of digital gene expression data. Bioinformatics 26, 139–140 (2010).

82. Liao, Y., Smyth, G.K. & Shi, W. featureCounts: an efficient general purpose program for assigning sequence reads to genomic features. Bioinformatics 30, 923–930 (2014).

83. Love, M.I., Huber, W. & Anders, S. Moderated estimation of fold change and dispersion for RNA-seq data with DESeq2. Genome Biol 15, 550 (2014).

84. Al-Refaie, N. et al. Fasting shapes chromatin architecture through an mTOR/RNA Pol I axis. Nat Cell Biol 26, 1903–1917 (2024).

85. Timmons, L., Court, D.L. & Fire, A. Ingestion of bacterially expressed dsRNAs can produce specific and potent genetic interference in Caenorhabditis elegans. Gene 263,103–112 (2001).

86. Giovannetti, M. et al. SIN-3 transcriptional coregulator maintains mitochondrial homeostasis and polyamine flux. iScience 27, 109789 (2024).

87. Muschet, C. et al. Removing the bottlenecks of cell culture metabolomics: fast normalization procedure, correlation of metabolites to cell number, and impact of the cell harvesting method. Metabolomics 12, 151 (2016).

88. Artati, A., Couacault, P. & Witting, M. Nontargeted Metabolomics Using the Sciex ZenoTOF 7600. Methods Mol Biol 2925, 1–23 (2025).

89. Koopman, M. et al. A screening-based platform for the assessment of cellular respiration in Caenorhabditis elegans. NatProtoc 11, 1798–1816 (2016).

90. Tijsterman, M., May, R.C., Simmer, F., Okihara, K.L. & Plasterk, R.H. Genes required for systemic RNA interference in Caenorhabditis elegans. Curr Biol 14, 111–116 (2004).

91. Christine A. Doronio, H.L., Elizabeth J. Gleason, William G. Kelly A Chromodomain Mutation Identifies Separable Roles for C. elegans MRG-1 in Germline and Somatic Development. bioRxiv (2022).

